# Enhancing georeferenced biodiversity inventories: automated information extraction from literature records reveal the gaps

**DOI:** 10.1101/2020.01.16.908962

**Authors:** Bjørn Tore Kopperud, Scott Lidgard, Lee Hsiang Liow

## Abstract

We use natural language processing (NLP) to retrieve location data for cheilostome bryozoan species (text-mined occurrences [TMO]) in an automated procedure. We compare these results with data from the Ocean Biogeographic Information System (OBIS). Using OBIS and TMO data separately and in combination, we present latitudinal species richness curves using standard estimators (Chao2 and the Jackknife) and range-through approaches. Our combined OBIS and TMO species richness curves quantitatively document a bimodal global latitudinal diversity gradient for cheilostomes for the first time, with peaks in the temperate zones. 79% of the georeferenced species we retrieved from TMO (N = 1780) and OBIS (N = 2453) are non-overlapping and underestimate known species richness, even in combination. Despite clear indications that global location data compiled for cheilostomes should be improved with concerted effort, our study supports the view that latitudinal species richness patterns deviate from the canonical LDG. Moreover, combining online biodiversity databases with automated information retrieval from the published literature is a promising avenue for expanding taxon-location datasets.

## Introduction

Global biogeographical and macroecological studies require data on aggregate entities, such as location-specific occurrences of taxa and regional species assemblages, in order to understand emergent patterns at global and/or temporal scales (McGill, 2019). Assembly of such detailed yet broad-scale data is highly labor-intensive; the sampling effort required for a specific research question can be daunting for any one researcher or single research team. This is one reason why collaborative and often public databases have gained traction (Klein et al., 2019). For instance, empirical global biogeographic analyses (Costello et al., 2017; Rabosky et al., 2018) are increasingly based on public databases of georeferenced taxonomic occurrences, such as the Ocean Biogeographic Information System (OBIS, www.iobis.org) and the Global Biodiversity Information Facility (GBIF, www.gbif.org). Analyzing such georeferenced databases with tools that alleviate incomplete or biased sampling (Saeedi et al., 2019) allows us to address questions on large-scale distributions of clades, especially those that are well-represented in such databases. Yet for less well-studied clades, prospects for obtaining large amounts of such data are lower. Answering patternbased questions such as ‘how many species of clade z are found in location y’ and more process-oriented questions such as ‘how did the current latitudinal diversity gradient form’ both require location-specific taxonomic data in substantial volume. In addition, *generalized* biogeographic patterns and processes will be supported more robustly if they include a greater diversity of clades.

Cheilostome bryozoans, though less well-studied than several metazoan clades of similar size, are ubiquitous in benthic marine habitats. They are the most diverse order of Bryozoa with 83% of a conservatively estimated 5869 extant described species (Bock & Gordon, 2013). Bryozoans are ecologically important habitat builders (Wood, Rowden, Compton, Gordon, & Probert, 2013) and are vital components of the marine food chain (Lidgard, 2008). Despite important analyses of regional species distributions (Barnes & Griffiths, 2008; Clarke & Lidgard, 2000; López Gappa, 2000; Moyano, 1991), their global species richness distribution has never been quantified. We argue that even with concerns about the incompleteness of OBIS records for the purpose of inferring regional to global diversity patterns (e.g. Klein et al., 2019; Lindsay et al., 2017; Reimer et al., 2019), it is worth exploring cheilostome data in OBIS. We do so in order to identify spatial gaps in sampling but also to ask if automated information retrieval can enhance the species occurrence data available in OBIS.

Automated information retrieval (Hirschberg & Manning, 2015) is one recent approach to the time-consuming manual activity of data compilation from the scientific literature. Automated text-mining is well-established in the biomedical realm (Christopoulou, Tran, Sahu, Miwa, & Ananiadou, 2020; Percha, Garten, & Altman, 2012), but has only recently been adopted for biodiversity studies (Kopperud, Lidgard, & Liow, 2019; Peters, Husson, & Wilcots, 2017). As far as we are aware, automated text-mining has never been applied to the literature for extraction of taxon occurrences in given locations for the purpose to understanding biogeography (but see Page, 2019). We use natural language processing tools (Bojanowski, Grave, Joulin, & Mikolov, 2017; De Marneffe et al., 2014), to compile cheilostome text-mined occurrence data (TMO) to complement and potentially enhance data from OBIS.

Taxon occurrence data from OBIS and TMO are not expected to be the same. We ask if they could, separately or in combination, shed light on a long-standing biogeographic hypothesis in the bryozoological literature. Many different groups of organisms show the canonical latitudinal diversity gradient (LDG), a species richness peak in tropical regions and decreasing species richness towards the temperate and polar zones (Hillebrand, 2004; Menegotto, Kurtz, & Lana, 2019). Despite being common across marine and terrestrial realms, and among diverse eukaryote clades, the LDG is not universal (Chaudhary, Saeedi, & Costello, 2016). Extratropical bimodal species richness peaks have been observed, for example in deep-sea brittle stars (Woolley et al., 2016), razor shells (Saeedi, Dennis, & Costello, 2017) and foraminiferans (Rutherford, D’Hondt, & Prell, 1999). Bimodality has also been suggested for cheilostome bryozoans (Barnes & Griffiths, 2008; Clarke & Lidgard, 2000; Schopf, 1970).

The TMO and OBIS data in combination support the view that the latitudinal diversity pattern of living cheilostomes is bimodal. These data reveal highest levels of estimated species richness in temperate latitudes, but TMO species richness has a peak in the northern hemisphere while OBIS has a peak in the temperate south. Moreover, the data sets differ significantly in the geographic richness patterns in Atlantic versus Pacific ocean basins (Barnes & Griffiths, 2008; Schopf, 1970). We discuss the pros and cons of TMO and public databases such as OBIS and how their differences can help us understand the uncertainties of the retrieved spatial diversity patterns, beyond what is estimated within the confines of each dataset.

## Methods

### OBIS Data Retrieval

We use the R-package robis (Provoost & Bosch, 2020) to access OBIS (28.11.2019) and retrieve latitude/longitude occurrence records of cheilostomes. We remove records without species epithets. For taxonomic ambiguities such as cf., aff., we disregard the uncertainty; for instance, *Microporella* cf. *ciliata* becomes *Microporella ciliata.* Records with genus names that are not accepted according to either the Working List of Genera and Subgenera for the Treatise on Invertebrate Paleontology (pers comm. Dennis P. Gordon, 2019), World Register of Marine Species (WoRMS Editorial Board, 2020) or www.bryozoa.net (Bock, 2020) are also removed. The result is 561 unique genus names and 2453 unique genus-species combinations (henceforth simply species) in 144917 retained OBIS records.

### TMO (Text-Mined Occurrence) Data Retrieval

We follow a previously detailed text-mining procedure (Kopperud et al., 2019) with modifications. We extract text from two collections of published works, our own corpus (3233 pdf documents) and the GeoDeepDive archive (GDD, https://geodeepdive.org/), which contains full-text contents of journal articles. Only English language publications and those likely to feature extant bryozoans were used for information extraction (see Appendix S1 in Supporting Information).

We use CoreNLP (Manning et al., 2014) for an initial natural language analysis prior to information extraction, including tokenization, named-entity recognition, and dependency grammar annotation (Hirschberg & Manning, 2015). We use a pre-trained machine-learning model to recognize location names in the text (Finkel, Grenager, & Manning, 2005). To facilitate extraction of species, we compile names from the Working List of Genera and Subgenera for the Treatise on Invertebrate Paleontology (pers comm. Dennis P. Gordon, 2019), World Register of Marine Species (WoRMS Editorial Board, 2020) and www.bryozoa.net (Bock, 2020) that we then use in rule-based recognition (Chang & Manning, 2014). For example, consider a sentence from Tilbrook et al. (2001, p. 50):

> “The avicularia resemble those seen in *B. intermedia* (Hincks, 1881b), from Tasmania and New Zealand, but this species is only just over half the size of *B. cookae.”*

This sentence contains two species names *(“B. intermedia”* and *“B. cookae”)* and two location names (“Tasmania” and “New Zealand”). Each species-location pair is a candidate relation. The sentence implies that *B. intermedia* is found in New Zealand (a positive relation), but does not say anything about where *B. cookae* is found (a negative relation). We automate this distinction using a machinelearning classifier that we trained using a dataset of 4938 unique candidates labelled as positive or negative by two persons. Part of our procedure resolves the genus name referred to as *‘B.’* above (see Appendix S1).

We use a test data set comprising 10% of the labelled candidates to evaluate several aspects of our machine-classifier: (i) Accuracy, the ratio of correct predictions to all predictions; (ii) precision, the ratio of true positive predictions to all positive predictions; (iii) recall, the ratio of true positive predictions to all positive labels; (iv) false positive rate (FPR), the ratio of false positive predictions to all negative labels; and (v) F1, the harmonic mean of precision and recall. Each of these metrics yields different information on the reliability of the extracted data. We treat taxonomic ambiguities within TMO data in the same manner as OBIS records (see previous section).

### From TMO location names to spatial data

Location names (e.g., New Zealand, Tasmania) are submitted to the Google geocoding service (https://developers.google.com/maps/documentation/geocoding/) to acquire a bounding box with four latitude-longitude coordinates and a centroid (Fig. S1). We remove species occurrences in locations represented by bounding boxes that are larger than about 2% of the Earth’s surface using area calculations assuming a spherical globe. See Fig. S2 for how the bounding box sizes are distributed, and Fig. S3 for how alternative thresholds impact the result.

### Estimating latitudinal species richness

We initially evaluate species richness in thirty-six 5° latitudinal bands using two standard richness estimators that perform relatively well under a suite of conditions (Walther & Moore, 2005): Chao2 and Jackknife using the function specpool in the R package vegan (Oksanen et al., 2019). We treat these latitudinal bands as independent. We then repeat the procedure using thirty-six equal area bands, since areas represented within equal angle bands decrease poleward. To apply these estimators, we divide each (equal angle or area) latitudinal band into 5° longitudinal sampling units. We use the bias-corrected form of 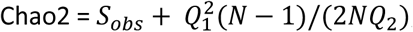, and incidence-based Jackknife = *S_obs_* + *Q*_1_(*N* — 1)/*N*. Here, *S_obs_* is the number of observed species in each band, *N* is the number of (longitudinal) sampling units, *Q*_1_ is the number of species observed in only one sampling unit, and *Q*_2_ is the number observed in two sampling units. Because terrestrial regions are not suitable habitats for marine cheilostomes, we mapped all landlocked longitudinal sampling bins (Fig. S4) based on a 1:10 m map of global coastlines (Patterson, 2019). We removed the landlocked bins prior to richness estimation. For OBIS data where spatial coordinates are points, it is trivial to assign data to sampling units. For TMO, we assume that a species occurs in all of the sampling units that intersect the bounding box associated with the location. TMO bounding boxes vary in size, but most are smaller in area than our sampling units (Fig. S2).

In addition to Chao2 and Jackknife estimators, we also determined range-through species richness. Here, we assume that a species spans its southernmost and northernmost occurrence record, regardless of whether it is observed in any intermediate latitudinal band.

The code and data required to reproduce the analyses and the figures are stored datadryad.org and will be available upon submission.

## Results

### Capturing species diversity: comparing OBIS and TMO

Applying the text-mining procedure to our corpora, we retrieved 1780 species in 382 genera, and 1915 unique location names among 9653 TMO records. Only 27% of the species in the OBIS data that we retained were also in TMO. Similarly, only 41% of these combinations in the TMO occurred in OBIS. 20% of the species richness is common to both (Figs. 1, S5). In combination with OBIS data, we have species-location information from 3323 species or 68% of 4921 described cheilostome species (Bock & Gordon, 2013).

**Fig. 1.**
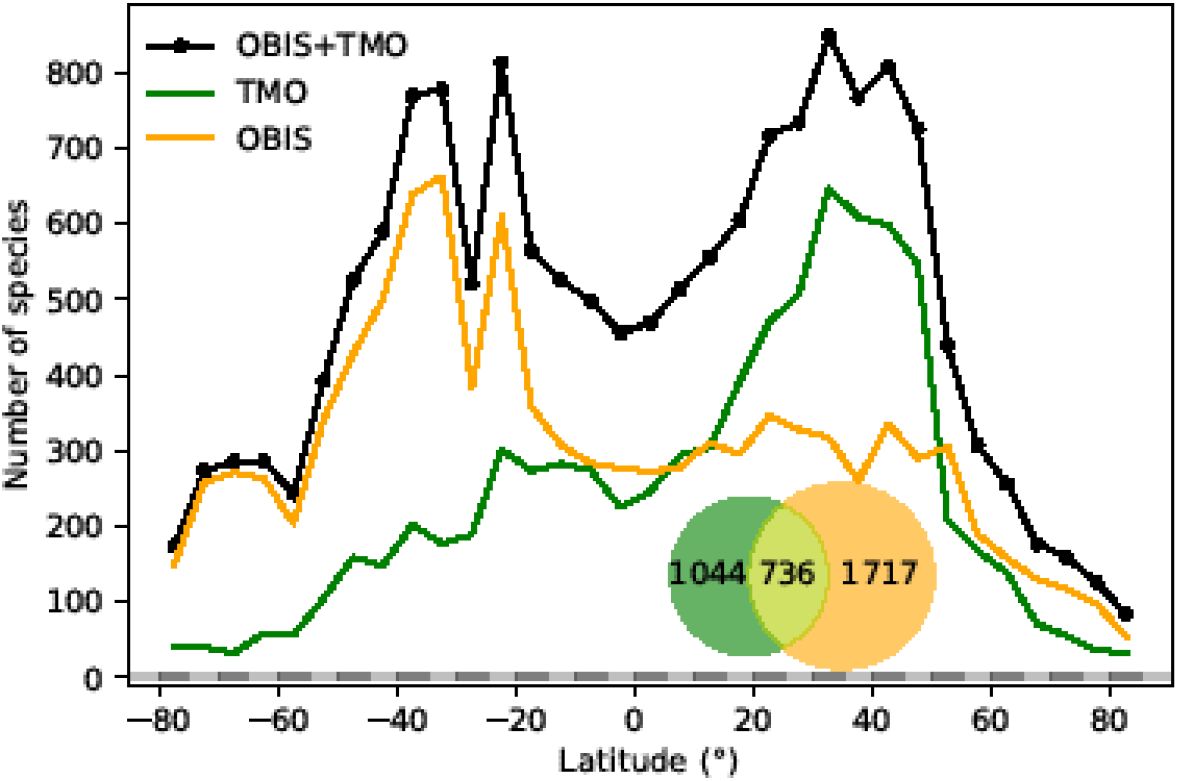
Global range-through latitudinal species richness for cheilostome bryozoans. The black line shows combined Ocean Biogeography Information System (OBIS) and text-mined occurrence (TMO) richness, and orange and green curves show range-through richness for OBIS and TMO separately. The inset is a Venn-diagram showing the global overlap in species between OBIS and TMO.

Our machine-classifier achieved an accuracy of 73.1%, F1 of 76.8%, recall of 78.9%, FPR of 34.3% and precision of 74.8% as estimated with our test set (Fig. S6b). These results are substantially better than a random classifier baseline, but not as good as the human annotator repeatability. Specifically, the FPR among annotators is about 15% (n = 200). A random classifier that is as unbalanced as our training data (60% positive labels) would yield 60% false positives, but a random classifier equaling our classifier’s recall of 78.9% would have the same false positive rate of 78.9% (see Appendix S1).

### Latitudinal species richness patterns

Combined TMO and OBIS data in plots of range-through species richness show a bimodal pattern with species richness peaks in both hemispheres surrounding 40° and −40° (Fig. 1). Inferred species richness in both of these peaks is about double that in the tropics. The two data sources contribute different latitudinal constituents, as suggested by the limited overlap in their species composition (Fig. 1 inset).

Chao2 and Jackknife estimated species richness from OBIS shows two peaks between −20° and −45° that are more than double the next highest peak between 25° and 50° (Fig. 2a). In contrast, TMO estimated richness shows a highest peak between 30° and 45° (Fig. 2b). With minor exceptions in the Antarctic where spatial distortion is largest, equiangular and equi-areal bands yield nearly identical inferences (compare Fig. 2a,c). The latitudinal pattern appears smoother when using larger latitudinal band sizes (Fig. S7), while retaining a qualitatively similar picture. Longitudinal sampling bins of varied sizes appear not to be important for the Jackknife and Chao2 estimators (Fig. S8).

**Fig. 2.**
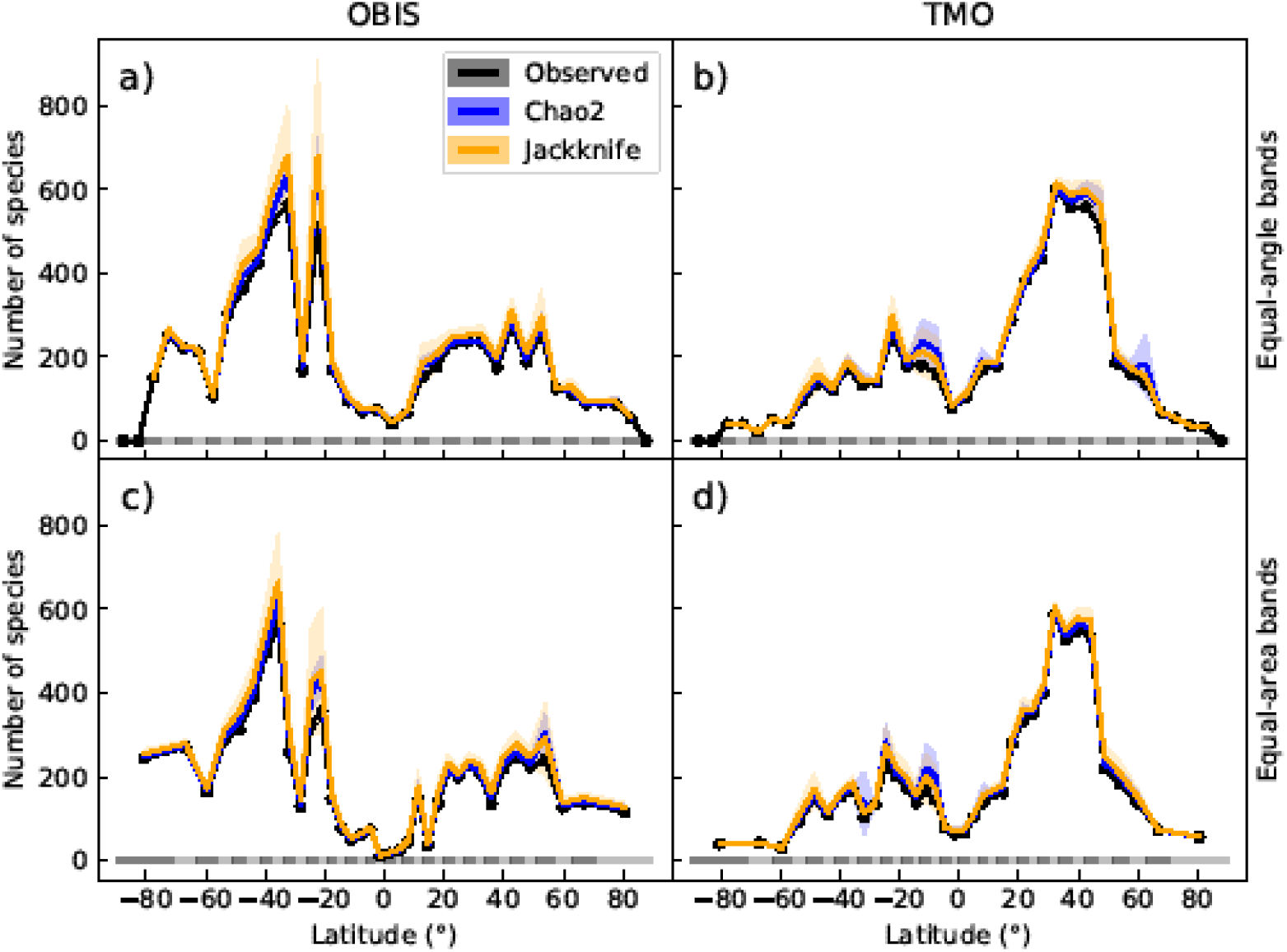
Global latitudinal species richness for cheilostome bryozoans, estimated using Chao2 and Jackknife. The top panels show richness for Ocean Biogeography Information System (OBIS) and text-mined occurrences (TMO) data in 5° equal-angle latitudinal bands. The lower panels show the equivalent in 5° equal-area latitudinal bands. Black lines show the observed richness, while blue and orange lines show the Chao2 and Jackknife estimates, respectively. The shaded areas are 95% confidence intervals. See Figs. S7 and S8 for alternative band and bin sizes.

The northern hemisphere peak in richness (Fig. 1) reflects TMO records from the Mediterranean and Japan, but also from the Atlantic Ocean (Fig. 3a,e), including the British Isles. Note that we did not include the Mediterranean as part of the Atlantic basin for Fig. 3. A portion of the TMO data are spatially imprecise, for example the location names “France”, “Spain” or “Morocco” may be associated with Mediterranean endemics, yet these records could contribute to the Atlantic richness counts in Fig. 3. The spatially precise OBIS data show a much lower peak in the Eastern Atlantic (Fig. 3e, orange line shifted slightly northward), reflecting data from the British Isles and northern Europe. Conversely, OBIS data mainly from Australia and New Zealand contribute disproportionately to the huge southern hemisphere peak. The richness captured by OBIS in Australia and New Zealand is not reflected by TMO species richness (Fig. 3b,d). The western Atlantic and eastern Pacific do not display such pronounced temperate zone peaks (Fig. 3c,f). Looking at individual ocean basins, TMO and OBIS are sometimes congruent and other times incongruent. For example, there is an absence of OBIS records in Japanese waters, and there are similarly few TMO and OBIS records in the Indian Ocean (Fig. 4).

**Fig. 3.**
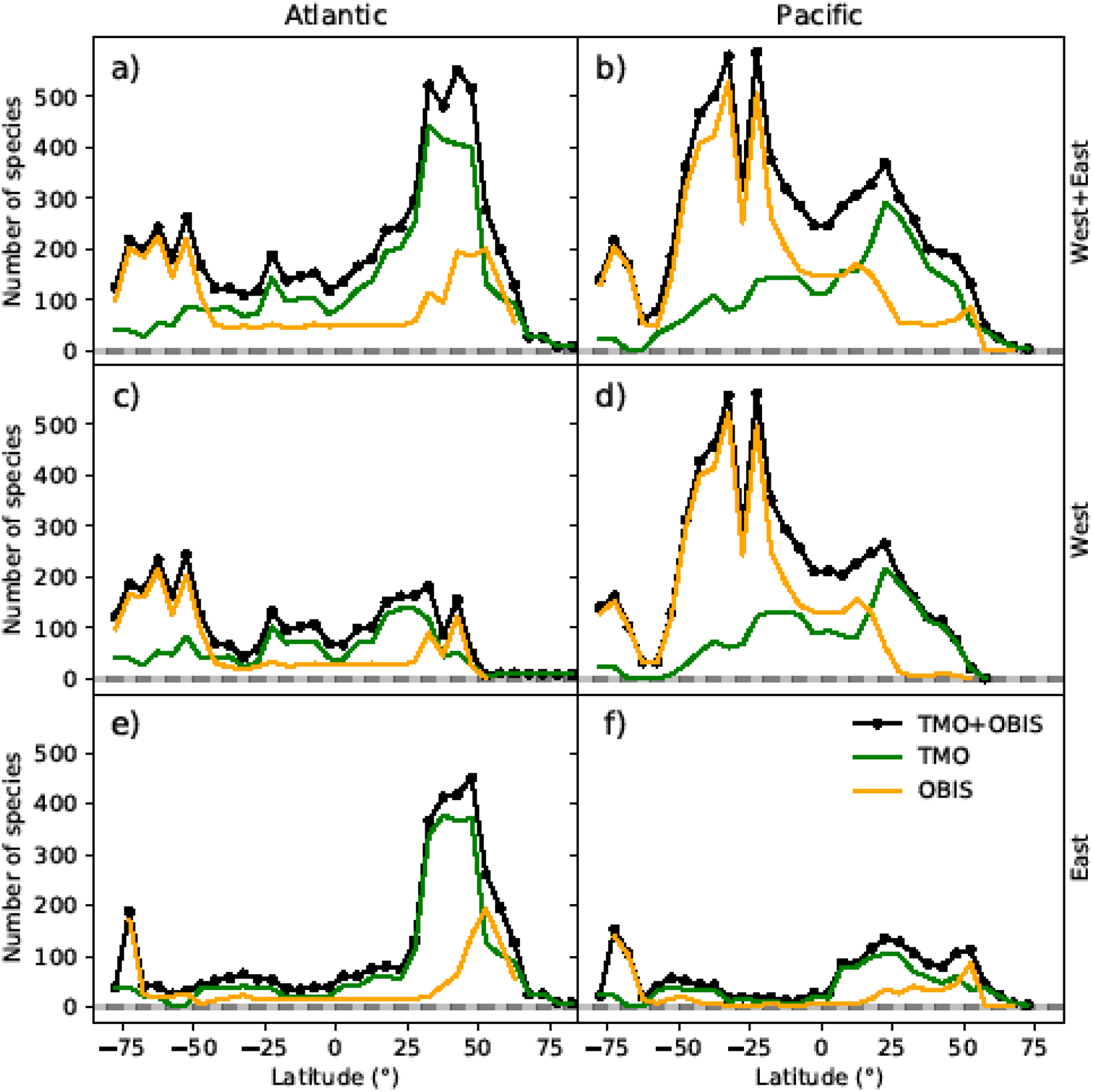
Range-through latitudinal species richness for cheilostome bryozoans in the Atlantic and Pacific Oceans. The left column shows species richness in the Atlantic, and the right column shows that in the Pacific. The panel rows represent the eastern, western or the entire ocean basins. Orange and green lines represent Ocean Biogeography Information System (OBIS) and text-mined occurrences (TMO), respectively, and black lines are the joint data. Note that in this figure, the Atlantic borders Greenland and Iceland in the north, and the Antarctic in the south, but does not include the Gulf of Mexico, the Caribbean, the Baltic Sea or the Mediterranean. The Pacific borders the Bering Strait in the north, and includes the South China Sea, the Java Sea, north and east Australia, Tasmania as well as the Antarctic border.

Such varied regional species richness patterns are in part influenced by the geographic occurrence of samples. Figure 4 summarizes the relative distribution of species-location records for TMO and OBIS data as global heatmaps. For OBIS data, there are about one order of magnitude fewer records in tropical regions than for subtropical and temperate ones (Fig. S9a). While there are also fewer TMO records in tropical regions, the effect is not as pronounced (Fig. S9b). Northern and southern hemisphere species richness peaks in the two data sets (Fig. 1) correspond with high regional densities of TMO and OBIS records, respectively (Fig. 3e,d).

**Fig. 4.**
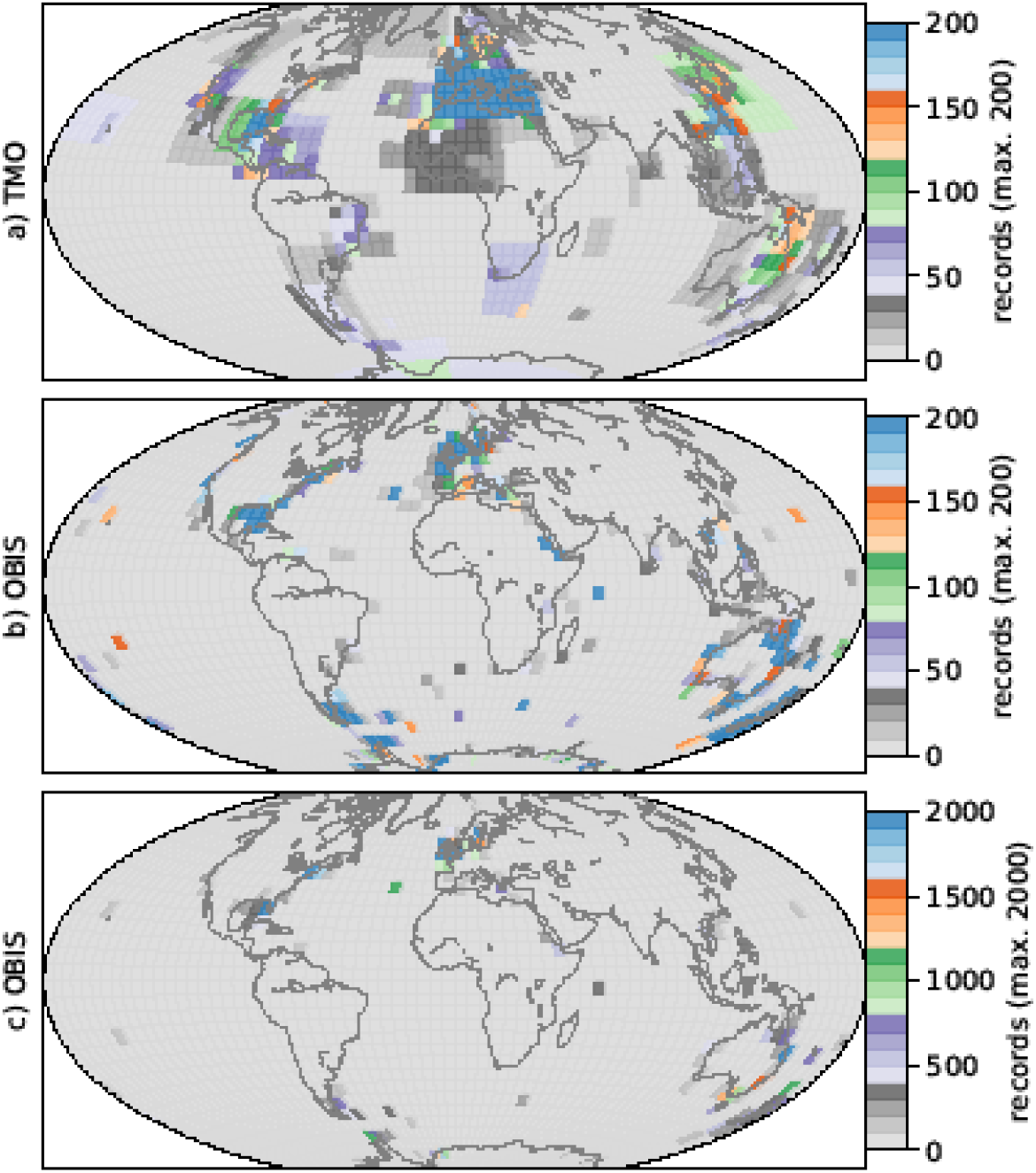
Heatmaps for cheilostome bryozoan occurrence records per 5° latitude by 5° longitude bins. The color axes are truncated for visualization purposes, to a maximum of 200, 200 and 2000 in a), b), c), respectively. There are about 750 maximum records per bin in the Mediterranean for the text-mined occurrences (TMO), and about 35000 maximum records in the British Isles for the Ocean Biogeography Information System (OBIS) data. The globe is plotted using the Hammer equal-area projection. See Fig. S11 for the same figure plotted using the plate carrée projection.

## Discussion

Causal hypotheses for a LDG and contrarian patterns are plentiful and can sometimes be tested in groups with rich and relatively unbiased spatial data from both extant and extinct taxa (Jablonski et al., 2013; Jablonski, Roy, & Valentine, 2006; Krug, Jablonski, & Valentine, 2007) or those with independent molecular phylogenetic evidence (e.g. Rabosky *et al.* 2018). We believe ours is the first study to quantify global cheilostome species biogeographic patterns. Using a combined TMO and OBIS perspective, and a bimodal latitudinal diversity gradient in cheilostome species richness is quite apparent. Yet, at present, we can merely speculate about what processes that may have led to their latitudinal pattern. Given the biases and heterogeneity of the data we explored which are striking when comparing our two data sources, we also need to consider (i) how this pattern coincides with previous observations, and (ii) methodological, sampling, and taxonomic concerns.

Two patterns in our analyses are similar to Schopf’s (1970) findings from then-scarce available data: higher species richness on the eastern margin of the Atlantic and the western margin of the Pacific compared to their opposite margins, and increasing richness with latitude away from the equator. Our combined data conforms with the first finding, but still doesn’t capture the richness of the severely-understudied Philippine-Indonesian region and its many archipelagoes (Gordon, 1999; Okada & Mawatari, 1958; Tilbrook & De Grave, 2005). Changes to the second finding are more nuanced, and may partly reflect relatively lower equatorial sampling density (Chaudhary et al., 2016; Chaudhary, Saeedi, & Costello, 2017; Fernandez & Marques, 2017; Menegotto & Rangel, 2018) apparent in both of our datasets (Fig. S9). However, our observed peaks of species richness are at significantly higher latitudes than those reported for bryozoans in Chaudhary et al. (2016).

Fossil and modern patterns of bryozoan abundance in cool-water carbonate sediments suggest that the lower tropical species richness is not merely a sampling artifact. Modern bryozoan-dominated carbonate platforms are far more common on cool-water temperate shelves than on tropical ones (James & Clarke, 1997; Schlanger & Konishi, 1975). Cenozoic tropical bryozoan faunas are both less abundantly preserved and less diverse than those from temperate latitudes, possibly reflecting biotic interactions, preservational biases, and cryptic existence in shallower-water habitats dominated by corals, calcareous algae, and other photobiont organisms (Taylor & Di Martino, 2014; Winston, 1986). A far-reaching study by Taylor & Allison (1998) showed that 94% of bryozoan-rich post-Paleozoic sedimentary deposits formed outside of the paleotropics, which may be especially significant if regional species richness and skeletal abundance are linked. About a third of all described bryozoan species occur south of −30°, and 87% of these are cheilostomes (Barnes and Griffiths, 2008).

We chose to discretize the data in latitudinal bands and longitudinal bins that are larger than those previously used e.g., in Rabosky et al. (2018). The choice of band- and bin sizes for species richness estimation is somewhat arbitrary. Differing choices suggest quantitatively dissimilar inferences, although the bimodality is still apparent in the cases we have explored (Figs. S7 and S8). A range-through latitudinal diversity approach (Fig. 3) assumes that any species that is not observed in a gap between two adjacent latitudinal bands should contribute to species richness in that gap, but this assumption is quite easily broken (Menegotto & Rangel, 2018). The bounding boxes used for TMO locations may also tend to bleed range margins to as opposed to OBIS point location data. Richness estimates may be inflated via range-through estimates, particularly in the tropics, compared to estimating richness independently in each latitudinal band which yields lower estimates (Fig. 2). Regardless, both methods for estimating species richness give a picture of bimodality.

Global biogeographic studies such as ours are more prone to the issues of sampling and taxonomic concerns than local or regional ones, simply due to their scope. Large sampling gaps are apparent in both TMO and OBIS datasets. The development and application of richness estimation models that distinguish true absences from non-observations (Iknayan, Tingley, Furnas, & Beissinger, 2014) may help improve inferences, but are likely insufficient to overcome blatant sampling gaps. Overall, there are relatively few records in the Indian Ocean, most of the South Atlantic, and eastern margin of the Pacific. TMO records for the Arctic are sparse, as are OBIS records for the northwest Pacific. Aside from a few extreme outliers from OBIS British Isles locations, species richness and number of records per 5° latitudinal band have a strong positive relationship (Fig. S10). Independent taxonomic surveys of underrepresented regions in one or both datasets corroborate the existence of significant gaps (Florence, Hayward, & Gibbons, 2007; Grischenko, Mawatari, & Taylor, 2000; Hirose, 2017; X. Liu & Liu, 2008; López Gappa, 2000; Moyano, 1991; Vieira, Migotto, & Winston, 2008). The OBIS records may partly reflect recent histories of active bryozoan research programs in the Antarctic (Barnes & Griffiths, 2008; Figuerola, Barnes, Brickle, & Brewin, 2017) and Australia and New Zealand (Wood et al., 2013) as well as contributions to OBIS that differ substantially among research institutions. On the other hand, TMO extracted extensive species-location information from the Mediterranean (27° to 50°) that are severely wanting in OBIS, demonstrating that combining disparate data sources can help bridge gaps in global biogeographic studies.

Taxonomic errors inevitably exist in large databases. Taxonomy is continuously subject to revisions (Bock & Gordon, 2013), not all of which are accounted for in our datasets. Many species await description; (Gordon, Bock, Souto-Derungs, & Reverter Gil, 2019) suggest that there are over 6,400 ‘known’ cheilostome species without commenting on nomenclatural status, suggesting that there are up to 600 ‘known’ species that need naming. Our machine-classifier is currently unable to extract location information for 18% of the species that were detected in our corpus of published works (Fig. S5). Our conversion of taxonomic ambiguities into certainties likely deflated species richness estimates, while mistaken inclusion of fossil species names may have inflated richness estimates. We have assumed these do not necessarily introduce spatial bias. Additionally, many bryozoan species determined by traditional morphological methods may actually consist of unrecognized species complexes (Fehlauer-Ale et al., 2014; Jackson & Cheetham, 1990; Lidgard & Buckley, 1994). While the portion of TMO data that is derived from the taxonomic literature may be less plagued by taxonomic misidentifications, the same cannot be easily argued for faunal lists or ecological surveys, which may also be part of OBIS data. However, in our experience, broad inferences based on synoptic, large-scaled databases tend to change significantly with different models, more so than data updates (Liow, Reitan, & Harnik, 2015; Sepkoski, 1993).

In terms of our text-mining task, we found that generating and classifying species-location candidates here is more challenging than classifying species-age candidates (Kopperud et al., 2019). An F1 result of about 77.5% is not uncommon for relation extraction studies (Henry, Buchan, Filannino, Stubbs, & Uzuner, 2020; Kim, Kim, & Lee, 2019), especially for datasets with low label assignment repeatability. Nonetheless, while the accuracy of the machine-classifier is less sensitive than human evaluation, its FPR is substantially lower than a null model. Note that the classifier merely provides a probabilistic measure of whether the sentence provides evidence that a species is present at a geographic location. In the event of a false positive, it is still possible that the species is actually present in that particular location. On the other hand, there is a wealth of species mentions for which we were not able retrieve any species-location candidates (Fig. S5). It is possible to extend our approach by considering cross-sentence candidates (Gupta, Rajaram, Schütze, & Runkler, 2019), although these methods are usually less accurate. Alternatively, we could go beyond standard NLP tools, which are relatively flexible and easy to adopt, and use non-linguistic features (such as tables and spatial layout) for information extraction, as has been suggested in the knowledge base creation literature (Schlichtkrull et al., 2018). However, methods for information extraction that combine linguistic and non-linguistic features are still at an early stage of development.

The main advantage of automatic information retrieval over collaborative data-entry is that of time and resource investment. The information retrieval procedure is largely independent of the size of the literature, or the taxonomic scope, say for cheilostomes versus all metazoans. Public biodiversity inventories such as GBIF and OBIS require large consortia and networks of research factions to contribute their data. Conversely, there is a wealth of biodiversity knowledge available in the published literature, and it is feasible for one person or a small team to extract substantial amounts of data quickly using automated information retrieval. We have used some supervised classification methods, which require us to generate training data. However as NLP is adopted in the biodiversity literature, it will become easier to use distantly supervised relation extraction (Hirschberg & Manning, 2015).

Biodiversity inventories such as OBIS are vital for supplying data for inferences of global biogeographic patterns. While we strongly support the continued development of these databases, we demonstrated that our automated information retrieval approach can enhance such inventories when answering global-scale questions, especially for under-studied taxa. To understand how the spatial diversity of cheilostomes has come to be will require continued and concerted efforts in taxonomic investigations (Bock & Gordon, 2013), compilation of more spatial data especially in areas currently devoid of deposited information (Klein et al., 2019), tool-development in automated data retrieval (Kopperud et al., 2019), and continued research in molecular phylogenetics (Orr et al., 2019).

## Supporting information

Appendix S3 - TMO references

## Acknowledgements

We thank Mali H. Ramfjell for compiling part of our training dataset, the GeoDeepDive group, especially Ian Ross and Shanan Peters, for providing access to articles, and OBIS and their contributors for their georeferenced taxonomic data. We thank Phil Bock for maintaining bryozoa.net, Dennis Gordon for an updated version of the Working List of Genera and Subgenera for the Treatise on Invertebrate Paleontology, and both for their contributions to WoRMS. This project has received funding from the European Research Council (ERC) under the European Union’s Horizon 2020 research and innovation programme (grant agreement No 724324 to L.H. Liow).

## Supporting Information

**Appendix S1:** Extended methods.

**Appendix S2:** Supplementary figures.

**Appendix S3:** Bibliographic references for TMO data.

**Code and data supplement**: Will be available on datadryad.org upon submission.

