## Appendix S1 - Extended methods for "Enhancing georeferenced biodiversity inventories: automated information extraction from literature records reveal the gaps"

**Supporting information to:** Enhancing georeferenced biodiversity inventories: automated information extraction from literature records reveal the gaps

**OBIS data filtering**

We used the R-package robis (Provoost & Bosch, 2020) to download OBIS data for cheilostome bryozoans with the function call ‘robis::occurrence(scientificname = "Cheilostomatida")’. We considered only records with Linnéan binomial names, i.e., containing a genus name and a species name. See the main text for how we treated taxonomic uncertainties. We did not vet the OBIS data (nor the TMO data described in the next section) for errors using manual case-by-case inspection.

**Automated retrieval of location-species data**

Our text-mined occurrences (TMO) are retrieved in a series of steps as already summarized in the main text. The steps are as follows. (i) We acquire the relevant literature. (ii) We annotate the text with linguistic features, such as recognizing genus or species names (henceforth taxon or taxa) and toponyms (i.e. place names) as entities. (iii) We extract all candidates: pairs of taxon names and toponyms that co-occur in a sentence. (iv) We build a machine-learning classifier that verifies that the candidates have correct toponyms. (v) We develop a machine-learning model that classifies each candidate according to whether or not the sentence says that the taxon is found in the location described by the toponym. (vi) We use a geocoding service to pin the toponyms to a location.

(i) Acquisition of literature

For the TMO, we used two different corpora (i.e., collections) of published works. In an earlier publication (Kopperud, Lidgard, & Liow, 2019), we used a corpus that was primarily focused on paleontological literature. We supplemented that corpus with publications that were likely to contain descriptions of extant cheilostomes, such as ecological surveys or taxonomic descriptions of new or revised taxa. This first corpus (our own collection as indicated in the main text) has 3223 published documents with associated pdfs. Of these, we filtered those that were not in English using a language detection tool (Nakatani, 2010), retaining about 80% of the extracted publications. The second corpus we used was extracted from the GeoDeepDive or GDD archive ([www.geodeepdive.org](http://www.geodeepdive.org)). GDD is a (U.S.) National Science Foundation funded project that gathers full-text contents of journal articles to facilitate data-mining. We searched all documents available on GDD, in Mar 2019 (when more than 10 million documents were available), and selected publications containing any cheilostome genus name found in [www.bryozoa.net](http://www.bryozoa.net) (Bock, 2020) and/or World Register of Marine Species (WoRMS Editorial Board, 2020). From these selected documents, we extract sentences that contain potential candidate pairs (see later section), that we then filter to remove non-English language ones. Intellectual property rights restrict GDD from redistributing documents in their entirety. Thus, while we could run our text-mining procedure on GDD and retrieve a summary of candidates (explained later), we did not have access to ‘read’ the main text. For this reason, we filtered non-English literature using the same language detection tool (Nakatani, 2010) but on the candidate level, instead of the document level.

Since our corpora consist of both paleontological and recent ecological or taxonomic literature, we sought to eliminate documents of the former. We used a machine-learning classifier as, first, there are too many articles to label, and second, since we did not have access to ‘read’ the documents in GDD as explained above. We manually annotated 839 publications being primarily about fossils, and 825 primarily about recent faunas. Using these annotations, we trained a supervised text classifier to predict if the document in question is either more about fossil or recent faunas. Specifically, we trained a support vector machine (SVM), implemented in scikit-learn (Pedregosa *et al.*, 2011). We used a radial-basis kernel function (allowing for a non-linear decision boundary) for the SVM, but otherwise used default settings. To enable this classifier, we used the number of mentions of geologic time intervals (e.g. “Upper Eocene”) and stratigraphic names (e.g. “Needmore Shale”, or “Lincolnshire Limestone”) per document as features (see the next section for how these are recognized). We evaluated the classifier using 5-fold cross validation and achieved an accuracy of 83.4 ± 1.8% (standard deviation). To counteract possible effects of class imbalance during cross validation, we weighed the inputs such that they contribute inversely proportional to their class frequency during training. We then applied the classifier to any documents that were not yet labeled, and after filtering we retained 58% of publications pertaining mainly to recent faunas.

(ii) Linguistic annotation

We used CoreNLP (Manning et al., 2014) for an initial natural language analysis, including tokenization, named-entity recognition, and dependency grammar annotation. Tokenization splits text into tokens, which are words and punctuation, after which sentences can be demarcated. In order to recognize bryozoan species names, we first assembled a list of known species names, based on the online compendium [www.bryozoa.net](http://www.bryozoa.net) and WoRMS. In order to recognize and count mentions of geologic time intervals and stratigraphic names, we used known names from GeoWhen (Rohde, 2005) and Macrostrat (S. E. Peters, Husson, & Czaplewski, 2018). We used these names to assemble a set of rules (TokensRegex, Chang & Manning, 2014) that recognize relevant names of species, stratigraphic units and geological time intervals. Finally, we applied CoreNLP to assign dependency grammar (i.e. relations among words in a sentence, De Marneffe *et al.*, 2014), which we used as a feature for the relation classifier explained later.

We used a generic, pre-trained machine-learning model (Finkel, Grenager, & Manning, 2005) to recognize toponyms (i.e. place names or locations) in text. We found that this generic named-entity recognition was prone to false positives such as author names (e.g. “Hincks” or “Darwin”). In order to remedy this, we trained a machine-learning classifier to verify whether the tagged entities were indeed proper toponyms (see section “Toponym verification”).

(iii) Candidate extraction

This was largely detailed in the main text, with the sentence below as an illustration (Tilbrook *et al.* 2001, page 50):

“The avicularia resemble those seen in *B. intermedia* (Hincks, 1881b), from Tasmania and New Zealand, but this species is only just over half the size of *B. cookae*.”

Whenever we found an abbreviated genus name, such as “*B.*”, we searched for genus names in the current and 14 previous sentences. In reverse chronological order, we looked for any span (i.e. one or more consecutive tokens) tagged as being a taxon that starts with the same capital letter, and chose the first genus name for the de-abbreviation (here “*Beania*”).

(iv) Toponym verification

As mentioned in section (ii) the generic named-entity recognition was prone to false positives. To reduce false positives here, ­­­we evaluated 6000 candidates and annotated whether the assigned toponym was indeed a toponym or a false positive. We used these annotations to train a machine-learning classifier; a neural network implemented in Keras (Chollet et al., 2019). For the previously mentioned classifiers (see sections i,ii), we used reference implementations and did not change the default settings considerably. For the toponym classifier, however, we made several decisions when engineering the classifier. For this reason, we detail the toponym classifier more closely than the other machine-learning models we have used.

A neural network can be thought of as a computational graph, with inputs, outputs, and several operations in the middle. For organizational purposes, we refer to sections of these operations as “layers”. Each layer has a set of associated coefficients or weights that are either set a priori or initialized randomly. For a set of inputs and labels, it is possible to evaluate a loss function that estimates how much the output diverges from the assigned label. Next, one can employ the back-propagation technique (Rumelhart, Hinton, & Williams, 1986) to compute partial derivatives or gradients for the loss value with respect to each coefficient in the network. These gradients, coupled with an algorithm for gradient descent, can be used to learn the coefficients, and consequently minimize the loss of the network during training. We split the annotation data in three parts: training-, validation- and test sets (roughly 80, 10, 10% candidates per set). We used the training data to learn the coefficients of the network. We used the validation data to fit the hyperparameters (e.g. learning rate, layer dimensions) and to decide when to stop training. By evaluating the trained network on the test data we get an estimate of how well the classifier performs on out-of-sample sentences. In order to remove possible confounding effects of similar sentences in a publication, we made sure that all candidates from a single publication were contained within only one of the training, validation or test datasets.

Our toponym classifier received two inputs: the sequence of tokens in the candidate’s sentence, and a sequence of indicator variables denoting where the toponym is in the sentence. These were fed to embedding layers: the tokens were embedded in a 300-dimensional vector space using a pre-trained fastText model (Bojanowski, Grave, Joulin, & Mikolov, 2017), and the indicator variables were embedded as two orthogonal unit-vectors. The coefficients for the embedding layers were constrained to be static during training. After concatenation, we fed the embedded values to a Long Short-Term Memory (LSTM, Hochreiter & Schmidhuber, 1997) recurrent layer, with dropout values of 0.2. Since the sentences were of variable length, we padded the shorter sentences and used masking to avoid any processing of the padded features during training. We used the final time step of the second recurrent layer as input to a final two-dimensional hidden layer, with a softmax activation function. Since the softmax function maps all real inputs to outputs of [0,1], while ensuring that they sum to one, we interpret the output as a probability mass. We used cross entropy as the loss function, and the ADAM algorithm (Kingma & Ba, 2014) for gradient descent. Unless otherwise stated, we used default settings in Keras for other parameters and options.

For each epoch (an iteration in which the classifier sees the entire training set), we computed the F1 metric (the harmonic mean of precision and recall). We trained the classifier for 50 epochs, and saved the coefficients whenever the validation F1 was better than the previously best validation F1, effectively conditioning the learning on maximizing the F1 metric, in line with most relation extraction studies (Christopoulou, Miwa, & Ananiadou, 2019). The toponym classifier achieved an F1 of 94.7%, accuracy of 93.2%, recall of 97.4%, precision of 92.2%, and a false positive rate of 14.0% (see Fig. S6a) when evaluated on the test set.

(v) Relation classifier

We manually annotated 4938 unique candidates (species-toponym pairs) to form our training dataset. If the sentence explicitly stated or strongly implied that the taxon was found in the mentioned location, we labelled the candidate as ‘positive’. If not, we labelled the candidate as ‘negative’. These annotations were made by two persons, with intra- and inter-annotator accuracies of 91% (n = 200) and 85.8% (n = 211), respectively. This classifier is analogous to the toponym classifier (see previous section), except we use neither the indicator variables nor the entire sentence. Instead, we use the sequence of tokens along shortest path in dependency grammar between the two spans in the sentence (see Xu *et al.*, 2015; Kopperud *et al.*, 2019). We trained our machine-classifier using the training set of these candidates to evaluate the classifier. This relation classifier achieved an F1 of 76.8%, accuracy of 73.1%, precision of 74.8%, recall of 78.9% and false positive rate of 34.3% (see Fig. S6b).

We discussed in the main text how our estimated false positive rate of 34.3% is better than a random classifier baseline, but not as good as the false positive rates between human annotators at 14% and 16% (n = 200, assuming annotator A is correct, then evaluating annotator B, and vice versa). Similarly, the classifier accuracy of 73.1% is better than a weighted coin random classifier, which if we assume is as unbalanced as the labelled candidates (60% positives), gives an accuracy baseline of 52% (0.6^2^ + 0.4^2^ = 0.52). Yet, this is not as good as the intra- and inter-annotator accuracies at 91% (n = 200) and 85.6% (n = 211), respectively. Incidentally, applying the relation classifier from (Kopperud et al., 2019) trained on relating taxon spans and geologic time interval spans, yielded virtually the same performance (evaluated on the same test data), despite not having been trained on the specific task or having ‘seen’ any toponym spans.

The relation classifier could potentially be improved by increasing the amount of training data, and/or using a more complex classifier, such one that takes into account context-dependent word embeddings (Devlin, Chang, Lee, & Toutanova, 2018; M. E. Peters et al., 2018). However, we believe that the major bottleneck is not lack of natural language understanding. Rather, the candidates themselves are not always linguistically sound, coherent and self-contained sentences. Specifically, much of the information that relays the relation between the two spans in the taxonomic literature of interest is coded in titles, sub-titles, variation in font type, font size, and spatial layout of the paragraphs. This also introduces errors for the sentence splitting procedure in CoreNLP. The features we used to capture the relation (Part-of-Speech and Dependency grammar) are inherently limited since they are designed to work on relatively coherent, self-contained and complete sentences. Standard Natural Language Processing tools are flexible and relatively easy to adopt, but for this particular problem it could be advantageous to use other, non-linguistic features to facilitate the information extraction, as has been suggested in the knowledge base creation literature (e.g. Schlichtkrull et al., 2018). At present, methods for information extraction using non-linguistic features are still at an early stage of development.

(vi) Geocoding: converting toponyms to coordinates

As stated in the main text, we used the Google geocoding service (https://developers.google.com/maps/documentation/geocoding/) to acquire a bounding box with four latitude-longitude coordinates and a centroid (Fig. S1). For any query, the geocoding service returns one record that best matches the toponym. Some toponyms are inherently more precise than others (e.g., “Suez Canal” vs “Red Sea”), and the service also provides a description of how precise it infers the record to be. These description codes are “ROOFTOP”, “RANGE_INTERPOLATED”, “GEOMETRIC_CENTER”, and “APPROXIMATE”, ranging from very fine to coarse spatial resolution. Our toponyms are in general not as precise as the most precise locations returned by the geocoding service. For our data, such precise matches are usually incorrect mappings of a toponym to a location (e.g. “Norfolk Ridge”, a marine ridge between New Zealand and New Caledonia is mapped to a street address in Norfolk, USA). To avoid such problems, we only used the categories “GEOMETRIC_CENTER” and “APPROXIMATE” for the geocoder results, retaining 88.3% of the unique locations. Despite considering only these coarse categories, the spatial resolutions of the toponyms are highly variable. To get an estimate of the spatial precision of the records, we calculated and used the area of the bounding box as a proxy for spatial precision. We removed mappings that did not have an associated bounding box (24.3%). We assumed that the shape of the Earth can reasonably be approximated by that of a sphere, and calculated the area of the bounding box as follows:

$A= 2\pi r^{2}(\sin\left( \alpha_{north} \right)-\sin\left( \alpha_{south} \right))(\beta_{east}-\beta_{west})/360^{\circ}$,

where *r* is the radius of the earth, *α* are the latitudes and *β* the longitudes of the bounding edges. The area calculation assumes that the planes demarcating the northern and southern edges are perpendicular to the axis of rotation, and that the planes demarcating the western and eastern edges are coincident with the axis of rotation. The bounding box area is always larger than the actual area of the location described by the toponym. For example, the “Mediterranean” or “Mediterranean Sea” is in reality about 2 500 000 km^2^ (Salah & Boxer, 2019) but our bounding box estimate is about 6 350 000 km^2^. We used these area estimates to remove species occurrence records in locations represented by bounding boxes that were larger than a certain threshold. For the analyses presented in the main text, the threshold was 1.86% of the Earth’s surface (see Figs. S2, S3).

**Regional maps**

In order to separate richness estimations for the different ocean basins (Fig. 3), we drew polygons (using [www.geojson.io](http://www.geojson.io)) to delimit ocean basins and partitions thereof. The coordinates and plots for the ocean polygons can be found in the code supplement under /data/regions.
