## Appendix S2 - Supplementary figures for "Enhancing georeferenced biodiversity inventories: automated information extraction from literature records reveal the gaps"

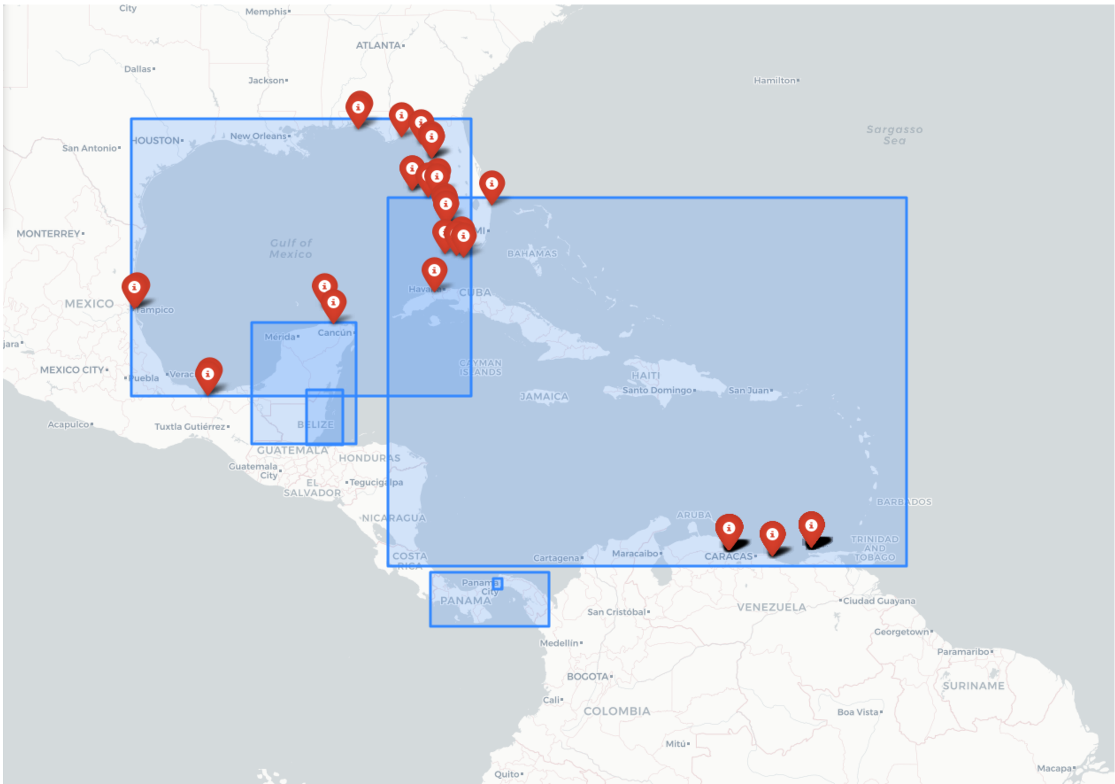


**Figure S1: An example of bounding boxes and OBIS records**. The red balloons indicate OBIS coordinates for *Schizoporella pungens* and blue rectangles the TMO bounding boxes for toponyms associated with *S. pungens*. The toponyms are "Pacific Panama", "Belize", "Panama Canal", "Yucatan Peninsula", "Gulf of Mexico" and "Caribbean" (the largest blue box on this figure). Another toponym included in the TMO data for *S. pungens* is “Fort Pierce Inlet” but it is too small to be visible here. Not shown is the location "Rio de Janeiro", a TMO false positive, but there are two OBIS records from Okinawa, Japan, corroborating evidence that *S. pungens* is a ship-borne invading species (McCann et al. 2019).

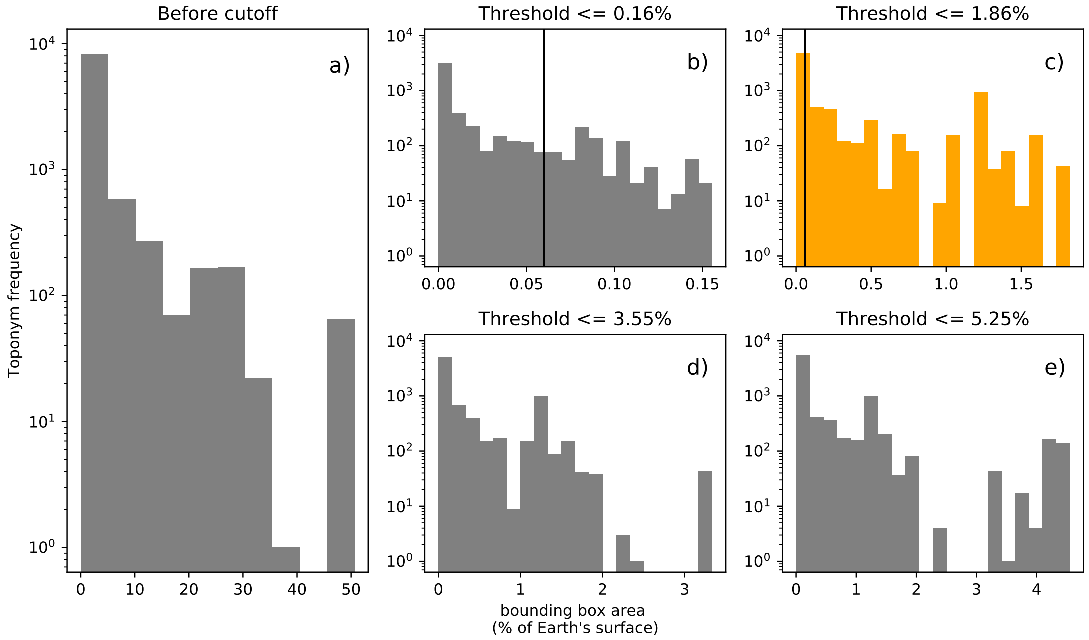


**Figure S2: Toponym bounding box size distributions**. a) shows the distribution prior to filtering, and b)-e) show the distribution after filtering records with bounding boxes larger than areas 0.16, 1.86, 3.55 and 5.25, in units of percentage of the Earth's surface, respectively. 81.9% of all TMO records are included in c), where 1.86% is the threshold that is used for the figures in the main text. Of all TMO records in c), 54% are smaller than a 5° latitude by 5° longitude bin at the equator (area 0.06% of the Earth's surface), indicated by the black vertical line. The frequencies are not unique toponyms, e.g. two mentions of "Adriatic Sea" results in two counts in these figures.

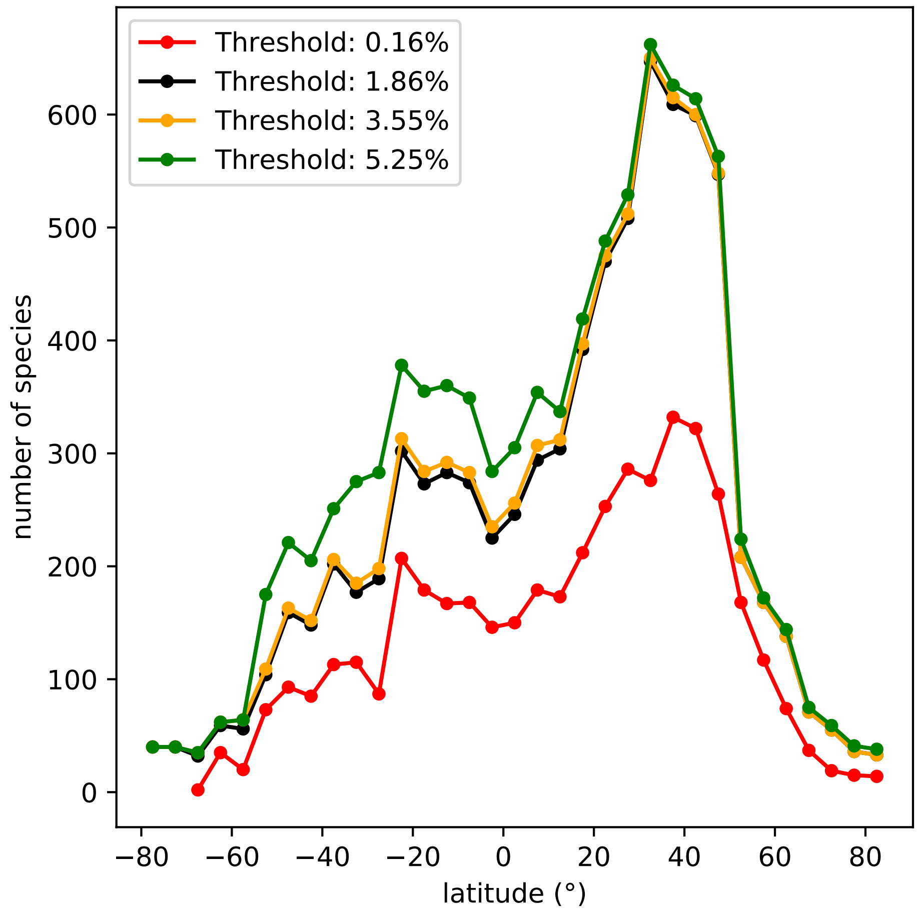


**Figure S3: TMO range-through species richness with varying thresholds for removing toponym locations.** The thresholds are in units of percentage of the Earth's surface. The latitudinal bands are 5° and equiangular. The analyses represented in the main text are based on the 1.86% threshold (black line). See also Fig. S2.

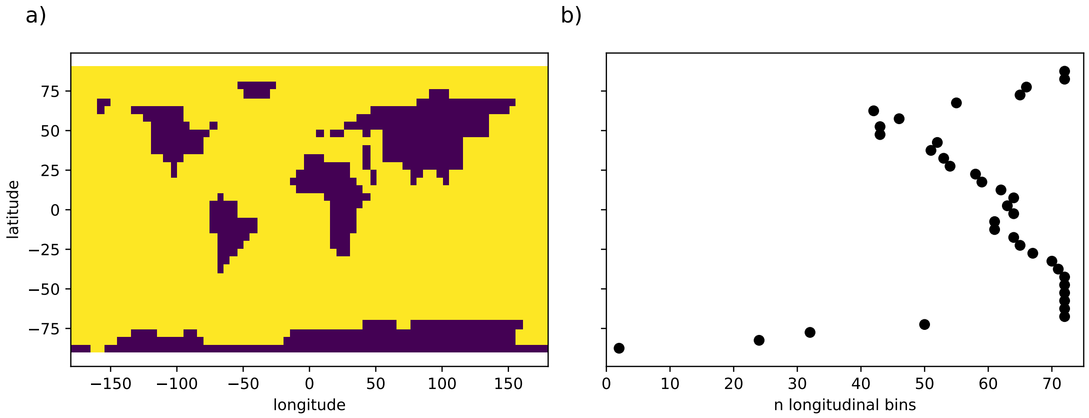


**Figure S4: Landlocked areas**. a) shows 5° by 5° land-locked bins as derived from a 1:10m coastlines map (Patterson, 2019), and b) the distribution of the resulting valid longitudinal sampling bins across latitude.

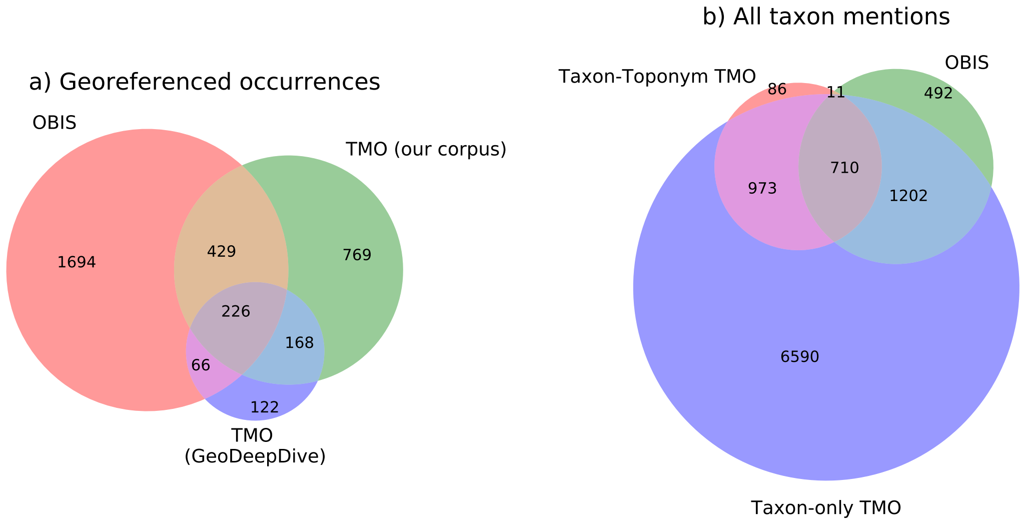


**Figure S5: Species overlap between TMO and OBIS records.** The Venn diagrams show species counts for a) the occurrences we were able to geo-reference, and b) all taxon mentions, regardless of geo-referencing. Taxon-toponym TMO refers to taxon mentions from the candidates discussed in the main text, while Taxon-only refers to all text-mined mentions of any bryozoan species with accepted genus names in the published literature. We were not able to retrieve taxon-only mentions from GeoDeepDive, and hence 86 and 11 species included in the taxon-only set are not included in the taxon-toponym set. Note that for b), the taxon-only TMO species counts sum to 9475, and the total is 10064. These numbers are far higher than the figures described by Bock & Gordon (2013), and Gordon et al. (2019). Since we do not take into account taxonomic revisions and synonymies, our counts also include invalid or outdated species names. This inflates the counts, however the figure still gives an impression of how well the cheilostome taxonomic inventory is represented in TMO and OBIS records.

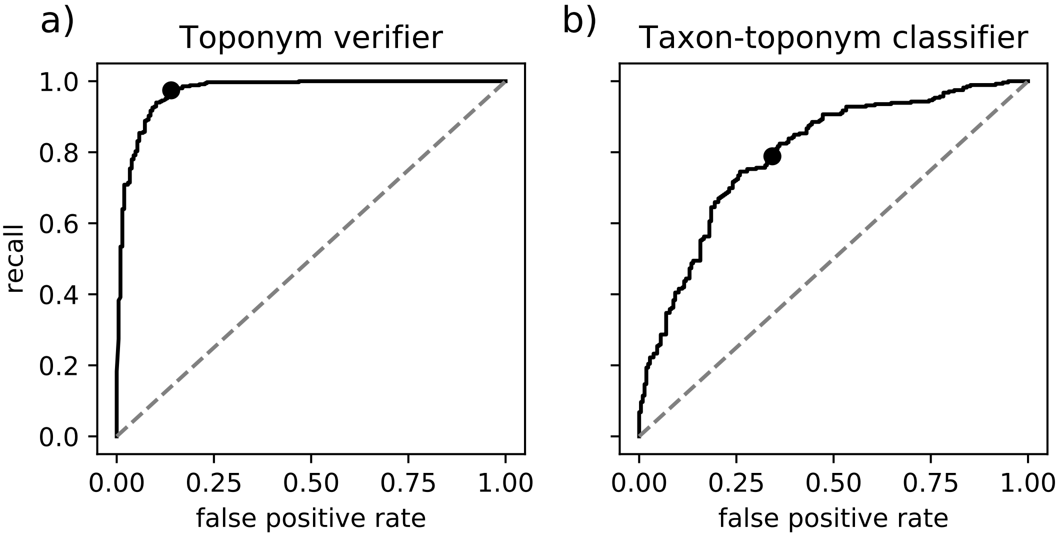


**Figure S6: Classifier receiver operating characteristic (ROC) curves**. a) shows the ROC for the toponym verification classifier. The black line is the performance of the toponym verifier evaluated on the test set (n = 557), and the dashed line is the expected performance for a random classifier. The standard decision boundary of 0.5 (black point) yields a false positive rate of 14% and recall of 97%. b) shows the equivalent for the taxon-toponym relation classifier (false positive rate of 34%, recall of 79%, n = 495).

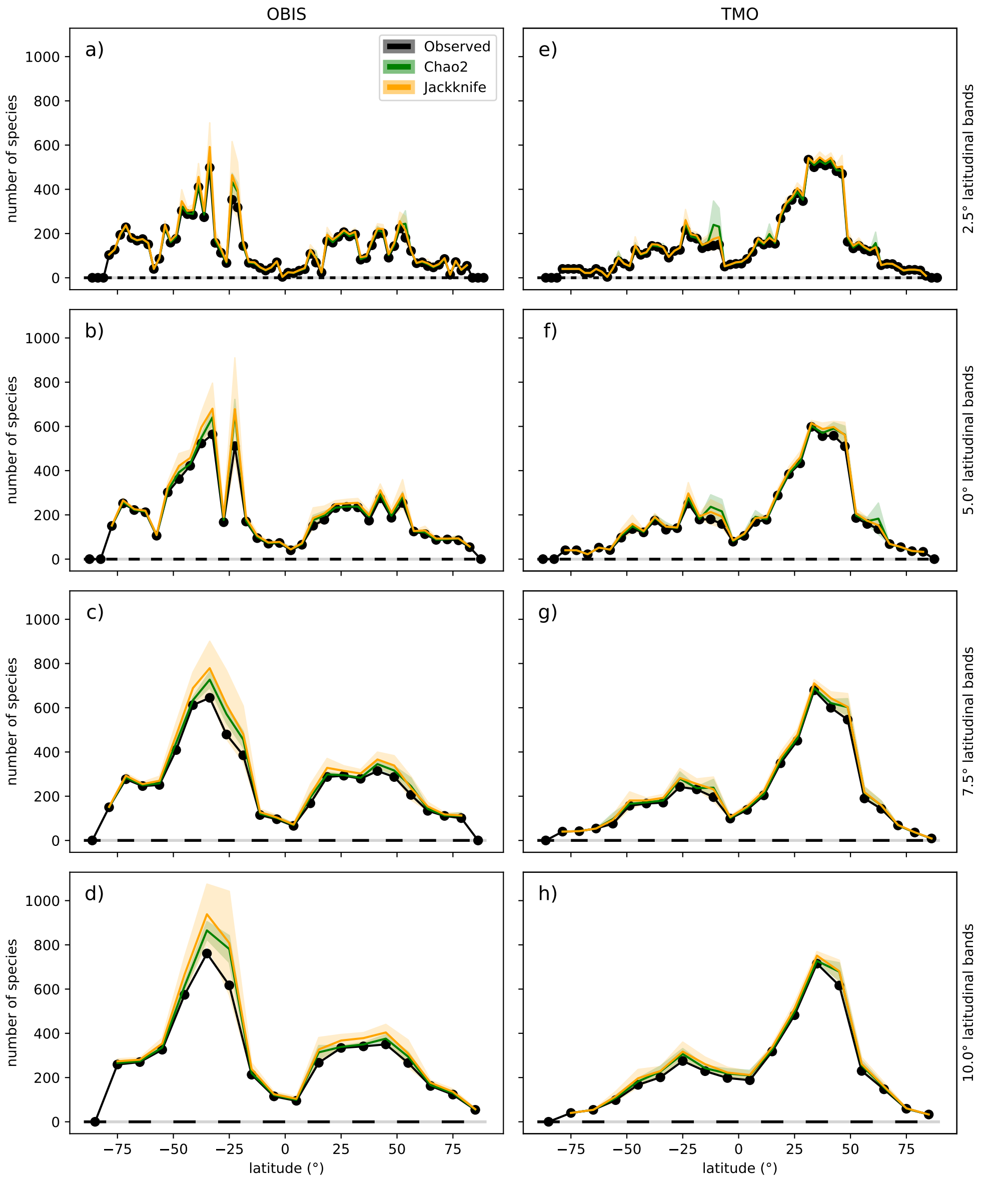


**Figure S7: Species richness estimates with varying latitudinal bands**. Estimates of Chao2 and Jackknife species richness with equiangular latitudinal bands that are 2.5°, 5° (identical to main text Fig. 2a,b), 7.5° and 10.0°. Panels a)-d) use OBIS data, while e)-f) use TMO data. All panels are conditional on 5.0° longitudinal sampling bins.

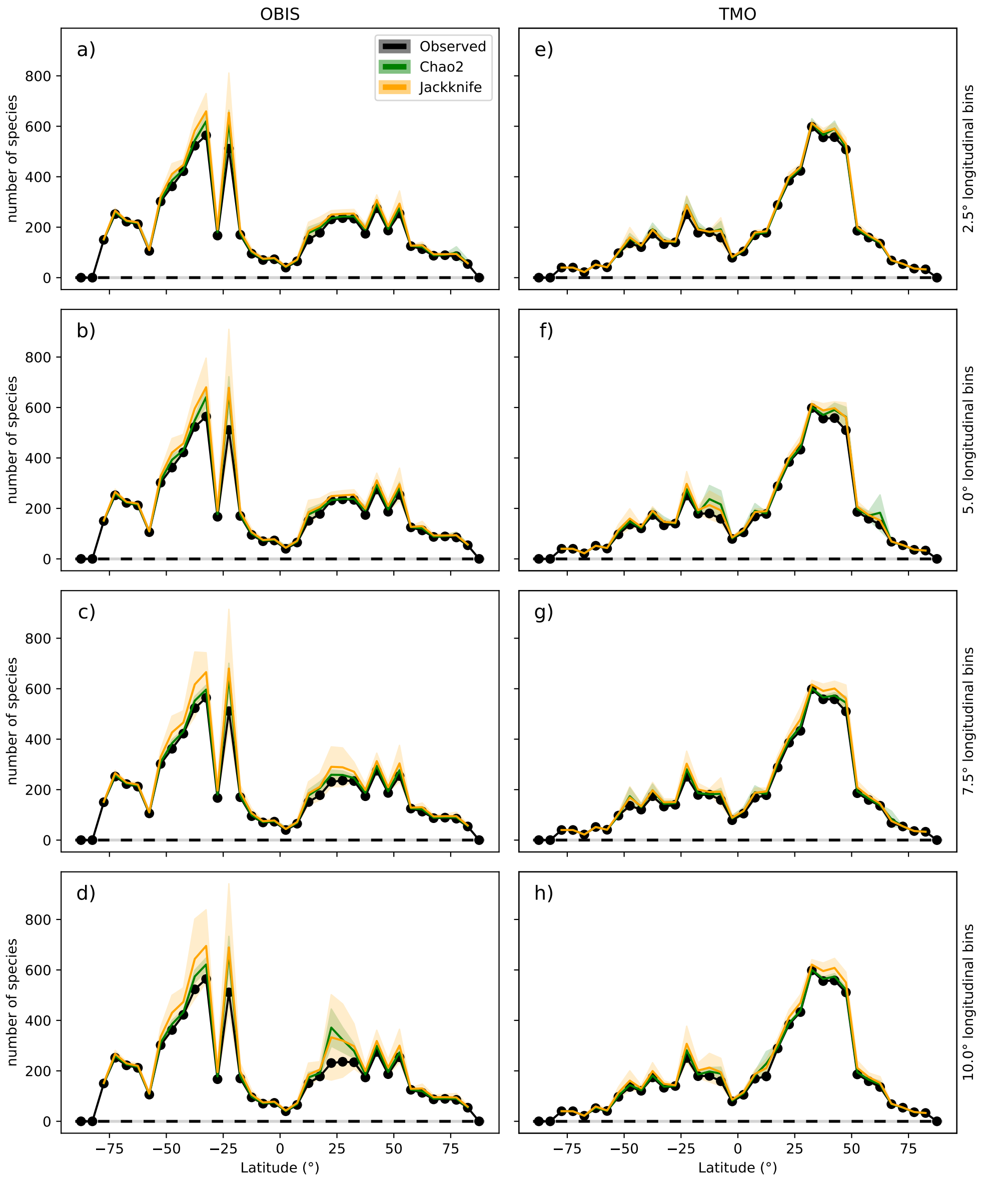


**Figure S8: Species richness estimates with varying longitudinal sampling bins**. Estimates of Chao2 and Jackknife species richness with longitudinal sampling bins that are 2.5°, 5° (identical to main text Fig. 2a,b), 7.5° and 10°. Panels a)-d) use OBIS data, while e)-f) use TMO data. All panels are conditional on 5° equiangular latitudinal bands.

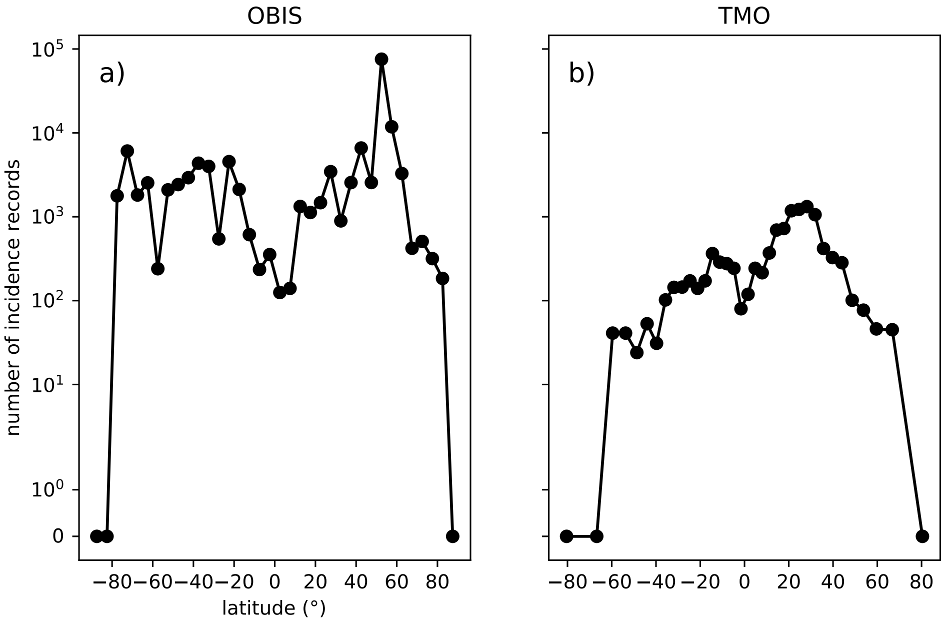


**Figure S9: Number of incidence records across latitude.** Each point represents data from a 5° latitudinal band. The y-axis is linear between 0 and 1, and log-transformed for y > 1.

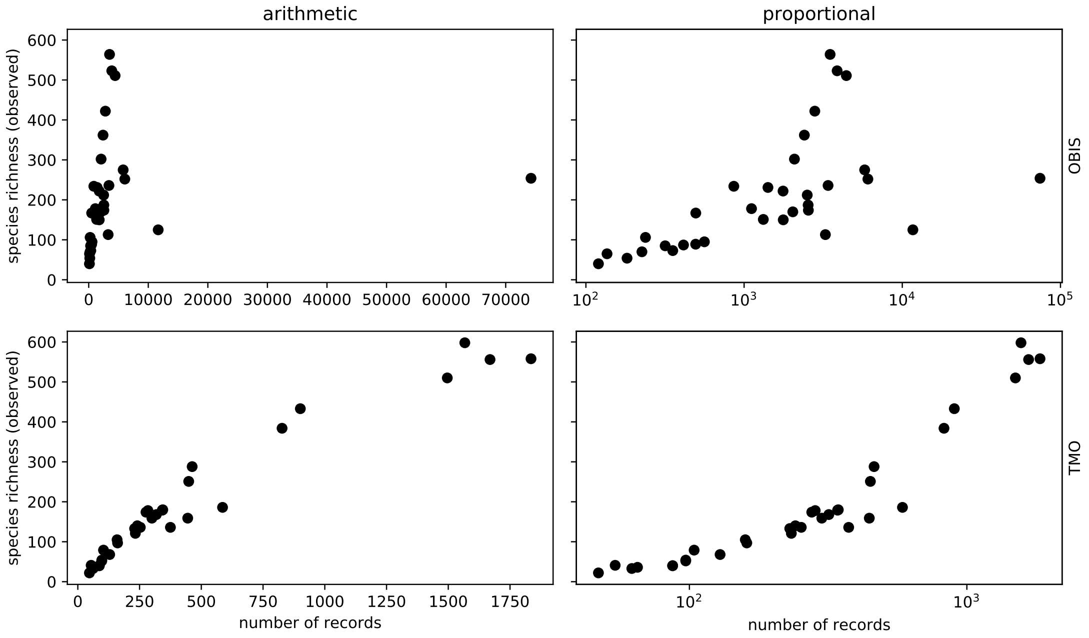


**Figure S10: Species richness versus the number of records**. Each point represents data from a 5° latitudinal band. The species richness estimates are the observed species counts, corresponding with the black lines in Fig. 2a,b. Bands with zero entries are omitted. Both rows represent the same data, however the x-axis is log-transformed in the right column.

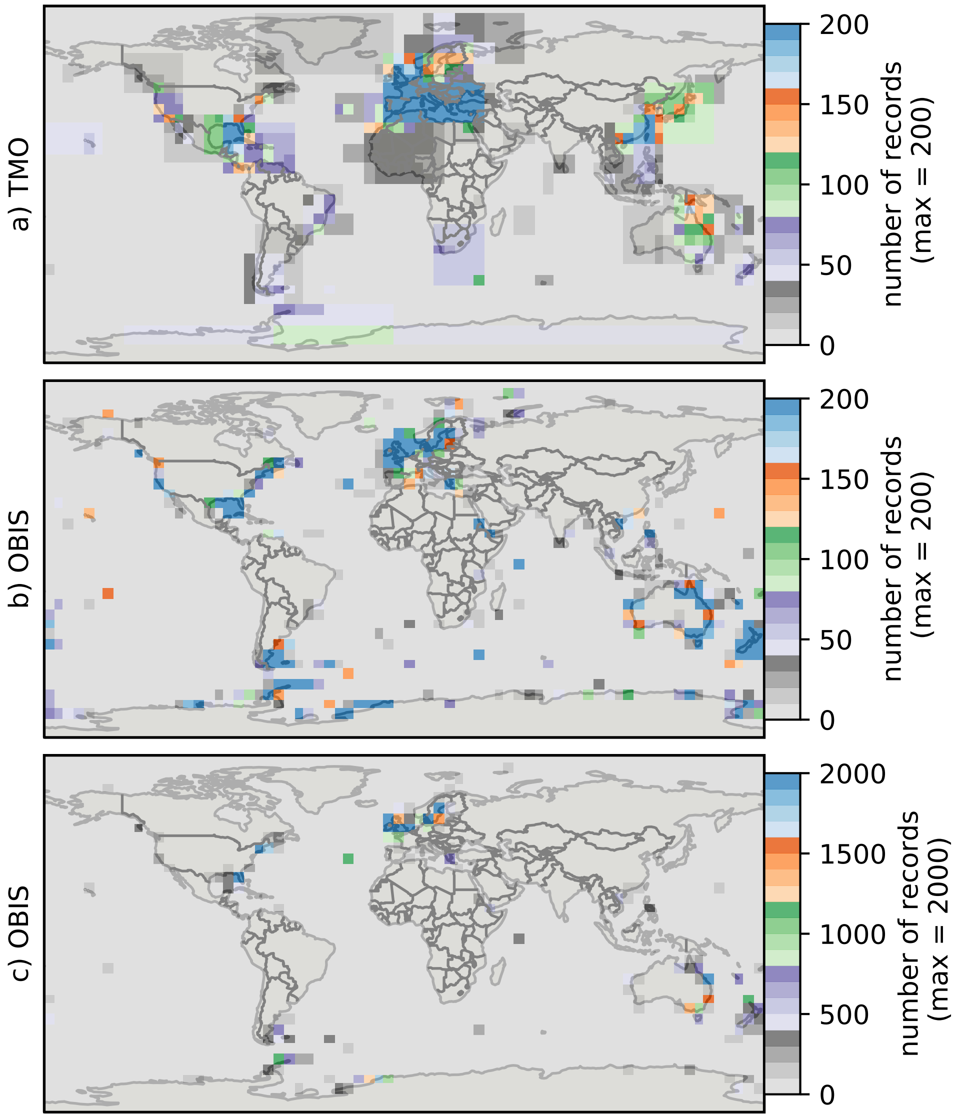


**Figure S11: Heatmaps for cheilostome bryozoan occurrence records per 5° latitude by 5° longitude bins.** The color axes are truncated for visualization purposes, to a maximum of 200, 200 and 2000 in a), b), c), respectively. There are about 750 maximum records per bin in the Mediterranean for the text-mined occurrences (TMO), and about 35000 maximum records in the British Isles for the Ocean Biogeography Information System (OBIS) data. The globe is plotted using the plate carrée projection, but otherwise equal to Fig. 4.
