## Appendix S3 - TMO references for "Enhancing georeferenced biodiversity inventories: automated information extraction from literature records reveal the gaps"


Abbott, M. B. (1973). Seasonal diversity and density in bryozoan populations of Block Island Sound (New York, U.S.A.). In G. P. Larwood (Ed.), *Living and Fossil Bryozoa* (pp. 37–51). London: Academic Press.

Abdelsalam, K. M. (2014). Benthic bryozoan fauna from the northern Egyptian coast. *Egyptian Journal of Aquatic Research*, *40*, 269–282.

Abdelsalam, K. M. (2016a). Fouling bryozoan fauna from Hurghada, Red Sea, Egypt. I. Erect species. *International Journal of Environmental Science and Engineering*, *7*, 59–70.

Abdelsalam, K. M. (2016b). Fouling bryozoan fauna from Hurghada, Red Sea, Egypt. II. Encrusting species. *Egyptian Journal of Aquatic Research*, *42*, 427–436.

Abdelsalam, K. M., & Ramadan, S. E. (2008a). Fouling Bryozoa from some Alexandria harbours, Egypt. (I) Erect species. *Mediterranean Marine Science*, *9*(1), 31–47.

Abdelsalam, K. M., & Ramadan, S. E. (2008b). Fouling Bryozoa from some Alexandria harbours, Egypt. (II) Encrusting species. *Mediterranean Marine Science*, *9*(2), 5–20.

Abdelsalam, K. M., Taylor, P. D., & Dorgham, M. M. (2017). A new species of *Calyptotheca* (Bryozoa: Cheilostomata) from Alexandria, Egypt, southeastern Mediterranean. *Zootaxa*, *4276*(4), 582–590.

Alder, J. (1864). Descriptions of new British Polyzoa, with remarks on some imperfectly known species. *Quarterly Journal of Microscopical Science*, *4*, 95–109.

Almeida, A. C. S., Alves, O., Peso-Aguiar, M., Dominguez, J., & Souza, F. (2015). Gymnolaemata bryozoans of Bahia State, Brazil. *Marine Biodiversity Records*, *8*.

Almeida, A. C., & Souza, F. B. (2014). Two new species of cheilostome bryozoans from the South Atlantic Ocean. *Zootaxa*, *3753*(3), 283–290.

Almeida, A. C., Souza, F. B., & Vieira, L. M. (2017). Malacostegine bryozoans (Bryozoa: Cheilostomata) from Bahia State, northeast Brazil: Taxonomy and non-indigenous species. *Marine Biodiversity*, 1–26.

Almeida, A. C., Souza, F., Farias, J., Alves, F. M. A., & Vieira, L. M. (2018). Bryozoa on disarticulated bivalve shells from Todos os Santos Bay, northeastern Brazil, with the description of two new species. *Zootaxa*, *4434*(3), 401–428.

Almeida, A. C. S., & Souza, F. B. C. (2014). Two new species of cheilostome bryozoans from the South Atlantic Ocean. *Zootaxa*, *3753*(3), 283–290.

Almeida, A. C. S., Souza, F. B. C., Gordon, D. P., & Vieira, L. M. (2015a). The non-indigenous bryozoan *Triphyllozoon* (Cheilostomata: Phidoloporidae) in the Atlantic: Morphology and dispersion on the Brazilian coast. *Zoologia*, *32*(6), 476–484.

Almeida, A. C. S., Souza, F. B. C., Menegola, C. M. S., Sanner, J., & Vieira, L. M. (2014). Taxonomic review of the family Colatooeciidae Winston, 2005 (Bryozoa, Cheilostomata), with description of seven new species. *Zootaxa*, *3868*(1), 1–61.

Almeida, A. C. S., Souza, F. B. C., Menegola, C. M. S., & Vieira, L. M. (2017). Diversity of marine bryozoans inhabiting demosponges in northeastern Brazil. *Zootaxa*, *4290*(2),

281–323.

Almeida, A. C. S., Souza, F. B. C., Sanner, J., & Vieira, L. M. (2015b). Taxonomy of recent Adeonidae (Bryozoa, Cheilostomata) from Brazil, with the description of four new species. *Zootaxa*, *4013*(3), 348–368.

Almeida, A., Souza, F. B., & Vieira, L. M. (2018). A new species of *Cellaria* (Bryozoa: Cheilostomata) from northeastern Brazil, with a tabular identification key to the Atlantic species. *Zoologia (Curitiba)*, *35*.

Alves, D. F., Barros-Alves, S. P., Lima, D. J., Cobo, V. J., & Negreiros-Fransozo, M. L. (2013). Brachyuran and anomuran crabs associated with *Schizoporella unicornis* (Ectoprocta, Cheilostomata) from southeastern Brazil. *Anais Da Academia Brasileira de Ciências*, *85*(1), 245–256.

Amui, A.-M., & Kaselowsky, J. (2006). Bryozoa from the Gulf of Aden and the Red Sea. Part I: Collections from the fifth expedition of the R.V. Meteor. *Fauna of Arabia*, *22*, 7–22.

Anderson, C. M., & Haygood, M. G. (2007). A-Proteobacterial symbionts of marine bryozoans in the genus *Watersipora*. *Applied and Environmental Microbiology*, *73*(1), 303–311.

Arístegui, J. (1985). The genus *Adeonellopsis* MacGillivray (Bryozoa: Cheilostomata) in the Canary Islands: *A. Distoma* (Busk) and *A. Multiporosa* sp. Nov. *Journal of Natural History*, *19*, 425–430.

Ayari, R., & Taylor, P. D. (2014). Some bryozoans from Tunisia, Western Mediterranean Sea. *Studi Trentini Di Scienze Naturali*, *94*, 11–20.

Álvarez, J. A. (1990). Notes on two species of the genus *Turbicellepora* Ryland, 1963 (Bryozoa, Cheilostomatida) of the Atlanto-Mediterranean region: *T. avicularis* (Hincks, 1860) and *t. Magnicostata* (Barroso, 1919). *Cahiers de Biologie Marine*, *31*, 473–483.

Bader, B., & Schäfer, P. (2005). Bryozoans in polar latitudes: Arctic and Antarctic bryozoan communities and facies. *Denisia*, *16*, 263–282.

Balazy, P., & Kuklinski, P. (2013). Mobile hard substrata–An additional biodiversity source in a high latitude shallow subtidal system. *Estuarine, Coastal and Shelf Science*, *119*, 153–161.

Banta, W. C. (1969). The recent introduction of *watersipora arcuata* Banta (Bryozoa, Cheilostomata) as a fouling pest in southern California. *Bulletin of the Southern California Academy of Sciences*, *68*(4), 248–251.

Banta, W. C. (1969). *Watersipora arcuata*, a new species in the subovoidea - cucullata - nigra complex (Bryozoa, Cheilostomata). *Bulletin of the Southern California Academy of Sciences*, *68*, 96–102.

Banta, W. C. (1991). The Bryozoa of the Galapagos. In M. James (Ed.), *Galapagos marine invertebrates: Taxonomy, biogeography and evolution in Darwin's islands* (pp. 371–389). New York: Plenum Publishing Company.

Banta, W. C., Perez, F. M., & Santagata, S. (1995). A setigerous collar in Membranipora chesapeakensis n. Sp.(Bryozoa): Implications for the evolution of cheilostomes from ctenostomes. *Invertebrate Biology*, 83–88. Retrieved from http://www.jstor.org/stable/10.2307/3226957

Banta, W. C., & Santagata, S. (1996). Annotated species list of the Bryozoa of Chuuk, Federated States of Micronesia. *Micronesica*, *29*(2), 305–312.

Barnes, D. K. A. (2006). A most isolated benthos: Coastal bryozoans of Bouvet Island. *Polar Biology*, *29*, 114–119.

Barnes, D. K. A., & Griffiths, H. J. (2008). Biodiversity and biogeography of southern temperate and polar bryozoans. *Global Ecology and Biogeography*, *17*, 84–99.

Barnes, D. K. A., & Kuklinski, P. (2004). Scale-dependent variation in competitive ability among encrusting Arctic species. *Marine Ecology Progress Series*, *275*, 21–32.

Barnes, D. K. A., & Kuklinski, P. (2010). Bryozoans of the Weddell Sea continental shelf, slope and abyss: Did marine life colonize the Antarctic shelf from deep water, outlying islands or in situ refugia following glaciations? *Journal of Biogeography*, *37*(9), 1648–1656.

Barnes, D. K. A., Kuklinski, P., Jackson, J. A., Keel, G. W., Morley, S. A., & Winston, J. E. (2011). Scott's collections help reveal accelerating marine life growth in Antarctica. *Current Biology*, *21*(4), R147–R148. https://doi.org/10.1016/j.cub.2011.01.033

Barnes, D. K. A., Webb, K. E., & Linse, K. (2007). Growth rate and its variability in erect Antarctic bryozoans. *Polar Biology*, *30*, 1069–1081.

Barnes, D. K., & Clarke, A. (1998). The ecology of an assemblage dominant: The encrusting bryozoan *Fenestrulina rugula*. *Invertebrate Biology*, *117*(4), 331–340.

Bastos, A. C., Moura, R. L., Moraes, F. C., Vieira, L. S., Braga, J. C., Ramalho, L. V., … Webster, J. M. (2018). Bryozoans are Major Modern Builders of South Atlantic Oddly Shaped Reefs. *Scientific Reports*, *8*(1), 9638.

Bayer, M. M., & Todd, C. D. (1997). Evidence for zooid senescence in the marine bryozoan *electra pilosa*. *Invertebrate Biology*, *116*(4), 331–340.

Bayer, M. M., Todd, C. D., Hoyle, J. E., & Wilson, J. F. B. (1997). Wave-related abrasion induces formation of extended spines in a marine bryozoan. *Proceedings of the Royal Society of London B*, *264*, 1605–1611.

Beaver, M., & von Dassow, M. (2013). *Feeding the Masses: Colonial Transport in the Marine Bryozoan Membranipora membranacea.* Retrieved from https://digital.lib.washington.edu/researchworks/handle/1773/27199

Benedix, G., Jacob, D. E., & Taylor, P. D. (2014). Bimineralic bryozoan skeletons: A comparison of three modern genera. *Facies*, *60*(2), 389–403. https://doi.org/10.1007/s10347-013-0380-2

Berning, B. (2012). Taxonomic notes on some Cheilostomata (Bryozoa) from Madeira. *Zootaxa*, *3236*, 36–54.

Berning, B. (2013). New and little-known Cheilostomata (Bryozoa, Gymnolaemata) from the NE Atlantic. *European Journal of Taxonomy*, *44*, 1–25. https://doi.org/10.5852/ejt.

2013.44

Berning, B., Achilleos, K., & Wisshak, M. (2019). Revision of the type species of *Metroperiella* (Bryozoa, Cheilostomatida), with the description of two new species. *Journal of Natural History*, *53*(3-4), 141–158. https://doi.org/10.1080/00222933.2019.1582725

Berning, B., Harmelin, J.-G., & Bader, B. (2017). New Cheilostomata (Bryozoa) from NE Atlantic seamounts, islands, and the continental slope: Evidence for deep-sea endemism. *European Journal of Taxonomy*, *347*, 1–51. https://doi.org/10.5852/ejt.2017.347

Berning, B., & Kuklinski, P. (2008). North-east Atlantic and Mediterranean species of the genus *Buffonellaria* (Bryozoa, Cheilostomata): Implications for biodiversity and biogeography. *Zoological Journal of the Linnean Society*, *152*(3), 537–566. https://doi.org/10.1111/j.1096-3642.2007.00379.x

Berning, B., & Ostrovsky, A. N. (2011). *Omanipora pilleri* nov. Gen. Nov. Spec., a new lepraliomorph bryozoan (Cheilostomata) from Oman. *Annalen Des Naturhistorischen Museums in Wien*, *113*, 511–523.

Berning, B., Tilbrook, K. J., & Ostrovsky, A. N. (2014). What, if anything, is a lyrula? *Studi Trentini Di Scienze Naturali*, *94*, 21–28.

Beukhof, E. D., Coolen, J. W. P., van der Weide, B. E., Cuperus, J., de Blauwe, H., & Lust, J. (2016). Records of five bryozoan species from offshore gas platforms rare for the Dutch North Sea. *Marine Biodiversity Records*, *9*(1). https://doi.org/10.1186/s41200-016-0086-6

Bishop, J. D. D. (1989). Colony form and the exploitation of spatial refuges by encrusting Bryozoa. *Biological Reviews*, *64*, 197–218.

Bock, P. E., & Cook, P. L. (2004a). Dimorphic brooding zooids in the genus *Adeona* Lamouroux from Australia (Bryozoa: Cheilostomata). *Memoirs of Museum Victoria*, *61*(2), 129–133.

Bock, P. E., & Cook, P. L. (2004b). New species of the bryozoan genera *Batopora* and Lacrimula (Batoporidae) from Australia. *Proceedings of the Royal Society of Victoria*, *116*(2), 283–288.

Bone, E. K., & Keough, M. J. (2005). Responses to damage in an arborescent bryozoan: Effects of injury location. *Journal of Experimental Marine Biology and Ecology*, *324*(2), 127–140.

Bone, E. K., & Keough, M. J. (2010). Does Polymorphism Predict Physiological Connectedness? A Test Using Two Encrusting Bryozoans. *The Biological Bulletin*, *219*(3), 220–230.

Boonzaaier, M. K., Florence, W. K., & Jones, M. S. (2014). Historical review of South African bryozoology: A legacy of European endeavour. *Annals of Bryozoology 4*, *4*.

Borg, F. (1930). On the bryozoan Fauna of Skelderviken. *Arkiv För Zoologi*, *21 A*(24), 1–13.

Bowden, D. A. (2005a). Quantitative characterization of shallow marine benthic assemblages at Ryder Bay, Adelaide Island, Antarctica. *Marine Biology*, *146*, 1235–1249.

Bowden, D. A. (2005b). Seasonality of recruitment in Antarctic sessile marine benthos. *Marine Ecology Progress Series*, *297*, 101–118.

Bradstock, M., & Gordon, D. P. (1983). Coral-like bryozoan growths in Tasman Bay, and their protection to conserve commercial fish stocks. *New Zealand Journal of Marine and Freshwater Research*, *17*, 159–163.

Branch, M. L., & Hayward, P. (2005). New species of cheilostomatous Bryozoa from subantarctic Marion and Prince Edward Islands. *Journal of Natural History*, *39*(29), 2671–2704.

Branch, M. L., & Hayward, P. J. (2007). The Bryozoa of subantarctic Marion and Prince Edward Islands: Illustrated keys to the species and results of the 1982-1989 University of Cape Town surveys. *African Journal of Marine Science*, *29*(1), 1–24.

Brown, D. A. (1949). On the polyzoan genus *Hippomenella* Canu & Bassler and its genotype Lepralia mucronelliformis Waters. *Journal of the Linnean Society, London (Zoology)*, *41*, 513–520.

Burgess, S. C., & Marshall, D. J. (2011). Temperature-induced maternal effects and environmental predictability. *Journal of Experimental Biology*, *214*(14), 2329–2336. https://doi.org/10.1242/jeb.054718

Busk, G. (1852). *Catalogue of Marine Polyzoa in the Collection of the British Museum. Part II. Cheilostomata (Part)*. London: Trustees of the British Museum.

Busk, G. (1875). *Catalogue of marine Polyzoa in the collection of the British Museum. Part III. Cyclostomata*. London: Trustees of the British Museum.

Cadée, G. C., Chimonides, P. J., & Cook, P. L. (1989). *Pseudolunularia* gen.n. (Cheilostomata), a lunulitiform bryozoan from the Indo-West Pacific. *Zoologica Scripta*, *18*(1), 43–48.

Cancino, J. M., Hughes, R. N., & Orellana, M. C. (1994). *A comparative study of larval release in bryozoans* (P. Hayward, Ed.). Olsen & Olsen, Fredensborg.

Canning-Clode, J., Souto, J., & McCann, L. (2013). First record of textitCelleporaria brunnea (Bryozoa: Lepraliellidae) in Portugal and in the East Atlantic. *Marine Biodiversity Records*, *6*, 1–5.

Cardigos, F., Tempera, F., Ávila, S. P., Gonçalves, J., Colaço, A., & Santos, R. S. (2006). Non-indigenous marine species of the Azores. *Helgoland Marine Research*, *60*, 160–169.

Carlton, J. T., Chapman, J. W., Geller, J. B., Miller, J. A., Ruiz, G. M., Carlton, D. A., … Breitenstein, R. A. (2018). Ecological and biological studies of ocean rafting: Japanese tsunami marine debris in North America and the Hawaiian Islands. *Aquatic Invasions*, *13*(1), 1–9.

Carter, M. C., & Gordon, D. P. (2007). Substratum and morphometric relationships in the bryozoan genus *Odontoporella*, with a description of a new paguridean-symbiont species from New Zealand. *Zoological Science*, *24*, 47–56.

Centurión, R., & López Gappa, J. (2011). Bryozoan assemblages on hard substrata: Species abundance distribution and competition for space. *Hydrobiologia*, *658*, 329–341.

https://doi.org/10.1007/s10750-010-0503-5

César-Aldariz, J., Fernández Pulpeiro, E., & Reverter Gil, O. (1999). A new species of the genus *Celleporella* (Broyzoa: Cheilostomatida) from the European Atlantic coast. *Journal of the Marine Biological Association*, *79*(1), 51–55.

Chainho, P., Fernandes, A., Amorim, A., Ávila, S. P., Canning-Clode, J., Castro, J. J., … Costa, M. J. (2015). Non-indigenous species in Portuguese coastal areas, coastal lagoons, estuaries and islands. *Estuarine, Coastal and Shelf Science*, *167*, 199–211.

Chandramouli, K. H., Zhang, Y., Wong, Y. H., & Qian, P.-Y. (2011). Comparative glycoproteome analysis: Dynamics of protein glycosylation during metamorphic transition from pelagic to benthic life stages in three invertebrates. *Journal of Proteome Research*, *11*(2), 1330–1340.

Cheetham, A. H. (2014). Jeremy Jackson, Bryozoa, and the Punctuated Equilibrium Debate. In S. J. Wyse Jackson Patrick N. (Ed.), *Annals of Bryozoology 4: Aspects of the history of research on bryozoans.* (Vol. 4, pp. 35–52). Dublin: International Bryozoology Association.

Cheetham, A. H., & Hayek, L.-A. C. (1988). Phylogeny reconstruction in the Neogene bryozoan *Metrarabdotos*: A paleontologic evaluation of methodology. *Historical Biology*, *1*(1), 65–83.

Chimenz, C., Nicoletti, L., & Lippi Boncambi, F. (1997). First record of the genus *Retevirgula* in the Mediterranean Sea, with description of R. *Akdenizae* sp. N. (Bryozoa, Cheilostomatida). *Italian Journal of Zoology*, *64*(3), 279–282.

Chimenz Gusso, C., & Tomassetti, P. (2004). Bryozoa Gymnolaemata from the coast of Tunisia and the Strait of Sicily. *Biologia Marina Mediterranea*, *11*(2), 412–415.

Chimonides, P. J., & Cook, P. L. (1994). Notes on the genus *Cranosina* (Bryozoa, Cheilostomida). *Zoologia Scripta*, *23*(1), 43–49.

Chiriboga, A., Ruiz, D., & Banks, S. (2012). CDF Checklist of Galapagos Bryozoans. In *Charles Darwin Foundation Galapagos species checklist.*

Cocito, S., & Ferdeghini, F. (2001). Carbonate standing stock and carbonate production of the bryozoan Pentapora fascialis in the north-western Mediterranean. *Facies*, *45*, 25–30.

Conroy, T. J., Cook, P. L., & Bock, P. E. (2001). New species of *Otionellina* and *Selenaria* (Bryozoa-Cheilostomata) from the South West Shelf, Western Australia. *Transactions of the Royal Society of South Australia*, *125*(1), 15–23.

Cook, P. L. (1966). Some "sand fauna" Polyzoa (Bryozoa) from eastern Africa and the northern Indian Ocean. *Cahiers de Biologie Marine*, *7*, 207–223.

Cook, P. L., & Bock, P. E. (1994). The astogeny and morphology of *Rhabdozoum wilsoni* Hincks (Anasca, Buguloidea). In P. J. Hayward, J. S. Ryland, & P. D. Taylor (Eds.), *Biology and Palaeobiology of Bryozoans* (pp. 47–50). Fredensborg: Olsen & Olsen.

Cook, P. L., & Bock, P. E. (2000). Two new genera of Bryozoa (Calloporidae) from New Zealand. *Journal of Natural History*, *34*(7), 1125–1133.

Cook, P. L., & Hayward, P. J. (1983). Notes on the family Lekythoporidae (Bryozoa, Cheilostomata). *Bulletin of the British Museum (Natural History), Zoology*, *45*(2), 55–76.

Corriero, G., Longo, C., Mercurio, M., Marchini, A., & Occhipinti-Ambrogi, A. (2007). Porifera and Bryozoa on artificial hard bottoms in the Venice Lagoon: Spatial distribution and temporal changes in the northern basin. *Italian Journal of Zoology*, *74*(1), 21–29. https://doi.org/10.1080/11250000601084100

Cortés, J., Nielsen, V., & Herrera-Cubilla, A. (2009). Bryozoans. In I. S. Wehrtmann & J. Cortés (Eds.), *Marine Biodiversity of Costa Rica* (Vol. 86, pp. 385–388, 413–416).

Cranfield, H. J., Gordon, D. P., Willan, R. C., Marshall, B. A., Battershill, C. N., Francis, M. P., … Read, G. B. (1998). Adventive marine species in New Zealand. *NIWA Technical Report*, *34*, 1–48.

Cranfield, H. J., Rowden, A. A., Smith, D. J., Gordon, D. P., & Michael, K. P. (2004). Macrofaunal assemblages of benthic habitat of different complexity and the proposition of a model of biogenic reef habitat regeneration in Foveaux Strait, New Zealand. *Journal of Sea Research*, *52*, 109–125.

Cukrov, M., & Novosel, M. (2014). New records of the estuarine bryozoan *Conopeum seurati* (Canu, 1928) along the eastern (Croatian) coast of the Adriatic Sea. *Studi Trentini Di Scienze Naturali*, *94*, 39–44.

Cumming, R. L. (2015). Two tropical species of *Stephanotheca* (Bryozoa, Cheilostomata, Lanceoporidae) from the Gulf of Carpentaria, Australia. *Zootaxa*, *3948*(2), 279–286.

Cumming, R. L., & Sebastian, P. (2018). New encrusting species of Lanceoporidae (Bryozoa, Cheilostomata) from the southern Great Barrier Reef, Australia. *Zootaxa*, *4500*(1), 104. https://doi.org/10.11646/zootaxa.4500.1.6

Cumming, R. L., & Tilbrook, K. J. (2014). Six species of *Calyptotheca* (Bryozoa, Cheilostomata, Lanceoporidae) from the Gulf of Carpentaria and northern Australia, with description of a new species. *Zootaxa*, *3827*(2), 147–169.

De Blauwe, H. (2005). A new species of Caulibugula (Bryozoa: Cheilostomatida) from France. *Bulletin de L'Institut Royal Des Sciences Naturelles de Belgique, Biologie*, *75*, 81–87.

De Blauwe, H. (2006). On the taxonomy and distribution of the family Pacificincolidae Liu & Liu, 1999 (Bryozoa, Cheilostomata), with the description of a new genus. *Bulletin de L'Institut Royal Des Sciences Naturelles de Belgique, Biologie*, *76*, 139–145.

De Blauwe, H., & Faasse, M. (2001). Extension of the range of the bryozoans Tricellaria inopinata and Bugula simplex in the North-East Atlantic Ocean (Bryozoa: Cheilostomatida). *Nederlandse Faunistische Mededelingen*, *14*, 103–112.

De Blauwe, H., & Faasse, M. (2004). Smittoidea prolifica Osburn, 1952 (Bryozoa, Cheilostomatida), a Pacific bryozoan introduced to The Netherlands (Northeast Atlantic). *Bulletin de L'Institut Royal Des Sciences Naturelles de Belgique, Biologie*, *74*, 33–39.

De Blauwe, H., & Gordon, D. P. (2014). New bryozoan taxa from a biodiversity hotspot in the Eastern Weddell Sea. *Studi Trentini Di Scienze Naturali*, *94*, 53–78.

De Blauwe, H., Kind, B., Kuhlenkamp, R., Cuperus, J., van der Weide, B., & Kerckhof, F. (2014). Recent observations of the introduced *Fenestrulina Delicia* Winston, Hayward & Craig, 2000 (Bryozoa) in Western Europe. *Studi Trentini Di Scienze Naturali*, *94*, 45–51.

de la Nuez-Hernández, D., Valle, C., Forcada, A., Gonzálz-Correa, J. M., & Fernández Torquemada, Y. (2014). Assessing the erect bryozoan *Myriapora truncata* (Pallas, 1766) as indicator of recreational diving impact on coralligenous reef communities. *Ecological Indicators*, *46*, 193–200.

Denisenko, N. (2015). New species of the genus *Parasmittina* (Bryozoa: Cheilostomata: Smittinidae) from the Chukchi Sea. *Zoosystemica Rossica*, *24*(2), 303–306.

Denisenko, N. V. (2011). Bryozoans of the East Siberian Sea: History of research and current knowledge of diversity. In *3rd Ser. Annals of Bryozoology 3: Aspects of the history of research on bryozoans* (pp. 1–15).

Denisenko, N. V. (2014). Deep-sea fauna of European seas: An annotated species check-list of benthic invertebrates living deeper than 2000 m in the seas bordering Europe. Bryozoa. *Invertebrate Zoology*, *11*(1), 89–98.

Denisenko, N. V. (2016). A new species of the genus *Callopora* (Bryozoa: Cheilostomatida: Calloporidae) from the Barents and Kara Seas. *Zoosystematica Rossica*, *25*(2), 183–188.

Denisenko, N. V. (2016). Two new species of the genus *Turbicellepora* Ryland, 1963 (Bryozoa: Celleporidae) found on Lophelia coral from the Greenland slope. *Zootaxa*, *4066*(2), 177–182.

Denisenko, N. V. (2018). New Cheilostomata (Bryozoa) species from sublittoral and bathyal zones off the Faroe Islands, with some comments on allied taxa. *Zootaxa*, *4375*(1), 116–126.

Denisenko, N. V., & Kukliński, P. (2008). Historical development of research and current state of bryozoan diversity in the Chukchi Sea. In P. N. Wyse Jackson & S. Jones (Eds.), *Annals of Bryozoology 2: Aspects of the history of research on bryozoans.* (pp. 35–50). Dublin: International Bryozoology Association.

Denisenko, N. V., Thomsen, E., & Tendal, O. S. (2013). Bryozoan epifauna on brachiopods from the Faroe Islands (NE Atlantic). *Fró skaparrit*, *60*, 96–113.

d'Hondt, J.-L. (2002). The French Pre-Lamarckian bryozoologists. In P. N. Wyse Jackson & M. E. Spencer Jones (Eds.), *Annals of Bryozoology: Aspects of the history of research on bryozoans* (pp. 81–95). Dublin: International Bryozoology Association.

d'Hondt, J.-L. (2008). The historical collections of Recent Bryozoa in the French National Collections. In P. N. Wyse Jackson & M. E. Spencer Jones (Eds.), *Annals of Bryozoology 2: Aspects of the history of research on bryozoans.* (pp. 59–70). Dublin: International Bryozoology Association.

d'Hondt, J.-L., & Mascarell, G. (2004). On some Bryozoa collected during the northern cruises of the "Pourquoi-Pas" during the years 1921-1929 and description of a new species. *Cahiers de Biologie Marine*, *45*, 281–285.

Dick, M. H. (2008). Unexpectedly high diversity of *Monoporella* (Bryozoa: Cheilostomata) in the Aleutian Islands, Alaska: Taxonomy and distribution of six new species. *Zoological*

*Science*, *25*, 36–52.

Dick, M. H., & Grischenko, A. V. (2017). Rocky-intertidal cheilostome bryozoans from the vicinity of the Sesoko Biological Station, west-central Okinawa, Japan. *Journal of Natural History*, *51*(3-4), 141–266.

Dick, M. H., Grischenko, A. V., & Mawatari, S. F. (2005). Intertidal Bryozoa (Cheilostomata) of Ketchikan, Alaska. *Journal of Natural History*, *39*(43), 3687–3784.

Dick, M. H., Mawatari, S. F., Sanner, J., & Grischenko, A. V. (2011). Cribrimorph and Other Cauloramphus Species (Bryozoa: Cheilostomata) from the Northwestern Pacific. *Zoological Science*, *28*(2), 134–147. https://doi.org/10.2108/ZSJ.28.134

Dick, M. H., Tilbrook, K. J., & Mawatari, S. F. (2006). Diversity and taxonomy of rocky-intertidal Bryozoa on the Island of Hawaii, USA. *Journal of Natural History*, *40*(38-40), 2197–2257.

D'Onghia, G., Cape, F., uto, Cardone, F., Carlucci, R., Carluccio, A., Chimienti, G., … Tursi, A. (2015). Macro- and megafauna recorded in the submarine Bari Canyon (southern Adriatic, Mediterranean Sea) using different tools. *Mediterranean Marine Science*, *16*(1), 180–196.

Dumont, J. P. C. (1981). A report on the cheilostome Bryozoa of the Sudanese Red Sea. *Journal of Natural History*, *15*, 623–637.

Durrant, H. M. S., Clark, G. F., Dworjanyn, S. A., Byrne, M., & Johnston, E. L. (2013). Seasonal variation in the effects of ocean warming and acidification on a native bryozoan, *Celleporaria nodulosa. Marine Biology*, *2013*(8), 1903–1911. https://doi.org/DOI 10.1007/s00227-012-2008-4

Dyrynda, P. E. J., & Ryland, J. S. (1982). Reproductive strategies and life histories in the cheilostome marine bryozoans *Chartella papyracea* and *Bugula* flabellata. *Marine Biology*, *71*, 241–256.

Faasse, M., Van Moorsel, G., & Tempelman, D. (2013). Moss animals of the Dutch part of the North Sea and coastal waters of the Netherlands (Bryozoa). *Nederlandse Faunistische Mededelingen*, *41*, 1–14.

Fehlauer-Ale, K. H., Mackie, J. A., Lim-Fong, G. E., Ale, E., Pie, M. R., & Waeschenbach, A. (2014). Cryptic species in the cosmopolitan *Bugula neritina* complex (Bryozoa, Cheilostomata). *Zoologica Scripta*, *43*(2), 193–205.

Fehlauer-Ale, K. H., Winston, J. E., Tilbrook, K. J., Nascimento, K. B., & Vieira, L. M. (2015). Identifying monophyletic groups within Bugula sensu lato (Bryozoa, Buguloidea). *Zoologica Scripta.*

Ferrario, J., d'Hondt, J.-L., Marchini, A., & Occhipinti-Ambrogi, A. (2015). From the Pacific Ocean to the Mediterranean Sea: Watersipora arcuata, a new non-indigenous bryozoan in Europe. *Marine Biology Research*, *11*(9), 909–919.

Ferrario, J., Rosso, A., Marchini, A., & Occhipinti-Ambrogi, A. (2018). Mediterranean non-indigenous bryozoans: An update and knowledge gaps. *Biodiversity and Conservation*, *27*(11), 2783–2794. https://doi.org/10.1007/s10531-018-1566-2

Ferretti, C., Magnino, G., & Balduzzi, A. (2007). Morphology of the larva and ancestrula of *Myriapora truncata* (Bryozoa, Cheilostomatida). *Italian Journal of Zoology*, *74*(4), 341–350. https://doi.org/10.1080/11250000701629572

Figuerola, B., Angulo-Preckler, C., Núñez-Pons, L., Moles, J., Sala-Comorera, L., García-Aljaro, C., … Avila, C. (2017). Experimental evidence of chemical defence mechanisms in Antarctic bryozoans. *Marine Environmental Research*, *129*, 68–75.

Figuerola, B., Ballesteros, M., & Avila, C. (2013). Description of a new species of Reteporella (Bryozoa: Phidoloporidae) from the Weddell Sea (Antarctica) and the possible functional morphology of avicularia. *Acta Zoologica*, *94*(1), 66–73.

Figuerola, B., Barnes, D. K., Brickle, P., & Brewin, P. E. (2017). Bryozoan diversity around the Falkland and South Georgia Islands: Overcoming Antarctic barriers. *Marine Environmental Research*, *126*, 81–94.

Figuerola, B., Gordon, D. P., & Cristobo, J. (2018). New deep Cheilostomata (Bryozoa) species from the Southwestern Atlantic: Shedding light in the dark. *Zootaxa*, *4375*(2), 211–249. https://doi.org/10.11646/zootaxa.4375.2.3

Figuerola, B., Gore, D. B., Johnstone, G., & Stark, J. S. (2019). Spatio-temporal variation of skeletal Mg-calcite in Antarctic marine calcifiers. *PLOS ONE*, *14*(5). https://doi.org/10.1371/journal.pone.0210231

Figuerola, B., Monleón-Getino, T., Ballesteros, M., & Avila, C. (2012). Spatial patterns and diversity of bryozoan communities from the Southern Ocean: South Shetland Islands, Bouvet Island and Eastern Weddell Sea. *Systematics and Biodiversity*, *10*(1), 109–123.

Figuerola, B., Núñez-Pons, L., Moles, J., & Avila, C. (2013). Feeding repellence in Antarctic bryozoans. *Naturwissenschaften*, *100*, 1069–1081. https://doi.org/10.1007/s00114-013-1112-8

Figuerola, B., Núñez-Pons, L., Monleón-Getino, T., & Avila, C. (2014). Chemo-ecological interactions in Antarctic bryozoans. *Polar Biology*, *37*, 1017–1030.

Figuerola, B., Sala-Comorera, L., Angulo-Preckler, C., Vázquez, J., Montes, M. J., García-Aljaro, C., … Avila, C. (2014). Antimicrobial activity of Antarctic bryozoans: An ecological perspective with potential for clinical applications. *Marine Environmental Research*, *101*, 52–59.

Figuerola, B., Taboada, S., Monleón-Getino, T., Vázquez, J., & Avila, C. (2013). Cytotoxic activity of Antarctic benthic organisms against the common sea urchin *Sterechinus neumayeri*. *Oceanography*, *1*(107), 2.

Florence, W. K., Hayward, P. J., & Gibbons, M. J. (2007). Taxonomy of shallow-water Bryozoa from the west coast of South Africa. *African Natural History*, *3*, 1–58.

Fortunato, H. (2015). Bryozoans in climate and ocean acidification research: A reappraisal of an under-used tool. *Regional Studies in Marine Science*, *2, Supplement*, 32–44. https://doi.org/10.1016/j.rsma.2015.08.010

Fortunato, H., Roloff, A., & Schäfer, P. (2014). Salinity effects on growth and development of Flustra foliacea (L.) Zooids: Experimental data. *Studi Trentini Di Scienze Naturali*, *94*, 111–118.

Fortunato, H., Schäfer, P., & Blaschek, H. (2012). Growth rates, age determination, and calcification levels in *Flustra foliacea* (L.) (Bryozoa: Cheilostomata): Preliminary assessment. In A. Ernst, P. Schäfer, & J. Scholz (Eds.), *Bryozoan Studies 2010* (pp. 59–74). Berlin: Springer.

Fortunato, H., & Spencer Jones, M. (2014). Collections and climate change research: *Flustra foliacea* (L.) (Bryozoa) in the Natural History Museum, London. *International Bryozoology Association, Dublin*, 53–73.

Franjević, D., Novosel, M., & Koletić, N. (2015). Freshwater and brackish bryozoan species of Croatia (Bryozoa: Gymnolaemata, Phylactolaemata) and their genetic identification. *Zootaxa*, *4032*(2), 221–228.

Fransen, C. H. J. M. (1986). Caribbean Bryozoa: Anasca and Ascophora Imperfecta of the inner bays of Curaçao and Bonaire. *Studies on the Fauna of Curaçao and Other Caribbean Islands*, *68*, 1–119.

Fuchs, J., Martindale, M. Q., & Hejnol, A. (2011). Gene expression in bryozoan larvae suggest a fundamental importance of pre-patterned blastemic cells in the bryozoan life-cycle. *EvoDevo*, *2*(1), 1–15.

Førde, H., Forbord, S., Handå, A., Fossberg, J., Arff, J., Johnsen, G., & Reitan, K. I. (2016). Development of bryozoan fouling on cultivated kelp (*Saccharina latissima*) in Norway. *Journal of Applied Phycology*, *28*(2), 1225–1234. https://doi.org/10.1007/s10811-015-0606-5

Gavira-O'Neill, K., Guerra-García, J. M., Moreira, J., & Ros, M. (2018). Mobile epifauna of the invasive bryozoan Tricellaria inopinata: Is there a potential invasional meltdown? *Marine Biodiversity*, *48*(2), 1169–1178. https://doi.org/10.1007/s12526-016-0563-5

Gerovasileiou, V., & Rosso, A. (2016). Marine Bryozoa of Greece: An annotated checklist. *Biodiversity Data Journal*, *4*, e10672. https://doi.org/doi: 10.3897/BDJ.4.e10672

Gluhak, T., Lewis, J. E., & Popijac, A. (2007). Bryozoan fauna of Green Island, Taiwan: First indications of biodiversity. *Zoological Studies*, *46*(4), 397–426.

Gontar, V. (2017). A new species of the genus Isoschizoporella (Bryozoa: Gymnolaemata: Cheilostomatida: Eminooeciidae) from the Weddell Sea Isoschizoporella (Bryozoa: Gymnolaemata: Cheilostomatida: Eminooeciidae) . *ZOOSYSTEMATICA ROSSICA*, *26*(2), 209–211.

Gontar, V. I. (1993). New deepwater species of Cheilostomida from the Kuril Islands and the Pacific Ocean (Bryozoa). *Zoosystematica Rossica*, *2*(1), 41–45.

Gontar, V. I. (2008). Three new species of the genus Smittina from the Weddell Sea, Antarctic (Bryozoa: Cheilostomata: Smittinidae). *Zoosystematica Rossica*, *17*(1), 7–9.

Gontar, V. I. (2012a). Evolution and distribution of the Antarctic and Arcitc cheilostomate Bryozoa. *Ecology & Safety*, *6*(1), 206–211.

Gontar, V. I. (2012b). The fauna of Bryozoa Cheilostomata of the Black Sea. *Ecology & Safety*, *6*(3), 100–129.

Gontar, V. I. (2013). Bryozoa of the Russian sea shore of the Baltic Sea. *Journal of International Scientific Publications: Ecology & Safety*, *7*(3), 99–107.

Gontar, V. I. (2014). Invasive species in the fauna of Bryozoa in the Bering and the Chuckchi Seas. *Journal of International Scientific Publications: Ecology & Safety*, *8*, 354–360.

Gontar, V. I. (2014). New additions to the fauna of Bryozoa Cheilostomata of the Black Sea. *Journal of International Scientific Publications: Ecology and Safety*, *8*, 361–369.

Gordon, D. P. (1971). Zooidal budding in the cheilostomatous Bryozoan Fenestrulina malusii var. Thyreophora. *New Zealand Journal of Marine and Freshwater Research*, *5*, 453–460.

Gordon, D. P. (1972). Biological relationships of an intertidal bryozoan population. *Journal of Natural History*, *6*, 503–514.

Gordon, D. P. (1974). Microarchitecture and function of the lophophore in the bryozoan *Cryptosula pallasiana*. *Marine Biology*, *27*(2), 147–163. https://doi.org/10.1007/BF00389068

Gordon, D. P. (1985). Additional species and records of gymnolaemate Bryozoa from the Kermadec region. *New Zealand Oceanographic Institute Records*, *4*(14), 159–183.

Gordon, D. P. (1988). The bryozoan families Sclerodomidae, Bifaxariidae, and Urceoliporidae and a novel type of frontal wall. *New Zealand Journal of Zoology*, *15*, 249–290.

Gordon, D. P. (1989a). Intertidal bryozoans from coral reef-flat rubble, Sa'aga, Western Samoa. *New Zealand Journal of Zoology*, *16*, 447–463.

Gordon, D. P. (1989b). New and little-known deep-sea taxa of umbonulomorph Bryozoa and their classification. *New Zealand Journal of Zoology*, *16*, 251–267.

Gordon, D. P. (1993). Bryozoa: The ascophorine infraorders Cribriomorpha, Hippothoomorpha and Umbonulomorpha mainly from New Caledonian waters. *Mémoires Du Muséum National d'Histoire Naturelle*, *158*, 299–347.

Gordon, D. P. (2009a). *Baudina* gen. Nov., constituting the first record of Pasytheidae from Australia, and Sinoflustridae fam. Nov., with a checklist of Bryozoa and Pterobranchia from Beagle Gulf. *The Beagle, Records of the Museums and Art Galleries of the Northern Territory*, *25*, 43–54.

Gordon, D. P. (2009b). New bryozoan taxa from a new marine conservation area in New Zealand, with a checklist of Bryozoa from Greater Cook Strait. *Zootaxa*, *1987*, 39–60.

Gordon, D. P. (2016). Bryozoa of the South China Sea - an overview. *Raffles Bulletin of Zoology*, *34*((Supplement)), 604–618.

Gordon, D., & Parker, S. (1991). An aberrant new genus and subfamily of the spiculate bryozoan family Thalamoporellidae epiphytic on *Posidonia*. *Journal of Natural History*, *25*(5), 1363–1378. https://doi.org/10.1080/00222939100770851

Gordon, D. P., & Grischenko, A. V. (1994). Bryozoan frontal shields: The types species of Desmacystis, Ramphostomella, Rhamphosmittina, Rhamphostomellina, and new genus Arctonula. *Zoologia Scripta*, *23*(1), 61–72.

Gordon, D. P., & Hastings, A. B. (1979). The interzooidal communications of *Hippothoa* sensu lato (Bryozoa) and their value in classification. *Journal of Natural History*, *13*, 561–579.

Gordon, D. P., & Mawatari, S. F. (1992). Atlas of marine-fouling Bryozoa of New Zealand ports and harbours. *Miscellaneous Publications New Zealand Oceanographic Institute*, *107*, 1–52.

Gordon, D. P., Mawatari, S. F., & Kajihara, H. (2002). New taxa of Japanese and New Zealand Eurystomellidae (Phylum Bryozoa) and their phylogenetic relationships. *Zoological Journal of the Linnean Society*, *136*, 199–216.

Gordon, D. P., & Parker, S. A. (1991). The plectriform apparatus - an enigmatic structure in malacostegine Bryozoa. In F. P. Bigey (Ed.), *Bryozoaires Actuels et Fossiles: Bryozoa Living and Fossil* (pp. 133–145). Nantes: Bulletin de la Société Sciences Naturelles de l'Ouest de la France, Mémoire HS 1.

Gordon, D. P., Ramalho, L. V., & Taylor, P. D. (2006). An unreported invasive bryozoan that can affect livelihoods - Membraniporopsis tubigera in New Zealand and Brasil. *Bulletin of Marine Science*, *78*(2), 331–342.

Gordon, D. P., & Rudman, W. B. (2006). *Integripelta acanthus* n. Sp. (Bryozoa: Eurystomellidae) - a tropical prey species of *Okenia hiroi* (Nudibranchia). *Zootaxa*, *1229*, 41–48.

Gordon, D. P., & Spencer Jones, M. E. (2013). The amathiiform Ctenostomata (phylum Bryozoa) of New Zealand - including four new species, two of them of probable alien origin. *Zootaxa*, *3647*(1), 75–95.

Gordon, D. P., & Taylor, P. D. (2017). Resolving the status of *Pyriporoides* and *Daisyella* (Bryozoa: Cheilostomata), with the systematics of some additional taxa of Calloporoidea having an ooecial heterozooid. *Zootaxa*, *4242*(2), 201–232. Retrieved from http://mapress.com/j/zt/article/view/zootaxa.4242.2.1

Gómez, A., Wright, P. J., Lunt, D. H., Cancino, J. M., Carvalho, G. R., & Hughes, R. N. (2007). Mating trials validate the use of DNA barcoding to reveal cryptic speciation of a marine bryozoan taxon. *Proceedings of the Royal Society of London, B*, *274*, 199–207.

Grischenko, A. V. (2002). History of investigations and current state of knowledge of bryozoan species diversity in the Bering Sea. In P. N. Wyse Jackson & M. E. Spencer Jones (Eds.), *Annals of Bryozoology: Aspects of the history of research on bryozoans* (pp. 97–116). Dublin: International Bryozoology Association.

Grischenko, A. V. (2013). First record of a bathyal bryozoan fauna from the Sea of Japan. *Deep-Sea Research II*, *86-87*, 172–180.

Grischenko, A. V. (2014). An unknown archive of German A. Kluge: A bryozoans collection in the Museum of Invertebrate Zoology at Perm State National Research University. In *Annals of Bryozoology 4: Aspects of the history of research on bryozoans.* (Vol. 4, pp. 75–92).

Grischenko, A. V., & Chernyshev, A. V. (2015). Triticella minini–a new ctenostome bryozoan from the abyssal plain adjacent to the Kuril–Kamchatka Trench. *Deep Sea Research Part II: Topical Studies in Oceanography*, *111*, 343–350.

Grischenko, A. V., Gordon, D. P., & Taylor, P. D. (1998). A unique new genus of cheilostomate bryozoan with reversed-polarity zooidal budding. *Asian Marine Biology*, *15*, 105–117.

Grischenko, A. V., & Mawatari, S. F. (2006). On the bryozoan diversity of Sagami Bay. *Mem Nat Sci Mus Tokyo*, *40*, 187–201.

Grischenko, A. V., Taylor, P. D., & Mawatari, S. F. (2004). Doryporella smirnovi sp. Nov. (Bryozoa: Cheilostomata) and its impact on phylogeny and classification. *Zoological Science*, *21*, 327–332.

Gruhl, A. (2008). Muscular systems in gymnolaemate bryozoan larvae (Bryozoa: Gymnolaemata). *Zoomorphology*, *127*(3), 143–159. https://doi.org/10.1007/s00435-008-0059-3

Gruhl, A. (2010). Ultrastructure of mesoderm formation and development in Membranipora membranacea (Bryozoa: Gymnolaemata). *Zoomorphology*, *129*(1), 45–60.

Grunbaum, D. (1997). Hydromechanical mechanisms of colony organization and cost of defense in an encrusting bryozoan, Membranipora membranacea. *Limnology and Oceanography*, *42*(4), 741–752. Retrieved from://000071037700012

Hall, D. N. (1982). Larval release in Celleporaria apiculata (Busk) (Bryozoa: Ascophora). *Journal of Natural History*, *16*, 195–200.

Hamond, R. (1975). The marine and brackish-water Bryozoa of Norfolk. *Transactions of the Norfolk and Norwich Naturalists' Society*, *22*(6), 406–423.

Hara, U. (2000). Bryozoan fragments from Eocene glacial erratics of McMurdo Sound, East Antarctica. In J. D. Stilwel & R. M. Feldman (Eds.), *Paleobiology and paleoenvironments of eocene fossiliferous erratics, mcmurdo sound, east antarctica, geophysical union, antarctic research series* (Vol. 76, pp. 321–323).

Harmelin, J.-G. (2014). *Monoporella bouchardii* (Audouin & Savigny, 1826) (Bryozoa, Cheilostomata): A forgotten taxon redescribed from Eastern Mediterranean material. *Cahiers de Biologie Marine*, *55*, 91–99.

Harmelin, J.-G. (2017). Bryozoan facies in the coralligenous community: Two assemblages with contrasting features at Port-Cros Archipelago (Port-Cros National Park, France, Mediterranean). *Scientific Reports of the Port-Cros National Park*, *31*, 105–123.

Harmelin, J.-G., Bitar, G., & Zibrowius, H. (2009). Smittinidae (Bryozoa, Cheilostomata) from coastal habitats of Lebanon (Mediterranean Sea), including new and non-indigenous species. *Zoosystema*, *31*(1), 163–187.

Harmelin, J.-G., Bitar, G., & Zibrowius, H. (2016). High xenodiversity versus low native diversity in the south-eastern Mediterranean: Bryozoans from the coastal zone of Lebanon. *Mediterranean Marine Science*, *17*(2), 417–439.

Harmelin, J.-G., & d'Hondt, J.-L. (1993). Transfers of bryozoan species between the Atlantic Ocean and the Mediterranean Sea via the Strait of Gibraltar. *Oceanologica Acta*, *16*(1), 63–72.

Harmelin, J.-G., Ostrovsky, A. N., Caceres-Chamizo, J. P., & Sanner, J. (2011). Bryodiversity in the tropics: Taxonomy of *Microporella* species (Bryozoa, Cheilostomata) with personate maternal zooids from Indian Ocean, Red Sea and southeast Mediterranean. *Zootaxa*, *2798*, 1–30.

Harmelin, J.-G., Vieira, L. M., Ostrovsky, A. N., Cáceres-Chamizo, J. P., & Sanner, J. (2012). *Scorpiodinipora costulata* (Canu & Bassler, 1929) (Bryozoa, Cheilostomata), a taxonomic and biogeographic dilemma: Complex of cryptic species or human-mediated cosmopolitan colonizer? *Zoosystema*, *34*(1), 123–138.

Hastings, A. B. (1927). Zoological results of the Cambridge Expedition to the Suez Canal 1924. Part 20. Report on the Polyzoa. *Transactions of the Zoological Society of London*, *22*, 331–354.

Hastings, A. B. (1941). The British species of Scruparia (Polyzoa). *Annals and Magazine of Natural History*, *(11)7*, 465–472.

Hastings, A. B. (1943). Polyzoa (Bryozoa) I. Scrupocellariidae, Epistomiidae, Farciminariidae, Bicellariellidae, Aeteidae, Scrupariidae. *Discovery Reports*, *22*, 301–510.

Hayward, P. J. (1976). The marine fauna and flora of the Isles of Scilly: Bryozoa II. *Journal of Natural History*, *10*, 319–330.

Hayward, P. J. (1978). Bryozoa from the west European continental slope. *Journal of Zoology, London*, *184*, 207–224.

Hayward, P. J. (1979). Deep water Bryozoa from the coasts of Spain and Portugal. *Cahiers de Biologie Marine*, *20*, 59–75.

Hayward, P. J. (1982). A new species of bryozoan epizooite from South Africa. *Journal of Natural History*, *16*, 769–773.

Hayward, P. J. (1988a). Mauritian cheilostome Bryozoa. *Journal of Zoology, London*, *215*, 269–356.

Hayward, P. J. (1988b). The Recent species of Adeonella (Bryozoa: Cheilostomata) including descriptions of fifteen new species. *Zoological Journal of the Linnean Society*, *94*, 111–191.

Hayward, P. J. (1992). Some Antarctic and sub-Antarctic species of Celleporidae (Bryozoa, Cheilostomata). *Journal of Zoology*, *226*(2), 283–310.

Hayward, P. J., & Parker, S. A. (1994). Notes on some species of Parasmittina Osburn, 1952 (Bryozoa: Cheilostomatida. *Zoological Journal of the Linnean Society*, *110*, 53–75.

Hayward, P. J., & Ryland, J. S. (1978). Bryozoa from the Bay of Biscay and Western Approaches. *Journal of the Marine Biological Association of the United Kingdom*, *58*, 143–159.

Hayward, P. J., & Ryland, J. S. (1993). Taxonomy of six Antarctic anascan Bryozoa. *Antarctic Science*, *5*(2), 129–136.

Hayward, P. J., & Ryland, J. S. (1995a). Bryozoa from Heron Island, Great Barrier Reef. 2. *Memoirs of the Queensland Museum*, *38*(2), 533–573.

Hayward, P. J., & Ryland, J. S. (1995b). The British species of Schizoporella (Bryozoa: Cheilostomatida). *Journal of Zoology, London*, *237*, 37–47.

Hayward, P. J., & Ryland, J. S. (1998). *Cheilostomatous Bryozoa. Part 1. Aeteoidea - Cribrilinoidea.* Shrewsbury: Field Studies Council.

Hayward, P. J., & Thorpe, J. P. (1988). Species of Chaperiopsis (Bryozoa, Cheilostomata) collected by Discovery Investigations. *Journal of Natural History*, *22*, 45–69.

Hayward, P. J., & Thorpe, J. P. (1989). Membraniporoidea, Microporoidea and Cellarioidea (Bryozoa, Cheilostomata) collected by Discovery Investigations. *Journal of Natural History*, *23*, 913–959.

Hayward, P. J., & Thorpe, J. P. (1995). Some British species of Schizomavella (Bryozoa: Cheilostomatida). *Journal of Zoology, London*, *235*, 661–676.

Hayward, P. J., & Winston, J. E. (2011). Bryozoa collected by the United States Antarctic Research Program: New taxa and new records. *Journal of Natural History*, *45*(37-38), 2259–2338.

Heindl, H., Thiel, V., Wiese, J., & Imhoff, J. F. (2012). Bacterial isolates from the bryozoan *Membranipora membranacea*: Influence of culture media on isolation and antimicrobial activity. *International Microbiology*, *15*, 17–32. https://doi.org/10.2436/20.1501.01.155

Helmkampf, M., Bruchhaus, I., & Hausdorf, B. (2008). Multigene analysis of lophophorate and chaetognath phylogenetic relationships. *Molecular Phylogenetics and Evolution*, *46*(1), 206–214.

Herrera-Cubilla, A., Dick, M. H., Sanner, J., & Jackson, J. B. C. (2006). Neogene Cupuladriidae of tropical America. I: Taxonomy of Recent *Cupuladria* from opposite sides of the Isthmus of Panama. *Journal of Paleontology*, *80*(2), 245–263.

Herrera-Cubilla, A., Dick, M. H., Sanner, J., & Jackson, J. B. C. (2008). Neogene Cupuladriidae of tropical America. II: Taxonomy of Recent *Discoporella* from opposite sides of the Isthmus of Panama. *Journal of Paleontology*, *82*(2), 279–298.

Hincks, T. (1860). Descriptions of new Polyzoa from Ireland. *Quarterly Journal of Microscopical Science*, *8*(32), 275–280.

Hincks, T. (1882). Report on the Polyzoa of the Queen Charlotte Islands. *Annals and Magazine of Natural History*, *10*, 459–471.

Hincks, T. (1886). The Polyzoa of the Adriatic, I. *Annals and Magazine of Natural History*, *(5)17*, 254–271.

Hirose, M. (2011). Orientation and righting behavior of the sand-dwelling bryozoan *Conescharellina catella*. *Invertebrate Biology*, *130*(3), 282–290. https://doi.org/10.1111/j.1744-7410.2011.00237.x

Hirose, M. (2012). Revision of the genus *Buchneria* (Bryozoa, Cheilostomata) from Japan. *Zookeys*, *241*, 1–19.

Hirose, M. (2017). Diversity of freshwater and marine bryozoans in Japan. In M. Motokawa & H. Kajihara (Eds.), *Species Diversity of Animals in Japan* (pp. 629–649). Springer.

Hirose, M., Mawatari, S. F., & Scholz, J. (2012). Distribution and diversity of erect bryozoan assemblages along the Pacific coast of Japan. In A. Ernst, P. Schäfer, & J. Scholz (Eds.), *Bryozoan Studies 2010* (Vol. 143, pp. 121–136). Berlin: Springer.

Hu, Y.-m., & Wang, F.-z. (1984). A new species of Ectoprocta from the Antarctic. *Acta Zootaxonomica Sinica*, *9*, 241–243.

Huang, Z., Li, C., & Liu, X. (1990). The bryozoan foulers of Hong Kong and neighbouring waters. In B. Morton (Ed.), *The marine flora and fauna of Hong Kong and southern China, 1986.: Vol. 2. Taxonomy and ecology* (pp. 737–765). Hong Kong: Hong Kong University Press.

Hughes, D. J., & Jackson, J. B. C. (1992). Distribution and abundance of cheilostome bryozoans on the Caribbean reefs of central Panama. *Bulletin of Marine Science*, *51*(3), 443–465.

Hughes, R. L., & Woollacott, R. M. (1978). Ultrastructure of potential photoreceptor organs in the larva of *Scrupocellaria bertholetti* (Bryozoa). *Zoomorphologie*, *91*, 225–234.

Hughes, R. L., & Woollacott, R. M. (1980). Photoreceptors of bryozoan larvae (Cheilostomata, Cellularioidea). *Zoologica Scripta*, *9*, 129–138.

Hughes, R. N., & Wright, P. J. (2014). Self-fertilisation in the Celleporella angusta clade and a description of Celleporella osiani sp. Nov. *Studi Trentini Di Scienze Naturali*, *94*, 119–124.

Hunter, E., Okano, K., Tomono, Y., & Fusetani, N. (1998). Functional partitioning of energy reserves by larvae of the marine bryozoan *Bugula neritina* (L.). *Journal of Experimental Biology*, *201*, 2857–2865. Retrieved from://000077039700007

Igić, L. (n.d.). *Some species of bryozoa from the Adriatic sea and from freshwaters, which are of special importance for fouling complex.* Center for Investigation of Sea "Ruđer Bošković", Rovinj, Croatia.

Ito, M., Onishi, T., & Dick, M. H. (2015). Cribrilina mutabilis n. Sp., an eelgrass-associated bryozoan (Gymnolaemata: Cheilostomata) with large variation in zooid morphology related to life history. *Zoological Science*, *32*, 485–497.

Iyengar, E. V., & Harvell, C. D. (2002). Specificity of cues inducing defensive spines in the bryozoan Membranipora membranacea. *Marine Ecology-Progress Series*, *225*, 205–218.

Javed, M., & Tirmizi, N. M. (1993). Four species of Bugula (Bryozoa: Bugulidae) new to the Pakistan coast (northern Arabian Sea). *Pakistan Journal of Zoology*, *25*(4), 285–288.

Jelly, E. C. (1889). *A synonymic catalogue of the Recent marine Bryozoa including fossil synonyms.* London: Dulau & Company.

Johnson, C. H. (2010). Effects of Selfing on Offspring Survival and Reproduction in a Colonial Simultaneous Hermaphrodite (Bugula Stolonifera, Bryozoa). *The Biological Bulletin*, *219*(1), 27–37.

Johnson, C. H., Winston, J. E., & Woollacott, R. M. (2012). Western Atlantic introduction and persistence of the marine bryozoan Tricellaria inopinata. *Aquatic Invasions*, *7*(3), 295–303.

Jr, G. R. M., & McKinney, F. K. (2000). A Theoretical Morphologic Analysis of Convergently Evolved Erect Helical Colony Form in the Bryozoa. *Paleobiology*, *26*(4), 556–577. https://doi.org/10.1666/0094-8373(2000)026<0556:atmaoc>2.0.co;2

Kaiser, S., Barnes, D. K. A., Linse, K., & Brandt, A. (2008). Epibenthic macrofauna associated with the shelf and slope of a young and isolated Southern Ocean island. *Antarctic Science*, *20*(3), 281–290.

Karagodina, N. P., Vishnyakov, A. E., Kotenko, O. N., Maltseva, A. L., & Ostrovsky, A. N. (2018). Ultrastructural evidence for nutritional relationships between a marine colonial invertebrate (Bryozoa) and its bacterial symbionts. *Symbiosis*, 1–10.

Kasaei, S. M., Nasrolahi, A., Abtahi, B., & Taylor, P. D. (2017). Bryozoa of the southern Caspian Sea, Iranian coast. *Check List*, *13*(4), 305–313. https://doi.org/10.15560/13.4.305

Kaselowsky, J., Scholz, J., & Levit, G. S. (2002). The biological potential of encrusting bryozoans. *Senckenbergiana Lethaea*, *82*(1), 181–192.

Kaselowsky, J., Scholz, J., Mawatari, S. F., Probert, P. K., Gerdes, G., Kadagies, N., & Hillmer, G. (2005). Bryozoans and microbial communities of cool-temperate to subtropical latitudes - paleoecological implications. I. Growth morphologies of shallow-water bryozoans settling on bivalve shells (Japan and New Zealand). *Facies*, *50*, 349–361.

Kelmo, F., Attrill, M. J., Gomes, R. C. T., & Jones, M. B. (2004). El Niño induced local extinction of coral reef bryozoan species from Northern Bahia, Brazil. *Biological Conservation*, *118*, 609–617.

Kelso, A., & Wyse Jackson, P. N. (2012). Invasive bryozoans in Ireland: First record of Watersipora subtorquata (d'Orbigny, 1852) and an extension of the range of *Tricellaria inopinata* d'Hondt and *Occhipinti ambrogi*, 1985. *BioInvasions Records*, *1*(3), 209–214.

Kelso, A., & Wyse Jackson, P. N. (2014). Albert Russell Nichols (1859-1933) and bryozoan research in Ireland during the late 19th and early 20th centuries. *International Bryozoology Association, Dublin*, 93–106.

Key, M. M., Hollenbeck, P. M., O'Dea, A., & Patterson, W. P. (2013). Stable isotope profiling in modern marine bryozoan colonies across the Isthmus of Panama. *Bulletin of Marine Science*, *89*(4), 837–856.

Kind, B., De Blauwe, H., Faasse, M., & Kuhlenkamp, R. (2015). *Schizobrachiella verrilli* (Bryozoa, Cheilostomata) new to Europe. *Marine Biodiversity Records*, *8*.

Kind, B., & Kuhlenkamp, R. (2018). Discovery of the non-indigenous bryozoan Smittoidea prolifica Osburn, 1952 near Helgoland: First record in 2011 for the German North Sea. *Marine Biodiversity*, *48*(2), 1237–1240.

Kobluk, D. R., Cuffey, R. J., Fonda, S. S., & Lysenko, M. A. (1988). Cryptic Bryozoa, leeward fringing reef of Bonaire, Netherlands Antilles, and their paleoecological application. *Journal of Paleontology*, *62*(3), 427–439.

Koçak, F. (2007). A new alien bryozoan *Celleporaria brunnea* (Hincks, 1884) in the Aegean Sea (eastern Mediterranean). *Scientia Marina*, *71*(1), 191–195.

Koçak, F., Baldu, A., i, & Benli, H. A. (2002). Epiphytic bryozoan community of *Posidonia oceanica* (L.) Delile meadow in the northern Cyprus (Eastern Mediterranean). *Indian Journal of Marine Sciences*, *31*(3), 235–238.

Koçak, F., & Önen, A. (2014). Checklist of Bryozoa on the coasts of Turkey. *Turkish Journal of Zoology*, *38*, 880–891.

Kozloff, E. N., & Price, L. H. (1996). *Marine Invertebrates of the Pacific Northwest* (Rep Sub). Seattle: Univ of Washington Press.

Krzeminska, M., Sicinski, J., & Kuklinski, P. (2018). Biodiversity and biogeographic affiliation of Bryozoa from King George Island (Antarctica). *Systematics and Biodiversity*, *16*(6), 576–586.

Kubota, K., & Mawatari, S. F. (1985). A systematic study of cheilostomatous bryozoans from Oshoro Bay, Hokkaido : 1. Anasca. *Environmental Science, Hokkaido*, *8*(1), 75–91.

Kubota, K., & Mawatari, S. F. (1986). A systematic study of ceilostomatous bryozoans from Oshoro Bay, Hokkaido. 2. Ascophora. *Environmental Science, Hokkaido*, *8*(2), 195–208.

Kuklinski, P. (2002). Fauna of Bryozoa from Kongsfjorden, West Spitsbergen. *Polish Polar Research*, *23*(2), 193–206.

Kuklinski, P. (2009). Ecology of stone-encrusting organisms in the Greenland Sea - a review. *Polar Research*, *28*, 222–237.

Kuklinski, P., & Bader, B. (2007). Comparison of bryozoan assemblages from two contrasting Arctic shelf regions. *Estuarine, Coastal and Shelf Science*, *73*(3-4), 835–843. https://doi.org/10.1016/j.ecss.2007.03.024

Kuklinski, P., & Barnes, D. K. A. (2009). A new genus and three new species of Antarctic cheilostome Bryozoa. *Polar Biology*, *32*, 1251–1259.

Kuklinski, P., Grischenko, A. V., & Jewett, S. C. (2015). Two new species of the cheilostome bryozoan Cheilopora from the Aleutian Islands. *Zootaxa*, *3963*(3), 434–442.

Kuklinski, P., & Hayward, P. J. (2004). Two new species of cheilostome Bryozoa from Svalbard. *Sarsia*, *89*, 79–84.

Kuklinski, P., & Taylor, P. D. (2006). A new genus and some cryptic species of Arctic and boreal calloporid cheilostome bryozoans. *Journal of the Marine Biological Association*, *86*, 1035–1046.

Kuklinski, P., & Taylor, P. D. (2007). Are bryozoans adapted for living in the Arctic? *Proceedings of the 14th International Bryozoology Association Conference, Boone, North Carolina, July 1-8, 2007, Virginia Museum of Natural History. Special Publication No. 15*, 101–110.

Kuklinski, P., & Taylor, P. D. (2008). Arctic species of the cheilostome bryozoan Microporella, with a redescription of the type species. *Journal of Natural History*, *42*(27-28), 1893–1906.

Kuklinski, P., Taylor, P. D., & Denisenko, N. V. (2007). Arctic cheilostome bryozoan species of the genus *Escharoides*. *Journal of Natural History*, *41*(1-4), 219–228.

Kutyumov, V. A., Maltseva, A. L., Kotenko, O. N., & Ostrovsky, A. N. (2016). Functional differentiation in bryozoan colonies: A proteomic analysis. *Cell and Tissue Biology*, *10*(2), 152–159. https://doi.org/10.1134/S1990519X16020073

Lacourt, A. W. (1949). Bryozoa of the Netherlands. *Archives Nederlandaises de Zoologie*, *8*, 289–321.

Landsborough, D. (1852). *A Popular History of British Zoophytes, or Corallines.* London: Reeve and Co.

Lauer, A. (2016). *Watersipora subtorquata* and the possible role of its associated microbes: An attempt to explain the extraordinary invasion success of this marine bryozoan species. In C. J. Hurst (Ed.), *The Mechanistic Benefits of Microbial Symbionts* (Vol. 2, pp. 239–268). https://doi.org/10.1007/978-3-319-28068-4_9

Lim, G. E., & Haygood, M. G. (2004). "Candidatus *Endobugula glebosa*," a specific bacterial symbiont of the marine bryozoan *Bugula* simplex. *Applied and Environmental Microbiology*, *70*(8), 4921–4929.

Linneman, J., Paulus, D., Lim-Fong, G. E., & Lopanik, N. B. (2014). Latitudinal variation of a defensive symbiosis in the *Bugula neritina* (Bryozoa) sibling species complex. *PLoS One*, *9*(10). https://doi.org/10.1371/journal.pone.0108783

Linse, K., Barnes, D. K. A., & Enderlein, P. (2006). Body size and growth of benthic invertebrates along an Antarctic latitudinal gradient. *Deep-Sea Research II*, *53*, 921–931.

Liu, X.-X. (1992). On the genus Membranipora (Anasca: Cheilostomata: Bryozoa) from south Chinese seas. *Raffles Bulletin of Zoology*, *40*(1), 103–144.

Liuzzi, M. G., López-Gappa, J., & Salgado, L. (2018). Bryozoa from the continental shelf off Tierra del Fuego (Argentina): Species richness, colonial growth-forms, and their relationship with water depth. *Estuarine, Coastal and Shelf Science*, *214*, 48–56. https://doi.org/10.1016/j.ecss.2018.09.014

Livingstone, A. A. (1924). Studies on Australian Bryozoa. No.1. *Records of the Australian Museum*, *14*, 189–212.

Livingstone, A. A. (1928). The Bryozoa. Supplementary Report. *Scientific Reports of the Australian Antarctic Expedition 1911-1914, Series C, Zoology and Botany*, *9*(1), 1–93.

Lodola, A., Savini, D., & Occhipinti-Ambrogi, A. (2012). First record of *Tricellaria inopinata* (Bryozoa: Candidae) in the harbours of La Spezia and Olbia, Western Mediterranean Sea (Italy). *Marine Biodiversity Records*, *5*.

Lomas, J. (1886). Report on the Polyzoa of the L. M. B. C. District. In *Report upon the fauna of Liverpool Bay and the neighboring seas* (Vol. 1, pp. 161–200). London: Liverpool Marine Biology Committee, University College Liverpool.

Lombardi, C., Cocito, S., Gambi, M. C., Cisterna, B., Flach, F., Taylor, P. D., … Cusack, M. (2011). Effects of ocean acidification on growth, organic tissue and protein profile of the Mediterranean bryozoan Myriapora truncata. *Aquatic Biology*, *13*(3), 251–262.

Lombardi, C., Gambi, M. C., Vasapollo, C., Taylor, P. D., & Cocito, S. (2011). Skeletal alterations and polymorphism in a Mediterranean bryozoan at natural CO2 vents.

*Zoomorphology*, *130*, 135–145.

Lombardi, C., Rodolfo-Metalpa, R., Cocito, S., Gambi, M. C., & Taylor, P. D. (2011). Structural and geochemical alterations in the Mg calcite bryozoan LiverpoolMyriapora truncata under elevated seawater pCO2 simulating ocean acidification. *Marine Ecology*, *32*(2), 211–221.

Lombardi, C., Taylor, P. D., & Cocito, S. (2014). Bryozoan constructions in a changing mediterranean sea. In *The Mediterranean Sea* (pp. 373–384). https://doi.org/10.1007/978-94-007-6704-1_21

Lombardi, C., Taylor, P. D., Cocito, S., Bertolini, C., & Calosi, P. (2017). Low pH conditions impair module capacity to regenerate in a calcified colonial invertebrate, the bryozoan *Cryptosula pallasiana*. *Marine Environmental Research*, *125*, 110–117. https://doi.org/10.1016/j.marenvres.2017.02.002

Long, E. R., & Rucker, J. B. (1960). A comparative study of cheilostome Bryozoa at Yokosuka, Maizuru, and Sasebo, Japan. *Pacific Science*, *23*, 56–69.

Long, E. R., & Rucker, J. B. (1970). Offshore marine cheilostome Bryozoa from Fort Lauderdale, Florida. *Marine Biology*, *6*, 18–25.

Lopanik, N., Lindquist, N., & Targett, N. (2004). Potent cytotoxins produced by a microbial symbiont protect host larvae from predation. *Oecologia*, *139*, 131–139.

Loxton, J., Kuklinski, P., Mair, J. M., Spencer Jones, M. E., & Porter, J. S. (2012). Patterns of magnesium-calcite distribution in the skeleton of some polar bryozoan species. In A. Ernst, P. Schäfer, & J. Scholz (Eds.), *Bryozoan Studies 2010* (Vol. 143, pp. 169–185). Berlin: Springer.

Loxton, J., Wood, C. A., Bishop, J. D. D., Porter, J. S., Jones, M. S., & Nall, C. R. (2017). Distribution of the invasive bryozoan *Schizoporella japonica* in Great Britain and Ireland and a review of its European distribution. *Biological Invasions*, *19*(8), 2225–2235.

López de la Cuadra, C. M., & García-Gómez, J. C. (1991). A new species of *Hemicyclopora* (Bryozoa, Cheilostomata) from the southern coast of Spain. In F. P. Bigey (Ed.), *Bryozoaires Actuels et Fossiles: Bryozoa Living and Fossil* (pp. 213–218). Nantes: Bulletin de la Société Sciences Naturelles de l'Ouest de la France, Mémoire HS 1.

López de la Cuadra, C. M., & García-Gómez, J. C. (1993). Little-known Atlantic cheilostome bryozoans at the entrance to the Mediterranean. *Journal of Natural History*, *27*(2), 457–469.

López de la Cuadra, C. M., & García-Gómez, J. C. (1994). Bryozoa Cheilostomata: The genus *Amphiblestrum* in the Western Mediterranean and the first of Atlantic waters. *Journal of Natural History*, *28*, 683–693.

López de la Cuadra, C. M., & García-Gómez, J. C. (1997). Studies on Recent Macroporidae (Bryozoa: Cheilostomatida), with new taxa and ontogeny of the ovicells. *Journal of Zoology, London*, *242*, 605–621.

López de la Cuadra, C. M., & García-Gómez, J. C. (2000). The cheilostomate Bryozoa (Bryozoa: Cheilostomatida) collected by the Spanish 'Antártida 8611' Expedition to the Scotia Arc and South Shetland Islands. *Journal of Natural History*, *34*, 755–772.

López de la Cuadra, C. M., & García-Gómez, J. C. (2001). New and little known ascophoran bryozoans from the Western Mediterranean, collected by 'Fauna Ibérica' expeditions. *Journal of Natural History*, *35*, 1717–1732.

López-Fé, C. M. (2006). Some bathyal cheilostome Bryozoa (Bryozoa, Cheilostomata) from the Canary Islands (Spain, Eastern Atlantic), with descriptions of three new species, a new genus, and a new family. *Journal of Natural History*, *40*(29-31), 1801–1812. Retrieved from http://dx.doi.org/10.1080/00222930601043763

López Gappa, J. J. (1986). A new bryozoan genus from the Weddell Sea, Antarctica. *Polar Biology*, *6*, 103–105.

López Gappa, J. J. (1989). Overgrowth competition in an assemblage of encrusting bryozoans settled on artificial substrata. *Marine Ecology Progress Series*, *51*, 121–130.

López Gappa, J. J., & Landoni, N. A. (2009). Space utilisation patterns of bryozoans on the Patagonian scallop Psychrochlamys patagonica. *Scientia Marina*, *73*(1), 161–171.

López Gappa, J. J., & Liuzzi, M. G., i. (2009). A new Antarctic *Osthimosia* (Bryozoa, Cheilostomata, Celleporidae) with dimorphic zooids. *Polar Biology*, *32*, 47–51.

López-Gappa, J., & Liuzzi, M. G. (2018). Recent discovery of non-indigenous bryozoans in the fouling assemblage of Quequén Harbour (Argentina, Southwest Atlantic). *Marine Biodiversity*, *48*(2), 1159–1167.

López-Gappa, J., Pérez, L. M., & Griffin, M. (2017). First record of a fossil selenariid bryozoan in South America. *Alcheringa*, *41*(3), 365–368.

Lörz, A.-N., Myers, A., & Gordon, D. P. (2014). An inquiline deep-water bryozoan/amphipod association from New Zealand, including the description of a new genus and species of Chevaliidae. *European Journal of Taxonomy*, *72*, 1–17.

Lyke, E. B., Reed, C. G., & Woollacott, R. M. (1983). Origin of the cystid epidermis during the metamorphosis of three species of gymnolaemate bryozoans. *Zoomorphology*, *102*(2), 99–110.

MacGillivray, P. H. (1887a). A catalogue of the marine Polyzoa of Victoria. *Transactions and Proceedings of the Royal Society of Victoria*, *23*, 187–224.

MacGillivray, P. H. (1887b). Descriptions of new or little-known Polyzoa, Part 12. *Transactions and Proceedings of the Royal Society of Victoria*, *23*, 179–186.

Mackie, J. A., Darling, J. A., & Geller, J. B. (2012). Ecology of cryptic invasions: Latitudinal segregation among Watersipora (Bryozoa) species. *Scientific Reports*, *2*(871), 1–10. https://doi.org/DOI 10.1038/srep00871

Madurell, T., Zabala, M., Dominguez-Carrió, C., & Gili, J. M. (2013). Bryozoan faunal composition and community structure from the continental shelf off Cap de Creus (Northwestern Mediterranean). *Journal of Sea Research*, *83*, 123–136.

Maltseva, A. L., Kotenko, O. N., Shabalin, K. A., Shavarda, A. L., Winson, M. K., & Ostrovsky, A. N. (2014). Novel brominated fungicidal alkaloid isolated from the marine bryozoan *Chartella membranaceatruncata* (Smitt, 1868). *Studi Trent. Sci. Nat*, *94*, 163–168.

Maplestone, C. M. (1900). Further descriptions of the Tertiary Polyzoa of Victoria. 4. *Proceedings of the Royal Society of Victoria (New Series)*, *13*, 1–9.

Maplestone, C. M. (1904). Tabulated list of the fossil cheilostomatous Polyzoa in the Victorian Tertiary deposits. *Proceedings of the Royal Society of Victoria (New Series)*, *17*, 182–219.

Maplestone, C. M. (1909). Polyzoa from the Gilbert Islands. *Proceedings of the Royal Society of Victoria (New Series)*, *21*, 410–419.

Maplestone, C. M. (1910). On a new species of Cellepora from the south Australian coast. *Proceedings of the Royal Society of Victoria (New Series)*, *23*, 39–41.

Marchini, A., Cunha, M. R., & Occhipinti Ambrogi, A. (2007). First observations on bryozoans and entoprocts in the Ria de Aveiro (NW Portugal) including the first record of the Pacific invasive cheilostome Tricellaria inopinata. *Marine Ecology*, *28*(Suppl. 1), 154–160.

Markert, A., Matsuyama, K., Rohde, S., Schupp, P., & Wehrmann, A. (2016). First record of the non-native Pacific bryozoan *Smittoidea prolifica* Osburn, 1952 at the German North Sea coast. *Marine Biodiversity*, *46*(3), 717–723.

Marques, A. C., Kloh, A. dos S., Migotto, A. E., Cabral, A. C., Rigo, A. P. R., Bettim, A. L., ... Kremer, L. P. (2013). Rapid assessment survey for exotic benthic species in the São Sebastião Channel, Brazil. *Latin American Journal of Aquatic Research*, *41*(2), 398–407.

Marshall, D. J., Bolton, T. F., & Keough, M. J. (2003). Offspring size affects the post-metamorphic performance of a colonial marine invertebrate. *Ecology*, *84*(12), 3131–3137.

Mathew, M., & Lopanik, N. B. (2014). Host Differentially Expressed Genes During Association With Its Defensive Endosymbiont. *The Biological Bulletin*, *226*(2), 152–163.

Matsuyama, K., Janssen, A., Martínez Arbizu, P., Martha, S. O., & Freiwald, A. (2014). Bryozoans from RV Sonne deep-sea cruises SO 167 'Louisville' and SO 205 'Mangan'. *Zootaxa*, *3856*(1), 100–116.

Matsuyama, K., Titschack, J., Baum, D., & Freiwald, A. (2015). Two new species of erect Bryozoa (Gymnolaemata: Cheilostomata) and the application of non-destructive imaging methods for quantitative taxonomy. *Zootaxa*, *4020*(1), 81–100.

Maturo, F. J. S., & Schopf, T. J. M. (1968). Ectoproct and entoproct type material: Reexamination of species from New England and Bermuda named by A.E. Verrill, J.W. Dawson and E. Desor. *Postilla*, *120*, 1–95.

Mawatari, S. F., & Suwa, T. (1998). Two new species of Japanese *Microporella* (Bryozoa, Cheilostomatida) in the Döderlein Collection, Musée Zoologique, Strasbourg. *Cahiers de Biologie Marine*, *39*(1), 1–7.

McCann, L. (2019). Bryozoa (Cheilostomata, Ctenostomata, and Cyclostomata) in Galapagos Island Fouling Communities. *Aquatic Invasions*, *14*(1), 85–131. https://doi.org/10.3391/ai.2019.14.1.04

McCann, L. D., Gray Hitchcock, N., Winston, J. E., & Ruiz, G. M. (2007). Non-native bryozoans in coastal embayments of the southern United States: New records for the

Western Atlantic. *Bulletin of Marine Science*, *80*(2), 319–342.

McCuller, M. I., & Carlton, J. T. (2018). Transoceanic rafting of Bryozoa (Cyclostomata, Cheilostomata, and Ctenostomata) across the North Pacific Ocean on Japanese tsunami marine debris. *Aquatic Invasions*, *13*(1), 137–162.

McCuller, M. I., Carlton, J. T., & Geller, J. B. (2018). *Bugula tsunamiensis* n. Sp. (Bryozoa, Cheilostomata, Bugulidae) from Japanese tsunami marine debris landed in the Hawaiian Archipelago and the Pacific Coast of the USA. *Aquatic Invasions*, *13*(1), 163–172.

McGhee, G., Jr. (2000). A theoretical morphologic analysis of convergently evolved erect helical colony form in the Bryozoa. *Paleobiology*, *26*(4), 556–577. https://doi.org/10.1666/0094-8373(2000)026<0556:ATMAOC>2.0.CO;2

McKinney, F. K., & Jaklin, A. (2000). Spatial niche partitioning in the *Cellaria* meadow epibiont association, northern Adriatic Sea. *Cahiers de Biologie Marine*, *41*(1), 1–17.

McKinney, F. K., & McKinney, M. J. (1993). Larval behaviour and choice of settlement site: Correlation with environmental distribution pattern in an erect bryozoan. *Facies*, *29*, 119–132.

Menon, N. R. (1972). Species of the genus Scrupocellaria Van Beneden (Bryozoa, Anasca) from Indian waters. *Internationale Revue Der Gesamten Hydrobiologie Und Hydrographie*, *57*(6), 913–931.

Menon, N. R., & Nair, N. B. (1982). Indian species of Malacostega (Polyzoa, Ectoprocta). *The Marine Biological Association of India*, *17*, 553–579.

Metcalfe, K., Gordon, D. P., & Hayward, E. (2007). An amphibious bryozoan from living mangrove leaves - *Amphibiobeania* new genus (Beaniidae). *Zoological Science*, *24*, 563–570.

Micael, J., Jardim, N., Núñez, C., Occhipinti Ambrogi, A., & Costa, A. C. (2016). Some Bryozoa species recently introduced into the Azores: Reproductive strategies as a proxy for further spread. *Helgoland Marine Research*, *70*(7). https://doi.org/10.1186/s10152-016-0458-7

Micael, J., Marina, J. G., Costa, A. C., & Occhipinti-Ambrogi, A. (2014). The non-indigenous *Schizoporella errata* (Bryozoa: Cheilostomatida) introduced into the Azores Archipelago. *Marine Biodiversity Records*, *7*, e133.

Micael, J., Tempera, F., Berning, B., López-Fé, C. M., Occhipinti-Ambrogi, A., & Costa, A. C. (2017). Shallow-water bryozoans from the Azores (central North Atlantic): Native vs. Non-indigenous species, and a method to evaluate taxonomic uncertainty. *Marine Biodiversity*, 1–12. https://doi.org/10.1007/s12526-017-0833-x

Micael, J., Tempera, F., Berning, B., López-Fé, C. M., Occhipinti-Ambrogi, A., & Costa, A. C. (2019). Shallow-water bryozoans from the Azores (central North Atlantic): Native vs. Non-indigenous species, and a method to evaluate taxonomic uncertainty. *Marine Biodiversity*, *49*(1), 469–480. https://doi.org/10.1007/s12526-017-0833-x

Min, B. S., Seo, J. E., Grischenko, A. V., & Gordon, D. P. (2017). Intertidal Bryozoa from Korea—new additions to the fauna and a new genus of Bitectiporidae (Cheilostomata) from Baengnyeong Island, Yellow Sea. *Zootaxa*, *4226*(4), 451–470.

Min, B. S., Seo, J. E., Grischenko, A. V., Lee, S.-K., & Gordon, D. P. (2017). Systematics of some calloporid and lacernid Cheilostomata (Bryozoa) from coastal South Korean waters, with the description of new taxa. *Zootaxa*, *4226*(4), 471–486.

Miranda, A. A., Almeida, A. C. S., & Vieira, L. M. (2018). Non-native marine bryozoans (Bryozoa: Gymnolaemata) in Brazilian waters: Assessment, dispersal and impacts. *Marine Pollution Bulletin*, *130*, 184–191. https://doi.org/10.1016/j.marpolbul.2018.03.023

Moosbrugger, M., Schwaha, T., Walzl, M. G., Obst, M., & Ostrovsky, A. N. (2012). The placental analogue and the pattern of sexual reproduction in the cheilostome bryozoan *Bicellariella ciliata* (Gymnolaemata). *Frontiers in Zoology*, *9*(1), 1–20.

Morgado, E. H., & Tanaka, M. O. (2001). The macrofauna associated with the bryozoan *Schizoporella errata* (Waters) in southeastern Brazil. *Scientia Marina*, *65*(3), 173–181.

Morris, B., & Mogelberg, D. (1973). Identification manual to the pelagic Sargassum fauna. *Special Publication of the Bermuda Biological Station for Research*, *11*, 1–63.

Morris, R. H., Abbott, D. P., & Haderlie, E. C. (1980). Bryozoa and Entoprocta: The Moss Animals. In *Intertidal invertebrates of California* (pp. 91–107). Stanford, Calif.: Stanford University Press.

Morrison, M., Jones, E. G., Consalvey, M., & Berkenbusch, K. (2014). *Linking marine fisheries species to biogenic habitats in New Zealand: A review and synthesis of knowledge.* New Zealand: Ministry for Primary Industries.

Moyano, H. I. (1982). Magellanic Bryozoa: Some ecological and zoogeographical aspects. *Marine Biology*, *67*, 81–96.

Moyano, H. I. (1999). Magellan Bryozoa: A review of the diversity and of the Subantarctic and Antarctic zoogeographical links. *Scientia Marina*, *63*, 219–226.

Moyano, H. I. (2011). On *Xenoflustra voigti* n. Gen., n. Sp. (Bryozoa, Cheilostomatida, Buguloidea), a new flustrine bryozoan from the south western Atlantic Ocean. *Anales Instituto Patagonia*, *39*(2), 67–71.

Moyano, H. I., & Gordon, D. P. (1980). New species of Hippothoidae (Bryozoa) from Chile, Antarctica and New Zealand. *Journal of the Royal Society of New Zealand*, *10*(1), 75–95.

Moyano G., H. I. (1986). Bryozoa marinos chilenos VI. Cheilostomata Hippothoidae: South eastern Pacific species. *Boletín de La Real Sociedad Española de Historia Natural*, *57*, 89–135.

Moyano G., H. I. (2002). Towards a general history of the south eastern Pacific bryozoans. In P. N. Wyse Jackson & M. E. Spencer Jones (Eds.), *Annals of Bryozoology: Aspects of the history of research on bryozoans* (pp. 171–183). Dublin: International Bryozoology Association.

Muravchik, M., & Griffin, M. (2004). Bryozoans from the Paraná Formation (Miocene), in Entre Ríos province, Argentina. *Ameghiniana*, *41*(1), 3–12.

Nall, C. R., Guerin, A. J., & Cook, E. J. (2015). Rapid assessment of marine non-native species in northern Scotland and a synthesis of existing Scottish records. *Aquatic Invasions*,

*10*(1), 107–121.

Nasto, I., Cardone, F., Mastrototaro, F., Panetta, P., Rosso, A., Sanfilippo, R., … Tursi, A. (2018). Benthic invertebrates associated with subfossil cold-water coral frames and hardgrounds in the Albanian deep waters (Adriatic Sea). *Turkish Journal of Zoology*, *42*(4), 360–371. https://doi.org/10.3906/zoo-1708-44

Nekliudova, U. A., Schwaha, T. F., Kotenko, O. N., Gruber, D., Cyran, N., & Ostrovsky, A. N. (2019). Sexual reproduction of the placental brooder *Celleporella Hyalina* (Bryozoa, Cheilostomata) in the White Sea. *Journal of Morphology*, *280*(2), 278–299. https://doi.org/10.1002/jmor.20943

Nesnidal, M. P., Helmkampf, M., Bruchhaus, I., & Hausdorf, B. (2011). The complete mitochondrial genome of *Flustra foliacea* (Ectoprocta, Cheilostomata)-compositional bias affects phylogenetic analyses of lophotrochozoan relationships. *BMC Genomics*, *12*(1), 572.

Newcomer, K., Marraffini, M. L., & Chang, A. L. (2018). Distribution patterns of the introduced encrusting bryozoan Conopeum chesapeakensis (Osburn 1944; Banta et al. 1995) in an estuarine environment in upper San Francisco Bay. *Journal of Experimental Marine Biology and Ecology*, *504*, 20–31. https://doi.org/10.1016/j.jembe.2018.04.001

Nikulina, E. A. (2007). *Einhornia*, a new genus for electrids formerly classified as the *Electra crustulenta* species group (Bryozoa, Cheilostomata). *Schriften Des Naturwissenschaftlichen Vereins Für Schleswig-Holstein*, *69*, 29–40.

Nikulina, E. A. (2008). Electra scuticifera sp. Nov.: redescription of Electra pilosa from New Zealand as a new species (Bryozoa, Cheilostomata). *Schriften Des Naturwissenschaftlichen Vereins Für Schleswig-Holstein*, *70*, 91–98.

Nikulina, E. A., De Blauwe, H., & Reverter-Gil, O. (2012a). Molecular phylogenetic analysis confirms the species status of Electra verticillata (Ellis and Solander, 1786). In A. Ernst, P. Schäfer, & J. Scholz (Eds.), *Bryozoan Studies 2010* (Vol. 143, pp. 217–236). Berlin: Springer.

Nikulina, E. A., Ostrovsky, A. N., & Claereboudt, M. (2012b). A new species of the genus *Electra* (Bryozoa, Cheilostomata) from southern Oman, Arabian Sea. In A. Ernst, P. Schäfer, & J. Scholz (Eds.), *Bryozoan Studies 2010* (Vol. 143, pp. 203–216). Berlin: Springer.

Nikulina, E. A., & Schäfer, P. (2006). Bryozoans of the Baltic Sea. *Meyniana*, *58*, 75–95.

Nikulina, E. N., Hanel, R., & Schäfer, P. (2007). Cryptic speciation and paraphyly in the cosmopolitan bryozoan Electra pilosa - impact of the Tethys closing on species evolution. *Molecular Phylogenetics and Evolution*, *45*, 765–776.

Nordgaard, O. (1905). Hydrographical and biological investigations in Norwegian fiords. *Bergens Museums Meereskrifter*, *2*, 155–254.

Norman, A. M. (1868). Notes on some rare British Polyzoa, with descriptions of new species. *Quarterly Journal of Microscopical Science*, *8*(32), 212–222.

Norman, A. M. (1903). Notes on the natural history of East Finmark. Polyzoa. *Annals and Magazine of Natural History, Ser. 7*, *12*, 87–128.

Novosel, M. (2005). Bryozoans of the Adriatic Sea. *Denisia*, *16*, 231–246.

Novosel, M., & Pozar-Domac, A. (2001). Checklist of Bryozoa of the eastern Adriatic Sea. *Natura Croatica*, *10*(4), 367–421.

Novosel, M., Pozar-Domac, A., & Pasaric, M. (2004). Diversity and distribution of the Bryozoa along underwater cliffs in the Adriatic Sea with special reference to thermal regime. *Marine Ecology*, *25*(2), 155–170.

Nowak, M., Kuklinski, P., & Sicinski, J. (2014). Small-scale biomass variability of Antarctic bryozoans. *Studi Trentini Di Scienze Naturali*, *94*, 181–188.

O'Dea, A. (2006). Asexual propagation in the marine bryozoan *Cupuladria exfragminis*. *Journal of Experimental Marine Biology and Ecology*, *335*(2), 312–322.

O'Dea, A., Herrera-Cubilla, A., Fortunato, H., & Jackson, J. B. C. (2004). Life history variation in cupuladriid bryozoans from either side of the Isthmus of Panama. *Marine Ecology-Progress Series*, *280*, 145–161.

O'Donoghue, C. H. (1924). The Bryozoa (Polyzoa) collected by the S.S. 'Pickle'. *Report No. 3 of the Fisheries and Marine Biological Survey. For the Year 1922. Special Reports*, *10*, 1–63.

O'Donoghue, C. H. (1957). Some South African Bryozoa. *Transactions of the Royal Society of South Africa*, *35*, 71–95.

O'Donoghue, C. H., & de Watteville, D. (1939). The fishery grounds near Alexandria. XX. Bryozoa. *Fouad I Institute of Hydrobiology & Fisheries, Notes and Memoirs*, *34*, 1–58.

O'Donoghue, C. H., & de Watteville, D. (1944). Additional notes on Bryozoa from South Africa. *Annals of the Natal Museum*, *10*, 407–432.

Okamura, B. (1988). The influence of neighbors on the feeding of an epifaunal bryozoan. *Journal of Experimental Marine Biology and Ecology*, *120*(2), 105–123. https://doi.org/10.1016/0022-0981(88)90083-4

Oliver, J.-C., & Florence, W. K. (2016). A new species of *Taylorius* (Bryozoa: Escharinidae) from the east coast of South Africa. *African Natural History*, *12*, 1–4.

Orr, R. J. S., Waeschenbach, A., Enevoldsen, E. L. G., Boeve, J. P., Haugen, M. N., Voje, K. L., ... Liow, L. H. (2019). Bryozoan genera *Fenestrulina* and *Microporella* no longer confamilial; multi-gene phylogeny supports separation. *Zoological Journal of the Linnean Society*, *186*(1), 190–199. https://doi.org/10.1093/zoolinnean/zly055

Osburn, R. C. (1912). Bryozoa from Labrador, Newfoundland and Nova Scotia collected by Dr Owen Bryant. *United States National Museum Proceedings*, *43*, 275–289.

Osburn, R. C. (1923). Part D. Bryozoa. In *Report of the Canadian Arctic expedition, 1913-18* (Vol. 8). Ottawa: F. A. Acland.

Osburn, R. C. (1927). The Bryozoa of Curacao. *Bijdragen Tot de Dierkunde*, *25*, 123–132.

Osburn, R. C. (1933). Bryozoa of the Mount Desert Region. In W. Procter (Ed.), *Biological Survey of the Mount Desert region* (pp. 291–385). Philadelphia: Wistar Institute of Anatomy and Biology.

Osburn, R. C. (1936). Bryozoa collected in the American Arctic by Captain R.A. Bartlett. *Journal of the Washington Academy of Science*, *26*, 538–543.

Osburn, R. C. (1940). Bryozoa of Porto Rico with a résumé of the West Indian bryozoan fauna. *Scientific Survey of Porto Rico and the Virgin Islands*, *16*, 321–486.

Osburn, R. C. (1944). A survey of the Bryozoa of Chesapeake Bay. *Chesapeake Biological Laboratory Publication*, *63*, 3–55.

Osburn, R. C. (1947). Bryozoa of the Allan Hancock Atlantic Expedition, 1939. *Allan Hancock Atlantic Expedition Report*, *5*, 1–66.

Osburn, R. C. (1950). *Bryozoa of the Pacific coast of America, Part 1, Cheilostomata-Anasca.* Los Angeles: University of Southern California Press.

Ostrovsky, A. N. (2008). The parental care in cheilostome bryozoans: A historical review. In P. N. Wyse Jackson & M. E. Spencer Jones (Eds.), *Annals of Bryozoology 2: Aspects of the history of research on bryozoans* (Vol. 2, pp. 211–245). Dublin: International Bryozoology Association.

Ostrovsky, A. N., Cáceres-Chamizo, J. P., Vávra, N., & Berning, B. (2011). Bryozoa of the Red Sea: History and current state of research. In P. N. Wyse Jackson & M. E. Spencer Jones (Eds.), *Annals of Bryozoology 3: Aspects of the History of Research on Bryozoans* (pp. 67–97). Dublin: International Bryozoology Association.

Ostrovsky, A. N., Vávra, N., & Porter, J. S. (2008). Sexual reproduction in gymnolaemate Bryozoa: History and perspectives of the research. In P. N. Wyse Jackson & M. E. Spencer Jones (Eds.), *Annals of Bryozoology 2: Aspects of the history of research on bryozoans.* Dublin: International Bryozoology Association.

Pagès-Escolà, M., Hereu, B., Garrabou, J., Montero-Serra, I., Gori, A., Gómez-Gras, D., … Linares, C. (2018). Divergent responses to warming of two common co-occurring Mediterranean bryozoans. *Scientific Reports*, *8*(1). https://doi.org/10.1038/s41598-018-36094-9

Peters, L., Konig, G., Terlau, H., & Wright, A. (2002). Four new bromotryptamine derivatives from the marine bryozoan *Flustra foliacea*. *Journal of Natural Products*, *65*(11), 1633–1637.

Pizzaferri, C. (2006). A new subspecies of Cupuladria cavernosa Cadée from the Pliocene of western Emilia (N. Italy) (Bryozoa Gymnolaemata Cheilostomatida Cupuladriidae). *Quaderno Di Studi E Notizie Di Storia Naturale Della Romagna*, *22*, 39–51.

Porter, J. S., Nunn, J. D., Ryland, J. S., Minchin, D., & Jones, M. E. S. (2017). The status of non-native bryozoans on the north coast of Ireland. *Bioinvasions Records*, *6*(4), 321–330.

Porter, J. S., Spencer Jones, M. E., Kuklinski, P., & Rouse, S. (2015). First records of marine invasive non-native Bryozoa in Norwegian coastal waters from Bergen to Trondheim. *BioInvasions Records*, *4*(3), 157–169. https://doi.org/doi: http://dx.doi.org/10.3391/bir.2015.4.3.02

Powell, N. A. (1968). Bryozoa (Polyzoa) of Arctic Canada. *Journal of the Fisheries Research Board of Canada*, *25*, 2269–2320.

Powell, N. A. (1968). Studies on Bryozoa (Polyzoa) of the Bay of Fundy region. II. - Bryozoa from fifty fathoms, Bay of Fundy (I). *Cahiers de Biologie Marine*, *9*, 247–259.

Powell, N. A. (1969). A checklist of Indo-Pacific Bryozoa in the Red Sea. *Israel Journal of Zoology*, *18*, 357–362.

Powell, N. A., & Crowell, G. D. (1967). Studies on Bryozoa (Polyzoa) of the Bay of Fundy Region. 1. Bryozoa of the intertidal zone of the Minas Basin and Bay of Fundy. *Cahiers de Biologie Marine*, *8*, 331–347.

Quaiyum, S., Fortunato, H., Gonzaga, L., & Okabe, S. (2018). Antimicrobial Activity in the Marine Cheilostome Bryozoan *Cryptosula* zavjalovensis Kubanin, 1976. *Journal of Antimicrobial Agents*, *04*(03). https://doi.org/10.4172/2472-1212.1000178

Rabaoui, L., Tlig-Zouari, S., Cosentino, A., & Ben Hassine, O. K. (2009). Associated fauna of the fan shell *Pinna nobilis* (Mollusca: Bivalvia) in the northern and eastern Tunisian coasts. *Scientia Marina*, *73*(1), 129–141.

Ramalho, L. V., & Calliari, L. (2015). Bryozoans from Rio Grande do Sul Continental Shelf, Southern Brazil. *Zootaxa*, *3955*(4), 569–587.

Ramalho, L. V., López-Fé, C. M., & Rueda, J. L. (2018). Three species of *Reteporella* (Bryozoa: Cheilostomata) in a diapiric and mud volcano field of the Gulf of Cádiz, with the description of *Reteporella victori* n. Sp. *Zootaxa*, *4375*(1), 90. https://doi.org/10.11646/zootaxa.4375.1.4

Ramalho, L. V., Muricy, G., & Taylor, P. D. (2005). Taxonomy and distribution of *Bugula* (Bryozoa: Cheilostomata; Anasca) in Rio de Janeiro State, Brasil. In H. I. Moyano, J. M. Cancino, & P. N. Wyse Jackson (Eds.), *Bryozoan Studies 2004* (pp. 231–243). Leiden: Balkema.

Ramalho, L. V., Muricy, G., & Taylor, P. D. (2008). Taxonomy of *Beania* Johnston, 1840 (Bryozoa, Flustrina) from Arraial do Cabo, Rio de Janeiro State, Brazil. *Arquivos Do Museu Nacional, Rio de Janeiro*, *66*(3-4), 499–508.

Ramalho, L. V., Taylor, P. D., Moraes, F. C., Moura, R., Amado-Filho, G. M., & Bastos, A. C. (2018). Bryozoan framework composition in the oddly shaped reefs from Abrolhos Bank, Brazil, southwestern Atlantic: Taxonomy and ecology. *Zootaxa*, *4483*(1), 155–186. https://doi.org/10.11646/zootaxa.4483.1.6

Ramalho, L. V., Taylor, P. D., & Muricy, G. (2014). New records of *Catenicella* de Blainville, 1830 (Catenicellidae: Cheilostomata: Ascophora) in Rio de Janeiro State, Brazil. *Check List*, *10*(1), 170–174.

Rao, K., & Ganapati, P. N. (1978). Ecology of fouling bryozoans at Visakhapatnam Harbour. *Proceedings of the Indian Academy of Sciences Section B*, *87*(3), 63–75.

Reverter-Gil, O., Berning, B., & Souto, J. (2015a). Diversity and systematics of *Schizomavella* species (Bryozoa: Bitectiporidae) from the bathyal NE Atlantic. *PLoS ONE*, *10*(10), e0139084. https://doi.org/doi:10.1371/journal.pone.0139084

Reverter Gil, O., & Fernández Pulpeiro, E. (1999a). Some little-known species of Bryozoa described by J. Jullien. *Journal of Natural History*, *33*(9), 1403–1418.

Reverter-Gil, O., & Fernández-Pulpeiro, E. (1997). Two new species of *Schizomavella* (Bryozoa, Cheilostomatida). *Cahiers de Biologie Marine*, *38*, 1–6.

Reverter Gil, O., & Fernández Pulpeiro, E. (1999b). Some records of bryozoans from NW Spain. *Cahiers de Biologie Marine*, *40*(1), 35–45.

Reverter-Gil, O., & Souto, J. (2015). Redescription of some species of Bryozoa described by J. Jullien and L. Calvet in the NE Atlantic. *European Journal of Taxonomy*, *157*, 1–17.

Reverter-Gil, O., Souto, J., & Fernandez-Pulpeiro, E. (2011). Revision of the genus *Crepis* Jullien (Bryozoa: Cheilostomata) with description of a new genus and family and notes on Chlidoniidae. *Zootaxa*, *2993*, 1–22.

Reverter-Gil, O., Souto, J., & Fernández-Pulpeiro, E. (2012a). A new genus of Lanceoporidae (Bryozoa, Cheilostomata). *Zootaxa*, *3339*, 1–29.

Reverter-Gil, O., Souto, J., & Fernández Pulpeiro, E. (2009). Three new species of Iberian cheilostomate Bryozoa. *Journal of the Marine Biological Association of the United Kingdom*, *89*(7), 1499–1506. https://doi.org/10.1017/S0025315409000496

Reverter-Gil, O., Souto, J., & Fernández-Pulpeiro, E. (2012b). New and little known species of Bryozoa from Iberian Atlantic waters. *Zoosystema*, *34*(1), 157–170.

Reverter-Gil, O., Souto, J., Novosel, M., & Tilbrook, K. J. (2015b). Adriatic species of *Schizomavella* (Bryozoa: Cheilostomata). *Journal of Natural History*, *50*(5-6), 281–321.

Reverter-Gil, O., Souto, J., & Trigo, J. E. (2019). New species and new records of bryozoans from Galicia (NW Spain). *Journal of Natural History*, *53*(3-4), 221–251. https://doi.org/10.1080/00222933.2019.1582815

Ridley, S. O. (1881). Account of Polyzoa collected during the survey of H.M.S. 'Alert,' in the Straits of Magellan and on the Coast of Patagonia. *Proceedings of the Zoological Society of London*, 44–61.

Riisgard, H. U., & Manriquez, P. (1997). Filter-feeding in fifteen marine ectoprocts (Bryozoa): Particle capture and water pumping. *Marine Ecology Progress Series*, *154*, 223–239. https://doi.org/10.3354/meps154223

Robertson, A. (1905). Non-incrusting cheilostomatous Bryozoa of the west coast of North America. *University of California Publications in Zoology.*

Robertson, A. (1908). The incrusting chilostomatous Bryozoa of the west coast of North America. *University of California Publications in Zoology*, *4*(5), 253–344.

Rodgers, P. J., & Woollacott, R. M. (2006). Systematics, variation, and developmental instability: Analysis of spine patterns in ancestrulae of a common bryozoan. *Journal of Natural History*, *40*(21-22), 1351–1368. Retrieved from://000241266500006

Rogick, M. D. (1956). Bryozoa of the United States Navy's 1947-1948 Antarctic Expedition, I-IV. *Proceedings of the United States National Museum*, *105*(3358), 221–317.

Ross, J. R. P. (1974). Reef associated Ectoprocta from central region, Great Barrier Reef. *Proceedings of the Second International Coral Reef Symposium*, *1*, 349–352.

Rosso, A., Di Martino, E., Sanfilippo, R., & Di Martino, V. (2012). Bryozoan communities and thanatocoenoses from submarine caves in the Plemmirio Marine Protected Area (SE Sicily). In A. Ernst, P. Schäfer, & J. Scholz (Eds.), *Bryozoan Studies 2010* (Vol. 143, pp. 251–269). Berlin: Springer.

Rosso, A., Gerovasileiou, V., Sanfilippo, R., & Guido, A. (2019). Undisclosed bryodiversity of submarine caves of the Aegean Sea (Eastern Mediterranean). *Proceedings of the 2nd Mediterranean Symposium on the conservation of Dark Habitats*, 47–52. Antalya, Turkey.

Rosso, A., Sciuto, F., Sanfilippo, R., & Jones, M. S. (2017). The bryozoan genus *Arbocuspis* (Cheilostomata, Electridae) from the Indian Ocean, with description of a new species from off southwestern Thailand, Andaman Sea. *Zootaxa*, *4282*(1), 95–110.

Rosso, A., & Taylor, P. D. (2002). A new anascan cheilostome bryozoan from Icelandic deep waters and its uniserial colony growth pattern. *Sarsia*, *87*, 35–46.

Rouse, S., Spencer Jones, M. E., & Porter, J. S. (2013). Spatial and temporal patterns of bryozoan distribution and diversity in the Scottish sea regions. *Marine Ecology*, *35*(s1), 85–102.

Rörig, L. R., Ottonelli, M., Itokazu, A. G., Maraschin, M., Lins, J. V. H., Abreu, P. C. V., … Ramalho, L. V. (2017). Blooms of bryozoans and epibenthic diatoms in an urbanized sandy Beach (Balneário Camboriú-SC-Brazil): Dynamics, possible causes and biomass characterization. *Brazilian Journal of Oceanography*, *65*(4), 678–694.

Rubin, J. A. (1985). Mortality and avoidance of competitive overgrowth in encrusting Bryozoa. *Marine Ecology Progress Series*, *23*, 291–299.

Russ, G. R. (1982). Overgrowth in a marine epifaunal community: Competitive hierarchies and competitive networks. *Oecologia*, *53*, 12–19.

Ryland, J. S. (1969). A nomenclatural index to "A History of the British Marine Polyzoa" by T. Hincks (1880). *Bulletin of the British Museum (Natural History), Zoology*, *17*(6), 205–260.

Ryland, J. S. (1995). Bryozoa. In P. J. Hayward & J. S. Ryland (Eds.), *Handbook of the marine fauna of north-west Europe* (pp. 629–661). Oxford, New York: Oxford University Press.

Ryland, J. S., Bishop, J. D., De Blauwe, H., El Nagar, A., Minchin, D., Wood, C. A., & Yunnie, A. L. (2011). Alien species of *Bugula* (Bryozoa) along the Atlantic coasts of Europe. *Aquatic Invasions*, *6*(1), 17–31.

Ryland, J. S., & Gordon, D. P. (1977). Some New Zealand and British species of *Hippothoa* (Bryozoa: Cheilostomata). *Journal of the Royal Society of New Zealand*, *7*(1), 17–49.

Ryland, J. S., & Hayward, P. (1992). Bryozoa from Heron Island, Great Barrier Reef. *Memoirs of the Queensland Museum*, *32*(1), 223–301.

Ryland, J. S., Holt, R., Loxton, J., Spencer Jones, M. E., & Porter, J. S. (2014). First occurrence of the non-native bryozoan *Schizoporella japonica* Ortmann (1890) in Western Europe. *Zootaxa*, *3780*(3), 481–502.

Ryland, J. S., & Stebbing, A. R. D. (1971). Settlement and orientated growth in epiphytic and epizoic bryozoans. *Fourth European Marine Biology Symposium*, 105–124.

Ryland, J. S., & Stebbing, A. R. D. (1971). Two little known bryozoans from the west of Ireland. *The Irish Naturalist's Journal*, *17*(3), 65–70.

Santagata, S. (2015). Ectoprocta. In A. Wanninger (Ed.), *Evolutionary Developmental Biology of Invertebrates 2 Lophotrochozoa (Spiralia)* (pp. 247–262). Retrieved from http://link.springer.com/chapter/10.1007/978-3-7091-1871-9_11

Santagata, S., Ade, V., Mahon, A. R., Wisocki, P. A., & Halanych, K. M. (2018). Compositional differences in the habitat-forming bryozoan communities of the antarctic shelf. *Frontiers in Ecology and Evolution*, *6*. https://doi.org/10.3389/fevo.2018.00116

Santana, F. T., Ramalho, L. V., & Gumarães, C. P. (2009). A new species of *Metrarabdotos* (Bryozoa, Ascophora) from Brazil. *Zootaxa*, *2222*(57-65).

Schäfer, P. (2008). Diversity patterns of modern Arctic and Antarctic bryozoans. In H. Okada, S. F. Mawatari, N. Suzuki, & P. Gautam (Eds.), *Origin and Evolution of Natural Diversity* (pp. 57–66).

Schäfer, P., Fortunato, H., & Blaschek, H. (2014). Calcification patterns in the cheilostome bryozoan Flustra foliacea (L.) From the North Sea and Baltic Sea. *Studi Trentini Di Scienze Naturali*, *94*, 213–221.

Scholz, J., Mawatari, S. F., & Hirose, M. (2008). The Bryozoan diversity mystery: Why do we have about 1000 Species in Japanese waters? *Origin and Evolution of Natural Diversity : Proceedings of the International Symposium, the Origin and Evolution of Natural Diversity, Held from 1-5 October 2007 in Sapporo, Japan*, 129–135.

Schopf, T. J. (1973). Ergonomics of polymorphism: Its relation to the colony as the unit of natural selection in species of the phylum Ectoprocta. In R. Boardman, A. Cheetham, & W. Oliver (Eds.), *Animal Colonies: Development and Function Through Time* (pp. 247–294).

Schopf, T. J. M. (1969). Paleoecology of ectoprocts (bryozoans). *Journal of Paleontology*, *43*(2), 234–244.

Schopf, T. J. M. (1974). Ectoprocts as associates of coral reefs: St. Croix, U.S. Virgin Islands. *Proceedings of the Second International Reef Symposium*, *1*, 353–356. Brisbane: Great Barrier Reef Committee.

Sears, M. A. B., & Woollacott, R. M. (2008). Alice Robertson: Educator and marine zoologist. In P. N. Wyse Jackson & M. E. Spencer Jones (Eds.), *Annals of Bryozoology 2: Aspects of the history of research on bryozoans.* (pp. 305–345). Dublin: International Bryozoology Association.

Sears, M. A., & Woollacott, R. M. (2011). Reverend William F. Lynch: A Life in Science and Education. In *Annals of Bryozoology 3: Aspects of the history of research on bryozoans.*

Sears, M. A., & Woollacott, R. M. (2014). *Benjamin Harrison Grave: American Marine Invertebrate Zoologist.*

Sebastian, P., & Cumming, R. L. (2016). Three new species of *Calyptotheca* (Bryozoa: Lanceoporidae) from the great barrier reef, tropical Australia. *Zootaxa*, *4079*(4), 467–479.

Seo, J. E. (1994). Two species of Celleporaria (Cheilostomata: Bryozoa) from Korea. *Korean Journal of Systematic Zoology*, *10*(2), 189–197.

Seo, J. E. (1996). Two new species of Membraniporoidea (Bryozoa: Cheilostomata) from Korea. *Korean Journal of Systematic Zoology*, *12*(1), 45–51.

Seo, J. E., & Gong, Y. H. (2006). A new species and two new records of cheilostomata (Bryozoa) from Korea. *Korean Journal of Systematic Zoology*, *22*(1), 13–16.

Seo, J. E., & Min, B. S. (2009). A faunistic study on cheilostomatous bryozoans from the shoreline of South Korea, with two new species. *Korean Journal of Systematic Zoology*, *25*(1), 19–40.

Seroy, S., & Grünbaum, D. (2018). Individual and population level effects of ocean acidification on a predator-prey system with inducible defenses: Bryozoan-nudibranch interactions in the Salish Sea. *Marine Ecology Progress Series*, *607*, 1–18. https://doi.org/10.3354/meps12793

Sharp, K. H., Davidson, S. K., & Haygood, M. G. (2007). Localization of 'Candidatus Endobugula sertula'and the bryostatins throughout the life cycle of the bryozoan Bugula neritina. *The ISME Journal*, *1*(8), 693–702.

Shier, D. E. (1964). Marine Bryozoa from northwest Florida. *Bulletin of Marine Science*, *14*(4), 603–662.

Silén, L. (1950). On the Mobility of Entire Zoids in Bryozoa. *Acta Zoologica*, *31*(2-3), 349–386. https://doi.org/10.1111/j.1463-6395.1950.tb00515.x

Skinner, L. F. (2014). Foraminifera as predators on encrusting bryozoans. *Journal of Foraminiferal Research*, *44*(1), 58–61.

Smith, A. M. (2014). Growth and Calcification of Marine Bryozoans in a Changing Ocean. *The Biological Bulletin*, *226*(3), 203–210. Retrieved from http://www.biolbull.org/content/226/3/203

Smith, A. M., & Girvan, E. (2010). Understanding a bimineralic bryozoan: Skeletal structure and carbonate mineralogy of *Odontionella cyclops* (Foveolariidae: Cheilostomata: Bryozoa) in New Zealand. *Palaeogeography, Palaeoclimatology, Palaeoecology*, *289*, 113–122. https://doi.org/doi:10.1016/j.palaeo.2010.02.022

Smith, A. M., & Gordon, D. P. (2011). Bryozoans of southern New Zealand: A field identification guide. *New Zealand Aquatic Environment and Biodiversity Report*, *75*, 1–64.

Smith, A. M., & Lawton, E. I. (2010). Growing up in the temperate zone: Age, growth, calcification and carbonate mineralogy of Melicerita chathamensis (Bryozoa) in southern New Zealand. *Palaeogeography, Palaeoclimatology, Palaeoecology*, *298*, 271–277. https://doi.org/doi:10.1016/j.palaeo.2010.09.033

Smith, A. M., Spencer Jones, M., & Wyse Jackson, P. N. (2014). Bryozoans of the

Krusenstern Expedition (1803-1806). *International Bryozoology Association, Dublin*, 183–194.

Sokolover, N., Ostrovsky, A. N., & Ilan, M. (2018). *Schizoporella errata* (Bryozoa, Cheilostomata) in the Mediterranean Sea: Abundance, growth rate, and reproductive strategy. *Marine Biology Research*, *14*(8), 868–882. https://doi.org/10.1080/17451000.2018.1526385

Sokolover, N., Taylor, P. D., & Ilan, M. (2016). Bryozoa from the Mediterranean coast of Israel. *Mediterranean Marine Science*, *17*(2), 440–458. https://doi.org/DOI:http://dx.doi.org/10.12681/mms.1390

Sosa-Yañez, A., Fernández, L. H., & Olivera, Y. (2014). Primer registro de Adeonellopsis subsulcata (Smitt, 1873)(Bryozoa Gymnolaemata) en Cuba. *Revista Ciencias Marinas Y Costeras*, *6*(1), 29–36.

Sosa-Yañez, A., Vieira, L. M., & Solís-Marín, F. A. (2015). A new cheilostome bryozoan genus, *Abditoporella* (Hippoporidridae), from the eastern Pacific. *Zootaxa*, *3994*(2), 275–282.

Soule, D. F., & Soule, J. D. (1975). Species groups in Watersiporidae. In *Documents Des Laboratoires de Géologie de La Faculté Des Sciences de Lyon*: *Vol. 3. Bryozoa 1974* (pp. 299–309). Lyon: Université Claude Bernard.

Soule, J. D. (1959). Results of the Puritan-American Museum of Natural History Expedition to western Mexico. 6. Anascan Cheilostomata (Bryozoa) of the Gulf of California. *American Museum Novitates*, *1969*, 1–54.

Soule, J. D. (1961). Results of the Puritan-American Museum of Natural History Expedition to western Mexico. 13. Ascophoran Cheilostomata (Bryozoa) of the Gulf of California. *American Museum Novitates*, *2053*, 1–66.

Soule, J. D., Soule, D. F., & Chaney, H. W. (1998). Two new tropical Pacific species of *Cribralaria* (Bryozoa: Cribrilinidae) and a review of known species. *Irene Mcculloch Foundation Monograph Series*, *3*, 1–23.

Souto, J. (2019). Secondary homonymy in Bryozoa: The case of *Reteporella jullieni* (Cheilostomatida). *Zootaxa*, *4565*(2), 292. https://doi.org/10.11646/zootaxa.4565.2.13

Souto, J., Berning, B., & Ostrovsky, A. N. (2016). Systematics and diversity of deep-water Cheilostomata (Bryozoa) from Galicia Bank (NE Atlantic). *Zootaxa*, *4067*(4), 401–459.

Souto, J., Kaufmann, M. J., & Canning-Clode, J. (2015a). New species and new records of bryozoans from shallow waters of Madeira Island. *Zootaxa*, *3925*(4), 581–593.

Souto, J., Nascimento, K. B., Reverter-Gil, O., & Vieira, L. M. (2019). Dismantling the *Beania magellanica* (Busk, 1852) species complex (Bryozoa, Cheilostomata): Two new species from European waters. *Marine Biodiversity*, *49*(3), 1505–1518. https://doi.org/10.1007/s12526-018-0925-2

Souto, J., Ramalhosa, P., & Canning-Clode, J. (2015). Three non-indigenous species from Madeira harbors, including a new species of Parasmittina (Bryozoa). *Marine Biodiversity*.

Souto, J., & Reverter-Gil, O. (2019). Identity of bryozoan species described by Jullien & Calvet from the Bay of Biscay historically attributed to Smittia. *Zootaxa*, *4545*(1), 105. https://doi.org/10.11646/zootaxa.4545.1.6

Souto, J., Reverter-Gil, O., & de Blauwe, H. (2015b). New and little known species of *Celleporina* Gray, 1848 (Bryozoa, Cheilostomata) from the Atlantic–Mediterranean region. *Journal of the Marine Biological Association of the United Kingdom*, *95*(4), 723–734.

Souto, J., Reverter-Gil, O., De Blauwe, H., & Fernández Pulpeiro, E. (2014a). New records of bryozoans from Portugal. *Cahiers de Biologie Marine*, *55*, 129–150.

Souto, J., Reverter-Gil, O., & Fernandez-Pulpeiro, E. (2010). Bryozoa from detritic bottoms in the Menorca Channel (Balearic Islands, western Mediterranean), with notes on the genus Cribellopora. *Zootaxa*, *2536*(1), 36–52.

Souto, J., Reverter-Gil, O., & Fernández-Pulpeiro, E. (2010). Gymnolaemate bryozoans from the Algarve (southern Portugal): New species and biogeographical considerations. *Journal of the Marine Biological Association of the United Kingdom*, *90*, 1417–1439. Retrieved from http://dx.doi.org/10.1017/S0025315409991640

Souto, J., Reverter-Gil, O., & Fernández-Pulpeiro, E. (2011). Redescription of some bryozoan species originally described by J. Jullien from Iberian waters. *Zootaxa*, *2827*, 31–53.

Souto, J., Reverter-Gil, O., & Ostrovsky, A. N. (2014b). New species of Bryozoa from Madeira associated with rhodoliths. *Zootaxa*, *3795*(2), 135–151. https://doi.org/http://dx.doi.org/10.11646/zootaxa.3795.2.3

Stebbing, A. R. D. (1971). The epizoic fauna of *Flustra foliacea* (Bryozoa). *Journal of the Marine Biological Association of the United Kingdom*, *51*, 283–300.

Stricker, S. A. (1989). Settlement and metamorphosis of the marine bryozoan *Membranipora membranacea. Bulletin of Marine Science*, *45*(2), 387–405.

Ström, R. (1977). Brooding patterns of bryozoans. In R. M. Woollacott & R. L. Zimmer (Eds.), *Biology of bryozoans* (pp. 23–55). Academic Press.

Sutherland, J. P. (1981). The fouling community at Beaufort, North Carolina: A study in stability. *The American Naturalist*, *118*, 499–519.

Sutherland, J. P., & Karlson, R. H. (1977). Development and stability of the fouling community at Beaufort, North Carolina. *Ecological Monographs*, *47*, 425–446.

Suwa, T., Dick, M. H., & Mawatari, S. F. (1998). A new species of *Microporella* (Bryozoa, Cheilostomata) from Alaska. *Zoological Science*, *15*, 589–592.

Suwa, T., & Mawatari, S. F. (1998). Revision of seven species of *Microporella* (Bryozoa, Cheilostomatida) from Hokkaido, Japan, using new taxonomic characters. *Journal of Natural History*, *32*, 895–922.

Tadesse, M., Tabudravu, J. N., Jaspars, M., Strøm, M. B., Hansen, E., Andersen, J. H., … Haug, T. (2011). The antibacterial ent-eusynstyelamide B and eusynstyelamides D, E, and F from the Arctic bryozoan *Tegella cf. spitzbergensis. Journal of Natural Products*, *74*(4), 837–841.

Taylor, P. D., & Monks, N. (1997). A new cheilostome bryozoan genus pseudoplanktonic on molluscs and algae. *Invertebrate Biology*, *116*(1), 39–51.

Taylor, P. D., & Tan, S.-H. A. (2015). Cheilostome Bryozoa from Penang and Langkawi, Malaysia. *European Journal of Taxonomy*, *149*, 1–34.

Taylor, P. D., Tan S.-H., A., Kudryavstev, A. B., & Schopf, J. W. (2016). Carbonate mineralogy of a tropical bryozoan biota and its vulnerability to ocean acidification. *Marine Biology Research*, *12*(7), 776–780.

Teixidó, N., Garrabou, J., Gutt, J., & Arntz, W. E. (2004). Recovery in Antarctic benthos after iceberg disturbance: Trends in benthic composition, abundance and growth forms. *Marine Ecology Progress Series*, *278*, 1–16.

Temkin, M. H. (1994). Gamete spawning and fertilization in the Gymnolaemate Bryozoan *Membranipora membranacea. The Biological Bulletin*, *187*(2), 143–155.

Temkin, M. H. (1996). Comparative fertilization biology of gymnolaemate bryozoans. *Marine Biology*, *127*(2), 329–339.

Temkin, M. H., & Bortolami, S. B. (2004). Waveform dynamics of spermatozeugmata during the transfer from paternal to maternal individuals of *Membranipora membranacea. The Biological Bulletin*, *206*(1), 35–45.

Tian, X.-R., Tang, H.-F., Li, Y.-S., Lin, H.-W., Chen, X.-L., Ma, N., … Zhang, P.-H. (2011). New cytotoxic oxygenated sterols from the marine bryozoan *Cryptosula pallasiana. Marine Drugs*, *9*(2), 162–183.

Tilbrook, K. J. (1999). Description of Hippopodina feegeensis and three other species of Hippopodina Levinsen, 1909 (Bryozoa: Cheilostomatida). *Journal of Zoology, London*, *247*, 449–456.

Tilbrook, K. J. (2000). New bryozoan family from the Indo-Pacific shows unexpected diversity. *Journal of Marine Biological Association of the United Kingdom*, *80*, 1129–1130.

Tilbrook, K. J. (2011). New genus for a unique species of Indo-West Pacific bryozoan. *Zootaxa*, *3134*, 63–67.

Tilbrook, K. J. (2012a). Bryozoa, Cheilostomata: First records of two invasive species in Australia and the northerly range extension for a third. *Check List*, *8*(1), 181–183.

Tilbrook, K. J. (2012b). Review of the bryozoan genus Bryopesanser Tilbrook, 2006 (Escharinidae: Cheilostomata) with the description of 11 new species. *Zootaxa*, *3165*, 39–63.

Tilbrook, K. J., & Cook, P. L. (2005). Petraliellidae Harmer, 1957 (Bryozoa: Cheilostomata) from Queensland, Australia. *Systematics and Biodiversity*, *2*(3), 319–339.

Tilbrook, K. J., & Gordon, D. P. (2015). Bryozoa from the Straits of Johor, Singapore, with the description of a new species. *Raffles Bulletin of Zoology*, *31*, 255–263.

Tilbrook, K. J., & Gordon, D. P. (2016). Checklist of Singaporean Bryozoa and Entoprocta. *Raffles Bulletin of Zoology.*

Tilbrook, K. J., & Grischenko, A. V. (2004). New sub-Arctic species of the tropical genus *Antropora* (Bryozoa: Cheilostomata): A gastropod-pagurid crab associate. *Journal of the Marine Biological Association of the United Kingdom*, *84*, 1001–1004.

Tilbrook, K. J., Hayward, P. J., & Gordon, D. P. (2001). Cheilostomatous Bryozoa from Vanuatu. *Zoological Journal of the Linnean Society*, *131*, 35–109.

Tilbrook, K. J., & Vieira, L. M. (2012). *Scrupocellaria* (Bryozoa: Cheilostomata) from the Queensland coast, with the description of three new species. *Zootaxa*, *3528*(1), 29–48.

Tzioumis, V. (1994). Bryozoan stolonal outgrowths: A role in competitive interactions? *Journal of the Marine Biological Association of the United Kingdom*, *74*(1), 203–210.

Vaughan, D. G., Barnes, D. K., Fretwell, P. T., & Bingham, R. G. (2011). Potential seaways across west Antarctica. *Geochemistry, Geophysics, Geosystems*, *12*(10).

Venkatraman, C., Rajan, R., Louis, S., Shrinivaasu, S., & Padmanaban, P. (2016). Bryozoans of Gulf of Mannar Marine Biosphere Reserve, Southeast Coast of India. *Records of the Zoological Survey of India*, *116*(2), 167–189.

Vieira, L. M., Almeida, A. C. S., & Winston, J. E. (2016). Taxonomy of intertidal cheilostome Bryozoa of Maceió, northeastern Brazil. Part 1: Suborders Inovicellina, Malacostegina and Thalamoporellina. *Zootaxa*, *4097*(1), 59–83. https://doi.org/http://doi.org/10.11646/zootaxa.4097.1.3

Vieira, L. M., Farrapeira, C. M. R., Amaral, F. D., & Lira, S. M. A. (2012). Bryozoan biodiversity in Saint Peter and Saint Paul Archipelago, Brazil. *Cahiers de Biologie Marine*, *53*, 159–167.

Vieira, L. M., & Gordon, D. P. (2010). Eutaleola, a replacement name for the homonym Euteleia (Bryozoa: Pasytheidae). *Zoologia*, *27*(4), 646–648.

Vieira, L. M., Gordon, D. P., & Correia, M. D. (2007). First record of a living ditaxiporine catenicellid in the Atlantic, with a description of *Vasignyella ovicellata* n. Sp. (Bryozoa). *Zootaxa*, *1582*, 49–58.

Vieira, L. M., Gordon, D. P., Souza, F. B. C., & Haddad, M. A. (2010a). New and little-known cheilostomatous Bryozoa from the south and southeastern Brazilian continental shelf and slope. *Zootaxa*, *2722*, 1–53.

Vieira, L. M., & Migotto, A. E. (2015). *Membraniporopsis tubigera* (Osburn, 1940) (Bryozoa) on floating substrata: Evidence of a dispersal mechanism in the western Atlantic. *Marine Biodiversity*, *45*(2), 155–156.

Vieira, L. M., Migotto, A. E., & Winston, J. E. (2008). Synopsis and annotated checklist of Recent marine Bryozoa from Brazil. *Zootaxa*, *1810*, 1–39.

Vieira, L. M., Migotto, A. E., & Winston, J. E. (2010b). Marcusadorea, a new genus of lepralioid bryozoan from warm waters. *Zootaxa*, *2348*, 57–68.

Vieira, L. M., Migotto, A. E., & Winston, J. E. (2010c). Shallow-water species of *Beania* Johnston, 1840 (Bryozoa, Cheilostomata) from the tropical and subtropical Western Atlantic. *Zootaxa*, *2550*, 1–20.

Vieira, L. M., & Spencer Jones, M. E. (2012). The identity of Sertularia reptans Linnaeus, 1758 (Bryozoa, Candidae). *Zootaxa*, *3563*, 26–42.

Vieira, L. M., Spencer Jones, M. E., & Taylor, P. D. (2014a). The identity of the invasive fouling bryozoan Watersipora subtorquata (d'Orbigny) and some other congeneric species. *Zootaxa*, *3857*(2), 151–182.

Vieira, L. M., Spencer Jones, M. E., & Winston, J. E. (2013a). Cradoscrupocellaria, a new bryozoan genus for Scrupocellaria bertholletii (Audouin) and related species (Cheilostomata, Candidae): Taxonomy, biodiversity and distribution. *Zootaxa*, *3707*(1), 1–63. https://doi.org/http://dx.doi.org/10.11646/zootaxa.3707.1.1

Vieira, L. M., Spencer Jones, M. E., & Winston, J. E. (2013b). Resurrection of the genus *Licornia* for *Scrupocellaria jolloisii* (Bryozoa) and related species, with documentation of *L. jolloisii* as a non-indigenous species in the western Atlantic. *Journal of the Marine Biological Association of the United Kingdom*, *93*(7), 1911–1921. https://doi.org/10.1017/S0025315413000301

Vieira, L. M., Spencer Jones, M. E., Winston, J. E., Migotto, A. E., & Marques, A. C. (2014b). Evidence for polyphyly of the genus *Scrupocellaria* (Bryozoa: Candidae) based on a phylogenetic analysis of morphological characters. *PLoS ONE*, *9*(4), e95296. https://doi.org/10.1371/journal.pone.0095296

Vieira, L. M., & Stampar, S. N. (2014). A new *Fenestrulina* (Bryozoa, Cheilostomata) commensal with tube-dwelling anemones (Cnidaria, Ceriantharia) in the tropical southwestern Atlantic. *Zootaxa*, *3780*(2), 365–374.

Vieira, L. M., Winston, J. E., & Fehlauer-Ale, K. H. (2012). Nine New Species of *Bugula* Oken (Bryozoa: Cheilostomata) in Brazilian Shallow Waters. *PLoS ONE*, *7*(7), e40492. https://doi.org/10.1371/journal.pone.0040492

Vigeland, I. (1971). Port Phillip Bay Survey 2. Bryozoa. *Memoirs of the National Museum of Victoria*, *32*, 65–81.

Voronkov, A., Hop, H., & Gulliksen, B. (2016). Zoobenthic communities on hard-bottom habitats in Kongsfjorden, Svalbard. *Polar Biology*, *39*(11), 2077–2095.

Waeschenbach, A., Porter, J. S., & Hughes, R. N. (2012). Molecular variability in the *Celleporella hyalina* (Bryozoa; Cheilostomata) species complex: Evidence for cryptic speciation from complete mitochondrial genomes. *Molecular Biology Reports*, *39*, 8601–8614. https://doi.org/10.1007/s11033-012-1714-9

Walters, L. J., & Wethey, D. S. (1986). Surface topography influences competitive hierarchies on marine hard substrata: A field experiment. *Biological Bulletin*, *170*(3), 441–449.

Wang, H., Zhang, H., Wong, Y. H., Voolstra, C., Ravasi, T., B Bajic, V., & Qian, P. Y. (2010). Rapid transcriptome and proteome profiling of a non-model marine invertebrate, Bugula neritina. *Proteomics*, *10*(16), 2972–2981.

Wanninger, A., Koop, D., & Degnan, B. M. (2005). Immunocytochemistry and metamorphic fate of the larval nervous system of Triphyllozoon mucronatum (Ectoprocta: Gymnolaemata: Cheilostomata). *Zoomorphology*, *124*(4), 161–170. https://doi.org/10.1007/s00435-005-0004-7

Ward, M. A., & Thorpe, J. P. (1991). Distribution of encrusting bryozoans and other epifauna on the subtidal bivalve *Chlamys opercularis*. *Marine Biology*, *110*, 253–259.

Wasson, B., & De Blauwe, H. (2014). Two new records of cheilostome Bryozoa from British waters. *Marine Biodiversity Records*, *7*.

Waters, A. W. (1879). On the Bryozoa (Polyzoa) of the Bay of Naples. Cheilostomata (continued). *Annals and Magazine of Natural History, Ser. 5*, *3*, 114–126.

Waters, A. W. (1883). On fossil chilostomatous Bryozoa from Muddy Creek, Victoria. *Quarterly Journal of the Geological Society of London*, *39*, 423–443.

Waters, A. W. (1897). Notes on Bryozoa from Rapallo and other Mediterranean localities. - Chiefly Cellulariidae. *Journal of the Linnean Society of London, Zoology*, *26*, 1–23.

Waters, A. W. (1905). Bryozoa from near Cape Horn. *Journal of the Linnean Society (Zoology)*, *29*, 230–251.

Waters, A. W. (1906). Bryozoa from Chatham Island and d'Urville Island, New Zealand, collected by Professor H.Schauinsland. *Annals and Magazine of Natural History*, *(7)17*, 12–23.

Waters, A. W. (1913). The marine fauna of British East Africa and Zanzibar, from collections made by Cyril Crossland, M.A., B.Sc., F.Z.S., In the years 1901-1902. Bryozoa - Cheilostomata. *Proceedings of the Zoological Society of London*, *1913*, 458–537.

Waters, A. W. (1918). Some collections of the littoral marine fauna of the Cape Verde Islands, made by Cyril Crossland, M.A., B.Sc., F.Z.S., In the summer of 1904. - Bryozoa. *Journal of the Linnean Society (Zoology)*, *34*, 1–45.

Waters, A. W. (1925). Some cheilostomatous Bryozoa from Oran (Algiers). *Annals and Magazine of Natural History*, *9*(15), 651–661.

Watts, P. C., & Thorpe, J. P. (2006). Influence of contrasting larval developmental types upon the population-genetic structure of cheilostome bryozoans. *Marine Biology*, *149*, 1093–1101.

Wejnert, K. E., & Smith, A. M. (2008). Within-colony variation in skeletal mineralogy of *Adeonellopsis* sp. (Cheilostomata: Bryozoa) from New Zealand. *New Zealand Journal of Marine and Freshwater Research*, *42*, 389–395.

Wendt, D. E. (1996). Effect of larval swimming duration on success of metamorphosis and size of the ancestrular lophophore in *Bugula neritina* (Bryozoa). *Biological Bulletin*, *191*, 224–233.

Wendt, D. E., & Woollacott, R. M. (1999). Ontogenies of phototactic behavior and metamorphic competence in larvae of three species of *Bugula* (Bryozoa). *Invertebrate Biology*, *118*(1), 75–84. https://doi.org/10.2307/3226915

Wicksten, M.-K. a. (1996). Anthozoans, bryozoans, brachiopods, and tunicates of Rocas Alijos. *Monographiae Biologicae*, *75*, 277–283.

Wieczorek, S. K., & Todd, C. D. (1997). Inhibition and facilitation of bryozoan and ascidian settlement by natural multi-species biofilms: Effects of film age and the roles of active and passive larval attachment. *Marine Biology*, *128*, 463–473.

Winston, J. E. (1984). Shallow-water bryozoans of Carrie Bow Cay, Belize. *American Museum Novitates*, *2799*, 1–38.

Winston, J. E. (1984). Why bryozoans have avicularia: A review of the evidence. *American Museum Novitates*, (2789), 1–26.

Winston, J. E. (2004). Bryozoans from Belize. *Atoll Research Bulletin*, *523*, 1–14.

Winston, J. E. (2009). Stability and change in the Indian River area bryozoan fauna over a twenty-four year period. *Smithsonian Contributions to the Marine Sciences*, *38*, 229–239.

Winston, J. E. (2016). Bryozoa of Floridan Oculina reefs. *Zootaxa*, *4071*(1), 1–81.

Winston, J. E., & Eiseman, N. J. (1980). Bryozoan-algal associations in coastal and continental shelf waters of eastern Florida. *Florida Scientist*, *43*(2), 65–74.

Winston, J. E., & Hakansson, E. (1986). The interstitial bryozoan fauna from Capron Shoal, Florida. *American Museum Novitates*, *2865*, 1–50.

Winston, J. E., & Hayward, P. J. (2011). Bryozoans of the Northeast Coast of the United States: Taxonomic history and summary of a new survey. In P. N. Wyse Jackson & M. E. Spencer Jones (Eds.), *Annals of Bryozoology 3: Aspects of the history of research on bryozoans* (pp. 201–217). Dublin: International Bryozoology Association.

Winston, J. E., & Heimberg, B. F. (1986). Bryozoans from Bali, Lombok, and Komodo. *American Museum Novitates*, *2847*, 1–49.

Winston, J. E., & Maturo, F. J. S. (2009). Bryozoans (Ectoprocta) of the Gulf of Mexico. In D. L. Felder & D. K. Camp (Eds.), *Gulf of Mexico Origin, Waters, and Biota. Volume I, Biodiversity* (pp. 1147–1164). College Station: Texas A&M University Press.

Winston, J. E., & Vieira, L. M. (2013). Systematics of interstitial encrusting bryozoans from southeastern Brazil. *Zootaxa*, *3710*(2), 101–146.

Winston, J. E., Vieira, L. M., & Woollacott, R. M. (2014). Scientific results of the Hassler Expedition. Bryozoa. No. 2. Brazil. *Bulletin of the Museum of Comparative Zoology*, *161*(5), 139–239.

Winston, J. E., & Woollacott, R. M. (2008). Redescription and revision of some red-pigmented *Bugula* species. *Bulletin of the Museum of Comparative Zoology*, *159*(3), 179–212. https://doi.org/10.3099/0027-4100(2008)159[179:RAROSR]2.0.CO;2

Wong, Y. H., Arellano, S. M., Zhang, H., Ravasi, T., & Qian, P.-Y. (2010). Dependency on de novo protein synthesis and proteomic changes during metamorphosis of the marine bryozoan *Bugula neritina*. *Proteome Science*, *8*(1), 25. https://doi.org/10.1186/1477-5956-8-25

Wood, A. C. L., & Probert, P. K. (2013). Bryozoan-dominated benthos of Otago shelf, New Zealand: Its associated fauna, environmental setting and anthropogenic threats. *Journal of the Royal Society of New Zealand*, *43*(4), 231–249.

Wood, A. C. L., Rowden, A. A., Compton, T. J., Gordon, D. P., & Probert, P. K. (2013). Habitat-forming bryozoans in New Zealand: Their known and predicted distribution in relation to broad-scale environmental variables and fishing effort. *PLoS ONE*, *8*(9), e75160.

Woollacott, R. M., & Zimmer, R. L. (1972a). Fine structure of a potential photoreceptor organ in the larva of *Bugula neritina* (Bryozoa). *Zeitschrift Für Zellforschung Und Mikroskopische Anatomie*, *123*(4), 458–469.

Woollacott, R. M., & Zimmer, R. L. (1972b). Origin and structure of the brood chamber in *Bugula neritina* (Bryozoa). *Marine Biology*, *16*(2), 165–170. https://doi.org/10.1007/BF00347954

Xing, J., & Qian, P. Y. (1999). Tower cells of the marine bryozoan *Membranipora membranacea*. *Journal of Morphology*, *239*(2), 121–130.

Yang, H. J., Seo, J. E., & Gordon, D. P. (2018a). Sixteen new generic records of Korean Bryozoa from southern coastal waters and Jeju Island, East China Sea: Evidence of tropical affinities. *Zootaxa*, *4422*(4), 493–518. https://doi.org/10.11646/zootaxa.4422.4.3

Yang, H. J., Seo, J. E., Min, B. S., Grischenko, A. V., & Gordon, D. P. (2018b). Cribrilinidae (Bryozoa: Cheilostomata) of Korea. *Zootaxa*, *4377*(2), 216–234. https://doi.org/10.11646/zootaxa.4377.2.4

Yorke, A. F., & Metaxas, A. (2011). Interactions between an invasive and a native bryozoan (*Membranipora membranacea* and *Electra pilosa*) species on kelp and *Fucus* substrates in Nova Scotia, Canada. *Marine Biology*, *158*(10), 2299.

Zabala, M., & Maluquer, P. (1988). Illustrated keys for the classification of Mediterranean Bryozoa. *Treballs Del Museu de Zoologica*, *4*, 1–294.

Zabala i Limousin, M., Maluquer, P., & Harnelin, J. G. (1993). Epibiotic bryozoans on deep-water seleractinian corals from the Catalonia slope (western Mediterranean, Spain, France). *Scientia Marina*, *57*(1), 65–78.

Zabin, C. J., Marraffini, M., Lonhart, S. I., McCann, L., Ceballos, L., King, C., … Ruiz, G. M. (2018). Non-native species colonization of highly diverse, wave swept outer coast habitats in Central California. *Marine Biology*, *165*(2), 31. https://doi.org/10.1007/s00227-018-3284-4

Zabin, C. J., Obernolte, R., Mackie, J. A., Gentry, J., Harris, L., & Geller, J. B. (2010). A non-native bryozoan creates novel substrate on the mudflats in San Francisco Bay. *Marine Ecology Progress Series*, *412*, 129–139.

Zágoršek, K. (1994). Late Eocene (Priabonian) Cheilostomata (Bryozoa) from the Liptov Basin - Western Carpathians (Slovakia). *Neues Jahrbuch Für Geologie Und Paläontologie Monatshefte*, *193*(3), 361–382.

Zágoršek, K., & Vávra, N. (2007). Bryozoan fauna from Steinebrunn (Lower Austria, Badenian) - a revision to establish a basis for comparisons with Moravian faunas. *Scripta Facultatis Scientiarum Naturalium Universitatis Masaryk Brunensis*, *36*, 65–72.

Zenetos, A., Gofas, S., Morri, C., Rosso, A., Violanti, D., García Raso, J. E., … Azzurro, E. (2012). *Alien species in the Mediterranean Sea by 2012. A contribution to the application of European Union's Marine Strategy Framework Directive (MSFD). Part 2. Introduction trends and pathways.*

Zhang, H., Wong, Y. H., Wang, H., Chen, Z., Arellano, S. M., Ravasi, T., & Qian, P. Y. (2011). Quantitative proteomics identify molecular targets that are crucial in larval settlement and metamorphosis of *Bugula neritina*. *Journal of Proteome Research*, *10*(1).

Zimmer, R. L., & Woollacott, R. M. (1989). Larval morphology of the bryozoan *Watersipora arcuata* (Cheilostomata: Ascophora). *Journal of Morphology*, *199*(2), 125–150. Retrieved from http://onlinelibrary.wiley.com/doi/10.1002/jmor.1051990202/abstract

Zongguo, H. (2001). Bryozoa [Ectoprocta]. In *Marine Species and Their Distribution in China's Seas* (pp. 369–383). Malabar: Krieger Publishing Company.
